# Recreational fisheries as a driver of salmonid population conservation

**DOI:** 10.1101/642660

**Authors:** Thomas A. Worthington, Ian Worthington, Ian P. Vaughan, Steve J Ormerod, Isabelle Durance

## Abstract

1. The need to monitor and protect biodiversity has never been greater, yet resources are often constrained economically. The ecosystem service paradigm could promote nature conservation while sustaining economic activity and other societal benefits, but most efforts to assess biodiversity-ecosystem service (B-ES) links have focused on diversity measures, with little attention on how species abundance relates to the magnitude of ES provision.
2. Here, we utilised four national scale, multi-decadal, Atlantic salmon (*Salmo salar*) datasets to investigate links between juvenile density, the abundance of returning adults, and two measures of recreational angling provision: rod catches and angling effort.
3. Recreational rod catches only tracked juvenile density and returning adult numbers in catchments where juvenile and adult numbers were decreasing, implying important early-warning of ES decline. In contrast, angling effort declined consistently through time.
4. *Synthesis and applications*. These data illustrate i) the difficulty in measuring ES in ways that explicitly relate human resource use to nature conservation, and ii) the need for better quantification of populations at all life stages that determine ES provision, particularly in species where long-distance movements bring exposure to multiple global pressures. We suggest additional opportunities (e.g., monitoring of smolts, eDNA and citizen science initiatives) to facilitate conservation efforts and increase capacity to monitor ecosystem service sustainability.

## INTRODUCTION

Over the last decade, the ecosystem services (ES) concept has gained considerable attention as a framework that could reconcile the needs of biodiversity conservation with economic growth and societal benefits derived from natural resources (Fisher, Turner, & Morling, 2009; Mekonnen & Hoekstra, 2011). However, despite increasing concerns over the persistent declines in the abundances of many species (WWF, 2018), investigations on the links between biodiversity and ecosystem services (B-ES links) have mostly focused on ‘richness’ based biodiversity indicators (Cardinale et al., 2012). While population abundance is a recognised key indicator of biodiversity (Díaz, Fargione, Chaplin III, & Tilman, 2006), little attention has focused on the implications of population decline on ES provision (Gaston et al., 2018).

Positive links between biodiverse ecosystems and yields of food or fibre with high market value are documented (Luck et al., 2009), but quantitative evidence linking biodiversity to less tangible ES is still scarce (Durance et al., 2016). This is particularly true for the non-material benefits (cultural ecosystem services (CES)) that people obtain from ecosystems through spiritual enrichment, cognitive development, reflection, recreation, or aesthetic experiences (Small, Munday, & Durance, 2017). And yet people appreciate, benefit and gain wellbeing from CES directly as an implicit link with the natural environment (Daniel et al., 2012; Schaich, Biding, & Plieninger, 2010) which should provide an impetus for greater conservation efforts (Angulo-Valdés & Hatcher, 2010; Plieninger, Dijks, Oteros-Rozas, & Bieling, 2013).

Concerns over the decline of salmonid fish such as Atlantic salmon (*Salmo salar*) provide a prime example where the ES paradigm might benefit both people and ecosystems through both tangible and CES values. Atlantic salmon are an important food source (albeit largely derived from aquaculture (Parrish, Behnke, Gephard, McCormick, & Reeves, 1998)), but wild fish also support a lucrative recreational angling industry with clear economic values (Butler, Radford, Riddington, & Laughton, 2009; Kennedy & Crozier, 1997). They also have a range of less tangible benefits to society including raising awareness of environmental issues (Cowx & Portocarrero Aya, 2011) or aesthetic value (Holmlund & Hammer, 2004; O’Reilly & Mawle, 2006). The value of Atlantic salmon is particularly pertinent in rural areas where the influx of visitors supports jobs and contributes significantly to local economies (Aprahamian, Hickley, Sheilds, & Mawle, 2010; Peirson, Tingley, Spurgeon, & Radford, 2001).

There are many ways to assess the value of an ES, but the real challenge lies in capturing both the social and ecological dimensions of the service. In the case of angling, the quality of this recreational ES results both from capturing a fish, but also in more intangible characteristics such as the angling experience (Arlinghaus, 2006; Ringold, Boyd, Landers, & Weber, 2013). These social dimensions of the service are notoriously difficult and resource intensive to account for (Chan, Guerry, et al., 2012), as is the case for most CES (see Small et al., 2017 for an overview). As a result, ES assessments are often limited to the use of proxy data (Eigenbrod et al., 2010; Naidoo et al., 2008), and in the case of Atlantic Salmon, adult rod catches and angling effort are the measures most often used to express the quality of the angling service.

Most conservation efforts for Atlantic Salmon have focused on juvenile monitoring programmes because the results of these surveys provide a cost-effective measure of the population. However, the species has a complex and very variable life history involving freshwater and marine migratory stages of variable duration (Hutchings & Jones, 1998; Klemetsen et al., 2003, Supporting Information Appendix S1). As a consequence, juvenile life stages are separated from the returning adult stages by the ocean migration phase where the mechanisms affecting survival are less well understood (Friedland, 1998; Friedland et al., 2009). The challenge thus lies in providing the mechanistic link between population size at one life history stage (e.g., juveniles) and service provision at another. It therefore seems unlikely that there are simple connections between the current monitoring of Atlantic salmon populations through juveniles and the ES provided by adults.

The high profile of Atlantic salmon means that significant amounts of data have been collected on both their biology and exploitation. The financial and resource costs of acquiring such data has been substantial (Hering et al., 2010; Kallis & Butler, 2001), and this gives impetus to the need to demonstrate clear links between expenditure on monitoring and the social benefits returned. These data are relevant to different steps of the ES ‘cascade’ which depicts how biodiversity drives the availability of ES and the benefits and value society derive from it (Haines-Young & Potschin, 2010; Small et al., 2017). In the case of Atlantic salmon, the different steps of the ES cascade include: juvenile density as a measure of biodiversity, returning adult numbers as a measure of potential ES use, and ultimately recreational rod catches or angling effort as a measure of actual ES use (Rows 2 and 3 in Fig. 1; Small et al., 2017).

**Fig. 1:**
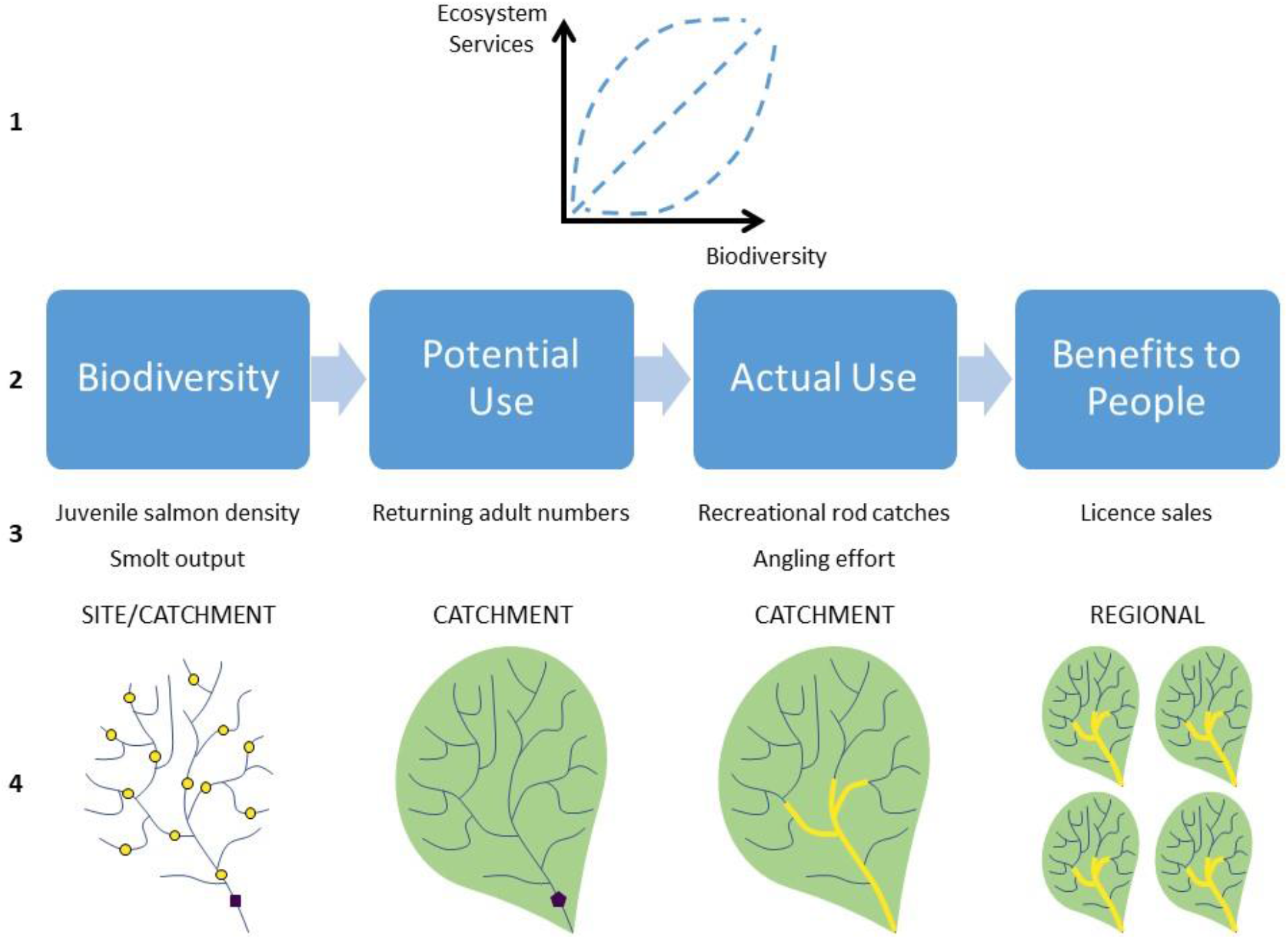
The relationship between spatial scale of the available data on Atlantic salmon populations and the levels of the ecosystem service cascade (Small et al., 2017). The theoretical location of juvenile density monitoring sites (yellow circles), smolt monitoring (purple circle), adult trap (yellow pentagon) and angling reaches (yellow lines) within the catchment.

In Europe, licencing schemes often record actual ES use (rod catch and angling effort), and are part of broader efforts aimed at supporting the conservation of salmon which stocks have been recognised as in decline (NASCO, 2016; World Conservation Monitoring Centre, 1996; WWF, 2001). It is important to note however, that the available data vary in terms of the scale of the investigation (Row 4 in Fig. 1) as well as the temporal and spatial coverage of the data (CEFAS, Environment Agency, & Natural Resources Wales, 2015a; Cowx & Fraser, 2003), with juvenile population abundance monitored at a local scale, returning adult numbers mostly sporadically at a single point in the catchment, and rod catches aggregated at administrative level. Yet, what the ES cascade framework highlights, is that to be meaningful, assessments of the quality of the ES need to reflect not just the immediate and proximal quality (e.g. rod catch), but also the resilience of the service in the short term (e.g. returning adults) or the long term (e.g. juveniles).

Here we analyse and compare four national scale and multi-decadal Atlantic salmon datasets: *juvenile density* as a measure of Atlantic salmon population biodiversity, *returning adult numbers* as a measure of potential ES use, and *rod catches* and *angling effort* as measures of actual ES use. Given the complexity of the Atlantic salmon’s life history and the considerable distances moved between life stages, our working hypothesis is that trends in *rod catches* and *angling effort* for adult fish are unlikely to track *juvenile density*, rendering current monitoring poorly suited to capture ecosystem service value.

## METHODS

### Study area

To provide a framework for the analysis, we identified catchments (*n* = 141, split into eight regions; Fig. 2) that support populations of Atlantic salmon from the UK National Fish Population Database (NFPD). These catchments covered most of England and Wales and ranged in size from Black Beck (12.5 km^2^) to the River Severn (10082.5 km^2^, median = 150.0 km^2^).

**Fig. 2:**
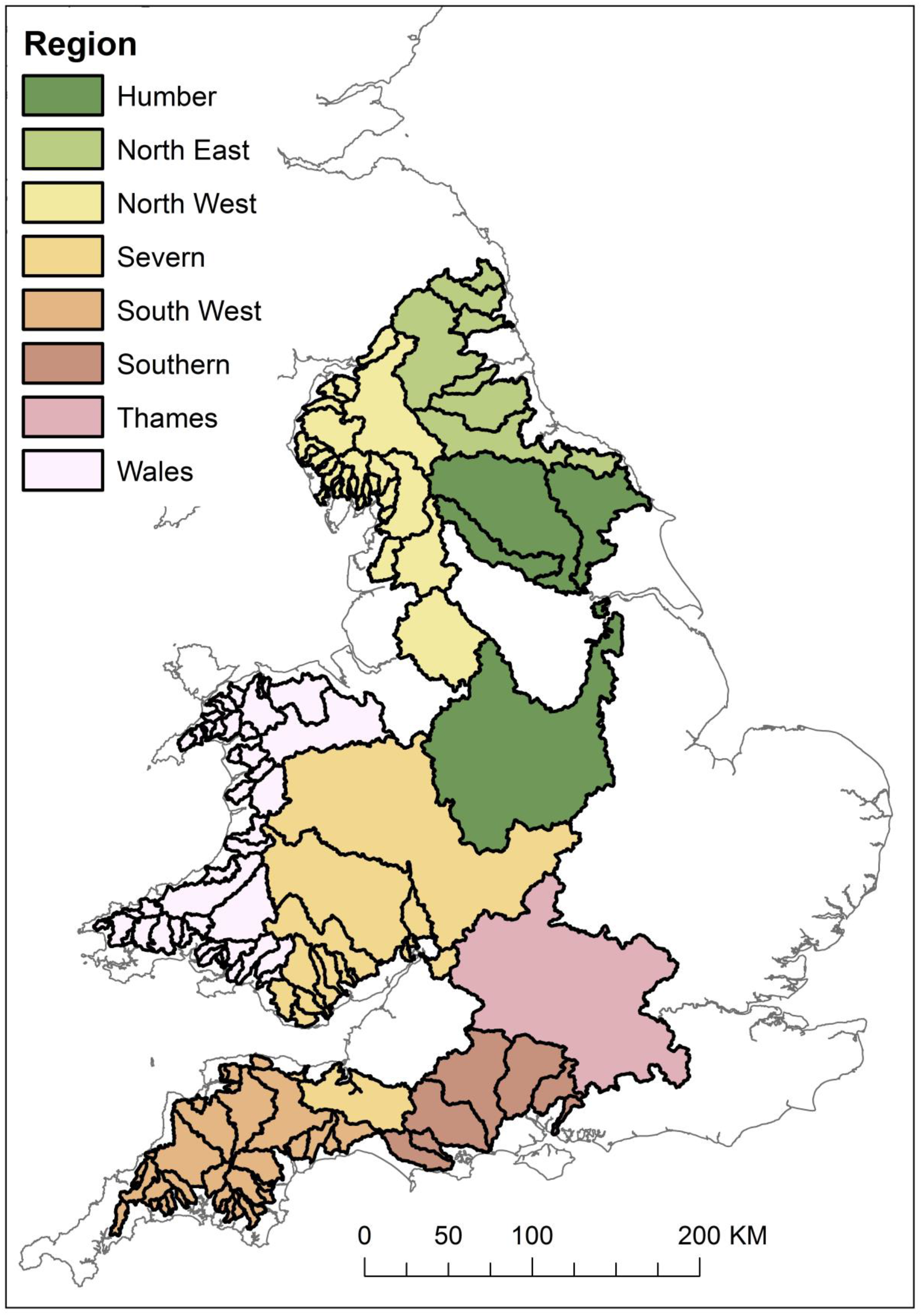
Location of the 141 salmon catchments and the eight regions.

### Juvenile density - a traditional ‘biodiversity’ measure for population monitoring

Juvenile density data were available for England and Wales between 1975 and 2018, with 21660 individual surveys spanning more than four decades. We retained Atlantic salmon data that were collected using electric fishing with semi-quantitative or quantitative methods, with the number of juveniles recorded during the first run converted to a density by dividing by the area surveyed. Within each salmon catchment, sites which had 10 or more years of data, with at least 6 years where salmon were recorded, were identified. This resulted in a final dataset of 7526 surveys at 501 locations used for the analysis. To examine temporal trends in juvenile Atlantic salmon density, we fitted a generalized additive mixed model (GAMM) to each of the eight regions, regressing density upon sampling year. For each region (except those regions containing a single salmon catchment) a model that allowed the non-linear relationship between density and time to take a different shape across individual salmon catchments was used (Supporting Information Table S1). Each model included a random effect for sampling site. The structure of the models allowed trends in juvenile salmon to be assessed at the catchment level. These trends were constructed using a variable number of sites (range 1 – 38 sites per catchment), with sampling at each site not consistent across years.

### Returning adult numbers – a measure of potential ES use

Locations with ≥10 years of data on the number of returning adult Atlantic salmon collected using electronic fish counters or traps (CEFAS & Environment Agency, 2000, 2002, 2005; CEFAS, Environment Agency, & Natural Resources Wales, 2015b, 2018), were only available from 14 out of the 141 salmon catchments. Temporal coverage at these locations ranged between 12 and 30 years. We analysed trends in returning adult fish with two generalized additive models (GAMs), with one GAM for those rivers where mean annual number of returning adults was <2500 and the other for rivers where mean annual number of returning adults was >2500 (Supporting Information Table S1). The model allowed the shape of the relationship between total catch and year to vary between salmon catchments.

### Rod catches and angling effort – measures of actual ES use

To examine trends in actual Atlantic salmon ES use, we assessed variation in recreational angling, both in terms of the number of fish caught (adult rod catches) and amount of time spent angling (angling effort). We collated data on the numbers of rod-caught Atlantic salmon recorded from angling licence returns for the ‘principal’ salmon rivers across England and Wales, and angling effort in terms of the reported number of days fished for salmon and migratory trout per license return (for a subset of the licence returns) from the Environment Agency’s Salmonid and Freshwater Fisheries Statistics for England and Wales (1994 – 2017) reports. Data on rod catches were available for 74 catchments and covered the period 1994 – 2017 for all but four catchments. Data on angling effort was available for 73 catchments, all covering the period 1994 – 2017. It should be noted that angling effort data recorded for the licence returns was for both salmon and sea trout and provides an indication of changes in angling behaviour.

We analysed trends in adult rod catches using generalized additive models (GAMs). A separate negative binomial GAM was fitted to the total rod catch for each region. As before the model allowed the non-linear relationship between total catch and time to take a different shape across individual salmon catchments. The final set of models assessed trend in angling effort, measured as catch per licence day. As previously, an individual Gaussian GAM was fitted to each region, with the model structure allowing a different non-linear trend in each catchment.

### Linking biodiversity to benefit along the ES cascade

The trends in the four core datasets (juvenile density, returning adults numbers, rod catches and angling effort) were visually compared for each catchment. Extended methods that detail data selection procedures and analyses are available in the Supporting Information Appendix S2.

## RESULTS

### Juvenile Atlantic salmon densities present non-uniform trends

Of the potential 141 salmon catchments across the eight regions of England and Wales (Fig. 2), 83 had sufficiently long data runs (>10 years) to model juvenile density (Supporting Information Table S2). There were significant trends in just over half of these (Supporting Information Table S2), but trajectories varied (Fig. 3 & Supporting Information Figs. S1 – S80). Data spatial and temporal coverage was most extensive in Wales and the south west, with model explanatory power good (R^2^ >= 30%) only in the north east, Wales and the north west (Supporting Information Table S2). Few catchments (n = 11) saw an increase in juvenile density, with the north west region having the largest number (Fig. 4A). In the Thames region, the raw data showed a population crash after 2005 with no salmon recorded in subsequent surveys, but no model could be fitted to capture these patterns. Trends in the south west were consistently negative (e.g., Fig. 3Bi), with only sporadic examples of increasing populations across the southern, Severn, and Welsh regions (Supporting Information Figs. S12A, S36A, S59A). Generally, trends in juvenile density were variable, and adjacent catchments had divergent trajectories (Fig. 4A).

**Fig. 3:**
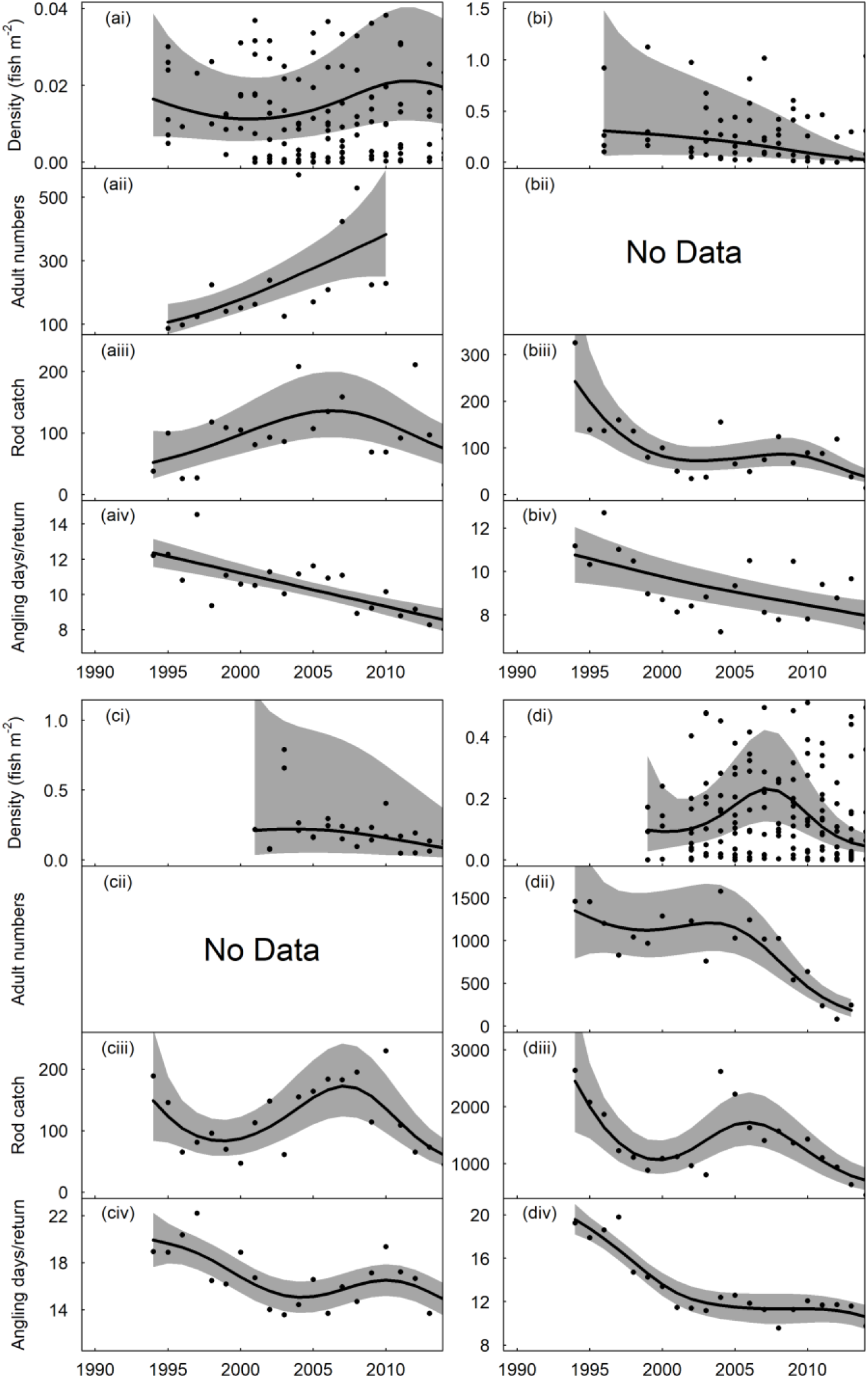
Trends in the a) River Tees (North East region), b) River Dart (South West region), c) Tawe (Wales region) and d) River Eden (North West region), chosen as representative examples of different trajectories observed. Boxes in each column represent: i) juvenile salmon density, fitted values from Gaussian additive mixed model, grey areas represent 95% point-wise confidence bands for the smoother, ii) returning adult numbers, fitted values from negative binomial generalized additive model, grey areas represent 95% point-wise confidence bands for the smoother, iii) rod catch, fitted values from negative binomial generalized additive model, grey areas represent 95% point-wise confidence bands for the smoother, and iv) angling effort measured as number of days fished per licence return, fitted values from Gaussian additive model, grey areas represent 95% point-wise confidence bands for the smoother. N.B. for ai and di plotting of raw values truncated to show trends.

**Fig. 4:**
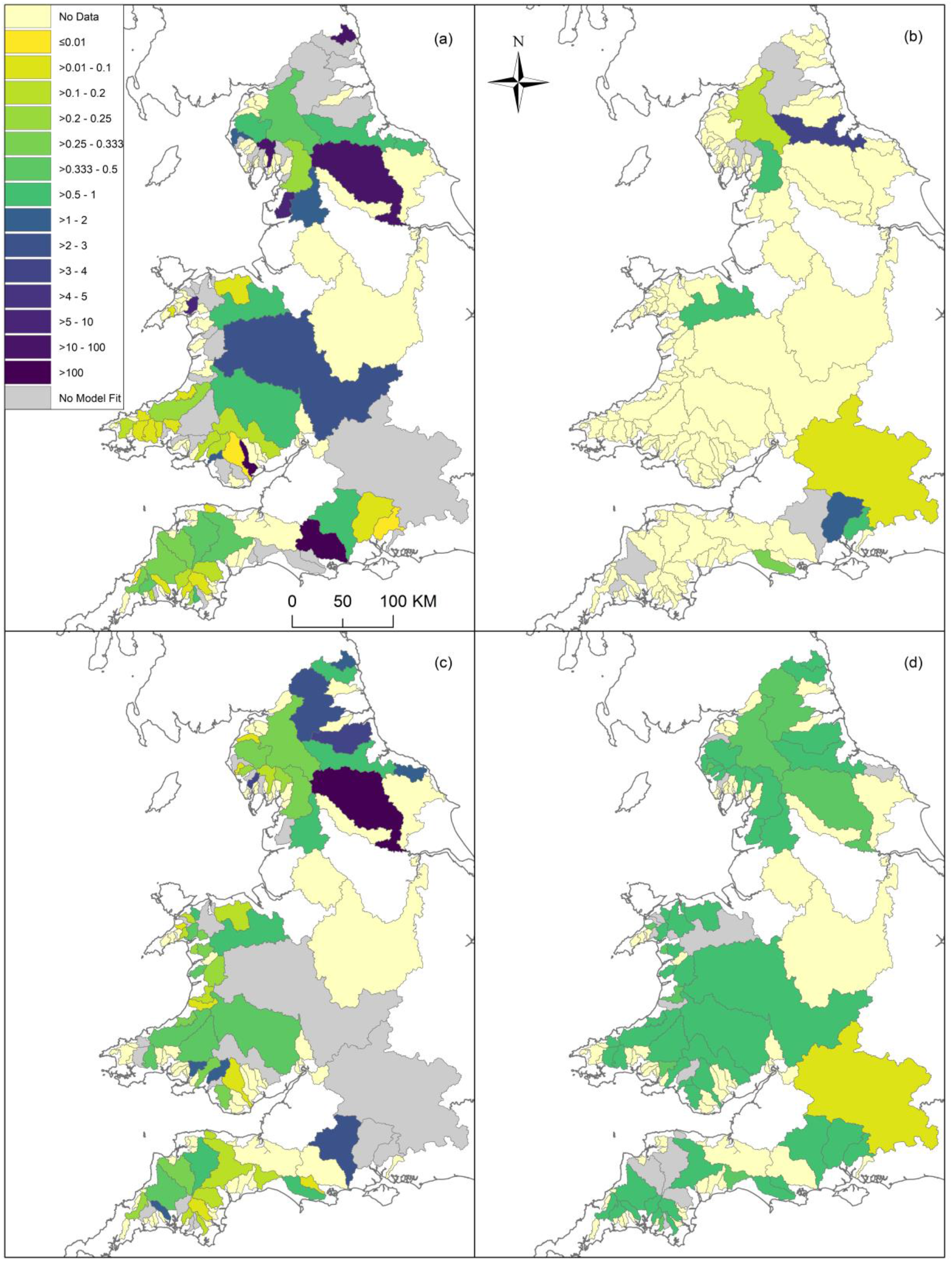
The change over time in A) juvenile density, B) returning adult numbers, C) total rod catch and D) angling effort, for salmon catchments across England and Wales. Change calculated as the ratio of the model mean value for the last year of data, divided by the model mean value of the first year of the dataset. For catchments where mean juvenile density for the latest year was predicted as less than zero, the prediction for the previous year was used to calculate change.

### Variable trends in returning adult numbers

The two models of returning adult numbers had strong model fit (mean annual number of returning adults <2500, *n* = 225, *R*^2^ = 0.79 and mean annual number of returning adults >4000, *n* = 89, *R*^2^ = 0.75). The fourteen catchments produced eight significant trends, with increases over time in returning adult numbers only observed in the River Tees (Fig 3Aii) and River Test (Supporting Information Fig. S10B) and an indication of potential recovery in the River Itchen (Supporting Information Fig. S9B). However, these increases in returning adult numbers were not matched by the trends in juvenile density. There was some evidence of analogous declining trends of juveniles and returning adults. Numbers of returning adults also exhibited a crash in the River Thames (Supporting Information Fig. S8B), with the declines in the River Eden (Fig 3Dii) and River Lune (Supporting Information Fig. S70B) operating on similar time scales.

### Rod catches tracked juvenile density only in catchments with negative trends

Actual ES use data in terms of total rod catch were available for over half the salmon catchments and produced 56 significant trends. Across the regions, model fit was generally high (Supporting Information Table S1). Increases in rod catches were most prevalent in the northeast catchments (Fig. 4C), although more recent reductions in catch were also observed (e.g., Fig 3Aiii & Supporting Information Figs. S2-4C). In the Thames region the raw data revealed a clear decline in adult rod catches that corresponded with the juvenile and adult data.

Despite some exceptions, the overall trend for southern, south west and Welsh regions was a decline in rod catches over the study period. In these regions only four catchments had mean modelled values higher in 2017 than in 1994 (Fig. 4C). In the northwest catchments the generally positive trends observed in the late 1990s and early 2000s have been negated by declines at the end of the timeseries (Supporting Information Fig. S68C & S70C). Increased mean modelled juvenile density was only translated to increased mean modelled rod catches for two catchments, one of which has seen rod catches decline for 2010 onwards (Supporting Information Fig. S1C). Conversely, declines in juveniles and returning adults were much more frequently translated to trends in rod catches (e.g. Fig. 3D).

### Angling effort declined and was unrelated to trends in juvenile density or adult rod catches

Across England and Wales actual ES use expressed as angling effort declined in every catchment (Fig. 4D). Declines in the number of days fished per licence return were apparent even in catchments that saw large increases in rod catches over time (e.g., Fig 3Aiv). Despite these overall declines, several catchments saw a stabilisation or increase in angling effort post 2005 (Supporting Information Figs. S25D, S34D, S51D).

## DISCUSSION

Overall, our results reveal that: i) juvenile Atlantic salmon densities presented non-uniform trends across the England and Wales, ii) trends in returning adult numbers were more closely matched to trends in juvenile density in catchments where juveniles were declining, iii) actual ES use in terms of adult rod catches more frequently tracked juvenile density and returning adult numbers in catchments with negative trends and iv) actual ES use in the form of angling effort declined consistently across England and Wales and was unrelated to trends in juvenile density or adult rod catches. Overall, our results show that along the ES cascade, from juvenile population abundance to ES use, there is some correlation between the elements of the cascade. However analogous trends are generally only present when juvenile populations are declining which permeates through to later levels of the cascade. This result corroborates findings in the terrestrial domain demonstrating that population decline has significant implications for ES provision (e.g., Gaston et al., 2018).

### Trends in juvenile populations vary across the country

Juvenile salmon density in English and Welsh catchments showed a range of trends. In the north (northeast, northwest and Humber regions), juvenile salmon density increased in certain catchments over the time series, corresponding with data from specific studies in the Tyne, Wear and Lune, which have also shown significant increases in adult numbers (Aprahamian, Wyatt, & Shields, 2006; CEFAS et al., 2015a; WWF, 2001). These trends have been attributed to the regulation of industrial and urban pollution that had previously decimated salmon populations (Mawle & Milner, 2003). In contrast, a discernible decline in juvenile salmon density was observed in the River Thames. While density was always relatively low, post 2005 no juvenile salmon were recorded, and this is thought to be a result of low flows and poor water quality in the lower river that act as a migratory barrier (Griffiths et al., 2011). Elsewhere trajectories of juvenile density through time were consistently negative (Fig. 4A). Where positive trends in juvenile density were observed they were in isolated catchments, challenging the assumption that adjacent rivers are subject to similar environmental conditions determining salmon production (Bowker, Ferns, Phillips, & Mawle, 1998; Milner, Davidson, Evans, Locke, & Wyatt, 2001). This work and others (e.g., Youngson et al., 2007) highlight how much populations of juvenile salmon fluctuate within time and space so that extensive datasets are required to identify any trends (CEFAS et al., 2015a).

### Linking salmonid population measures to ecosystem provision

Linking measures of biodiversity to ES provision remains a key challenge (Balvanera et al., 2014; Cardinale et al., 2012; Tolonen et al., 2014). For the Atlantic salmon, the difficulty lies not only in finding biodiversity measures that can be related effectively through the ES cascade to the recreational services and benefits (Fig. 1), but also in finding measures that account for their complex life history spent over large geographical ranges (Chaput, 2012; Hendry, Sambrook, & Waterfall, 2007). While there is extensive monitoring of juvenile populations, the relatively high capital costs of operating smolt and adult Atlantic salmon monitoring facilities, have significantly limited monitoring of adult populations in the UK (Cowx & Fraser, 2003; Youngson et al., 2007) and across the species’ distribution generally (ICES, 2018). Consequently, despite the extensive datasets curated in England and Wales, information on Atlantic salmon across the ES cascade is limited to a few catchments. This means that, currently, efforts to link Atlantic salmon biodiversity to ES provision can only realistically rely on measures of juvenile populations.

Our findings also seem to support the idea that the B-ES relationship is not necessarily linear and positive (Cardinale et al., 2012; de Groot, Alkemade, Braat, Hein, & Willemen, 2010; Duncan, Thompson, & Pettorelli, 2015; Harrison et al., 2014). Indeed, our results show a mixed correspondence in the trends of juvenile density, returning adult numbers and the two measures of ‘actual ES use’: rod catches and angling effort. Analogous responses of adult rod catches were generally only apparent in catchments where trends in juvenile density were negative. Increases in juvenile density were only translated to clear increases in rod catches for the Upper Ouse. In addition, despite certain catchments exhibiting increases in either returning adult numbers (potential ES use) or rod catches, trends in the other measure of actual ES use, angling effort, was unrelated. Across England and Wales as a whole, angling effort (the number of days fished) has reduced by over half since 1994 (CEFAS et al., 2015b).

### Assumptions in B-ES Linkages

The variable results in B-ES linkages are also likely to stem from the many assumptions around what characterises and determines the angling ES. For the Atlantic salmon, the size of the ‘potential use’ component (e.g. adult numbers) may not be the main determining factor of ‘actual use’ or ‘benefits to people’ (Fig. 1). Anglers display individual behaviours and abilities (Youngson, MacLean, & Fryer, 2002) so that rod catches and effort may not in fact reflect population trends (Hendry et al., 2007). Numbers of adults alone is unlikely to be the sole factor affecting angling effort, as participation is affected by a range of socio-economic elements (Aprahamian et al., 2010; Potter, MacLean, Wyatt, & Campbell, 2003) and satisfaction influenced by a variety of factors outside actually catching a fish (Arlinghaus, 2006). On the River Lune for example, increased adult salmon numbers resulted in significant increases in the proportion of anglers catching salmon and increases in the number of salmon caught per angler; however, a significant decline in the number of anglers was observed (Aprahamian et al., 2010). Further, prevailing environmental conditions, such as discharge, may have complementary or conflicting impacts on salmon populations (Nislow & Armstrong, 2012) and angler catch (Aprahamian & Ball, 1995; L’Abée-Lund & Aspås, 1999).

### Novel Approaches

To efficiently monitor changes in Atlantic salmon populations, while also better capturing the services or benefits they provide to people, we propose several strategies.

#### Better monitoring across the whole life cycle from juveniles to adult returning fish

While long-term data sites are central to detecting shifts in abundance, and considerable amounts of data have been collected, just over half of the catchments identified as supporting salmon populations had monitoring sites with more than 10 years of data. It should be noted that for a proportion of these 141 catchments salmon population are historically small and therefore recorded captures are sporadic. However, most catchments with suitable data had fewer than five sites with which to build a model. While juvenile monitoring is clearly valuable in appraising local effects on production, recruitment and inter-site variations, for example as a result of habitat quality, pollution and catchment land use, it is unlikely to be a good estimator of production at scales above the site level (Hendry et al., 2007). Greater spatial coverage in the monitoring of downstream migrating smolts and returning adults, based on existing and new techniques (e.g., eDNA, Levi et al., 2019).applied alongside traditional juvenile monitoring, would allow early detection of population bottlenecks (e.g., freshwater vs. marine stressors) At the moment, such a joined up approach is limited to a few ‘index’ rivers (CEFAS et al., 2015a), which cover a small proportion of the species’ global distribution (Prévost, 2015).

#### Better capture of the benefits derived from Atlantic salmon populations

Given the need to assess not just the ES quality at a given time but also its resilience, ES measures need to integrate both the immediate and long-term ecological mechanisms that support these services. Also, ideally, given the wider societal benefits of Atlantic salmon, particularly non-use values, their quantification will need to range across the spectrum of monetary, non-monetary and qualitative methods (Daniel et al., 2012), in essence to go beyond simple, market-orientated valuation approaches (Carpenter et al., 2009; Chan, Guerry, et al., 2012; Chan, Satterfield, & Goldstein, 2012; Daily et al., 2009; Plieninger et al., 2013). In this context, citizen science initiatives could be used - alongside more traditional measures on the amount of time fished and the number and size of salmon caught - to collect data on distance walked and levels of enjoyment thereby providing a more complete indication of the societal benefits associated with Atlantic salmon (Krasny, Russ, Tidball, & Elmqvist, 2014).

Our evidence suggests that it is currently challenging to clearly link our main Atlantic salmon biodiversity monitoring effort focused on a juvenile density to ES provision. Further investments on adult monitoring would allow to better bridge the link between the extensive juvenile population data and ecosystem provision. We suggest that utilising these monitoring data on both juvenile and adult stages of these migratory fish is key for a more integrated characterisation of ES delivery would both facilitate conservation efforts and increase capacity to monitor ES sustainability.

## AUTHORS’ CONTRIBUTIONS

TW, ID and SO conceived the ideas and designed methodology; TW with input from IV analysed the data; TW led the writing of the manuscript. All authors contributed critically to the drafts and gave final approval for publication.

## ACKNOWLEDGEMENTS

The work was carried out under the Diversity in Upland Rivers for Ecosystem Service Sustainability (DURESS) project (Grant reference NERC NE/J014818/1). The DURESS project was funded by the Natural Environment Research Council (NERC), through the Biodiversity and Ecosystem Service Sustainability (BESS) programme. T. W. was also supported by the MARS project (Managing Aquatic ecosystems and water Resources under multiple Stress), funded by the European Union under the 7th Framework Programme, contract no. 603378. The data were provided by the Environment Agency and Natural Resources Wales (Contains Natural Resources Wales information © Natural Resources Wales and database right. All rights reserved). We thank K. Whitlock, G. Peirson and M. Diamond of the Environment Agency, and I. Davidson and P. Gough of Natural Resources Wales for helpful comments on an earlier draft of the manuscript.

