## Supporting Information for "Recreational fisheries as a driver of salmonid population conservation"

### Supporting Information for
### "Recreational fisheries as a driver of salmonid population conservation"

#### This supporting information includes the following:

1. **Appendix S1:** Supplementary Introduction
2. **Appendix S2:** Supplementary Materials and Methods and Table S1: The structure of the different models across the four Atlantic salmon and ecosystem service datasets.
3. **Appendix S3:** Supplementary Results, Table S2: The 141 catchments identified for analysis of trends in juvenile salmon density, the number of sites within the catchment that have at least 10 years of data (with at least 6 years where salmon were recorded), whether a trend in juvenile salmon density was established, and the area of the catchment, and Figs S1-80: Trends in juvenile salmon density, rod catches, and angling effort for study rivers.

**APPENDIX S1: Supplementary Introduction**

Although life histories can vary spatially and temporally, the general pattern for Atlantic salmon is that of reproduction in freshwater during autumn or winter, whereby females release eggs into an area of prepared gravel (a ‘redd’) where they are fertilized by males (Fleming, 1998; Jonsson & Jonsson, 2011). Eggs hatch the following spring, and juveniles spend one to eight years in freshwater (Klemetsen et al., 2003). When juveniles reach a certain size (~15 cm) they undergo a series of physiological, behavioural, and morphological adaptations and migrate downstream as smolts to the sea (Jonsson & Jonsson, 2011; McCormick, Hansen, Quinn, & Saunders, 1998). Atlantic salmon remain in the sea to grow for up to five years before reaching maturity and returning to their natal river to spawn (Jonsson & Jonsson, 2011; Klemetsen et al., 2003). That being said, different life histories occur. For example, populations of non-anadromous or land-locked Atlantic salmon are apparent (Klemetsen et al., 2003; Mills, 1989), as well as males within a population maturing and reproducing within freshwater prior to seaward migration (Jonsson & Jonsson, 2011; Klemetsen et al., 2003).

**APPENDIX S2: Supplementary Materials and Methods**

Study area

To provide a framework for the analysis, England and Wales were split into 141 'salmon catchment' units (SI Appendix, Table. S2). Catchments supporting salmon populations were identified by superimposing the location of fisheries surveys (see below) that recorded Atlantic salmon over the Water Framework Directive (WFD) River Waterbody Catchments (Draft 2, available at http://www.geostore.com/environment-agency, herein referred to as 'WFD subcatchments') layer within a GIS software (ArcGIS). Salmon catchments were delimited by combining WFD subcatchments to form a watershed that encompassed a single river and tributaries from the headwaters to the estuary. A small number of catchments with only one or two records of Atlantic salmon were excluded from the analysis, as were samples from the Rivers Esk and Sark which cross the England-Scotland border.

To examine trends in different areas of England and Wales, the salmon catchments were split into eight regions (Fig. 2). The delimitation of regions was generally based on the UK's River Basin Districts (RBDs) (https://www.gov.uk/government/publications/river-basin-district-map). However, adjustments to the boundaries were made to better link the analysis with the exploitation of Atlantic salmon in terms of commercial net and recreational rod catches. The changes generally reflected the location of mixed stock fisheries (MSF) for Atlantic salmon, the presence of which makes determining the origin of the caught fish in commercial net catches impossible without genetic analysis. We took the definition of a MSF to be a fishery operating outside a river's estuary limits (Potter & Ó Maoiléidigh, 2006). Potter & Ó Maoiléidigh (2006) highlighted 10 Atlantic salmon MSFs operating within English and Welsh waters; furthermore, we used the definition above and descriptions and locations of commercial fisheries from Russel et al. (1995) and CEFAS, Environment Agency, & Natural

Resources Wales (2015b), to alter the regional boundaries by placing all catchments contributing to a MSF into the same region (for example, the Yorkshire Esk was moved from the Humber to the North East region). In addition to the presence of MSFs, the boundaries of the RBDs were realigned based on the physical structure of the rivers. Specifically, this was related to the inclusion of all chalk rivers into a single 'Southern' region.

Juvenile density - a traditional 'biodiversity' measure for population monitoring

Environment Agency (EA) and National Resources Wales' (NRW) monitoring programme for juvenile Atlantic salmon (fry and parr) is split into two main components: temporal sites which are surveyed biennially and spatial sites which are sampled every six years (CEFAS, Environment Agency, & Natural Resources Wales, 2015a). These are supplemented with sites surveyed specifically for the WFD (every 6 years) and local investigations (CEFAS et al., 2015a). Surveys are generally completed between June and September using electric fishing (CEFAS et al., 2015a). Data on juvenile salmon were available in three components

1) An original version of the National Fish Population Database (NFPD) containing fisheries surveys from England and Wales between 1990 and 2013 (requested from the EA)
2) An updated version of the NFPD containing fisheries surveys from England between 1975 and 2018 (available at: https://data.gov.uk/dataset/d129b21c-9e59-4913-91d2-82faef1862dd/nfpd-freshwater-fish-survey-relational-datasets).
3) Wales Freshwater Fish Data containing fisheries surveys from Wales between 2014 and 2018 (requested from the NRW)

The three components were merged to form a single dataset, whilst ensuring that surveys were only included once. The combined dataset consisted of 70344 surveys across England and Wales between 1975 and 2018. The vast majority of salmon recorded in the databases are juveniles. Of the 1,078,621 individual salmon captures recorded in the original NFPD, length data are available for 28.7% of these; only 0.58% are larger than the value of 160 mm for adult Atlantic salmon (Aprahamian, Davidson, & Cove, 2008; Forseth, Letcher, & Johansen, 2011).

The dataset was then filtered to use in the models. Surveys that were collected using either a semi-quantitative single catch samples (1 run) or a quantitative catch depletion samples (2 or more runs) were retained in the database. However, as converting between Catch per Unit Effort (CPUE) and the other methods may introduce errors into the models, CPUE surveys were removed. Further, only samples collected using electric fishing were retained. The number of salmon recorded in every survey at each site was determined. At sites where multiple surveys were taken in the same year, the yearly sum was calculated (Clews, Durance, Vaughan, & Ormerod, 2010). Only sites that had 10 or more years of data, with at least 6 years where salmon were recorded, were retained.

Because there was a difference in effort in terms of the number of electric fishing runs between the semi–quantitative and quantitative methodologies, only the first run of the quantitative surveys was used to provide a relative index of salmon abundance, rather than absolute numbers. Surveys with missing data on the area sampled (n = 7) were assigned an area based on the mean of other surveys from the same location. Density was calculated by dividing the number of Atlantic salmon from run 1 by the area surveyed. A final dataset of 7526 surveys at 501 locations was used in the analysis. Each survey location was spatially assigned to one of the 141 salmon catchments.

Returning adult numbers – a measure of potential ES use

Data on the number of returning adult Atlantic salmon, surveyed using electronic fish counters or traps, were accessed for 14 catchments from the annual Salmon Stocks and Fisheries in England and Wales reports (CEFAS & Environment Agency, 2000, 2002, 2005; CEFAS et al., 2015b; CEFAS, Environment Agency, & Natural Resources Wales, 2018). Counters or traps were only used if they provided 10 or more years of data, with temporal coverage ranging between 12 and 30 years. Catches are retrospectively updated and corrected in subsequent reports (CEFAS et al., 2015a); therefore, the most recent data were used. Analysis on the River Leven should be treated with caution as data from before 2000 only recorded salmon greater than 4 lb (1.8 kg).

Rod catches and angling effort – measures of actual ES use

To examine whether trends in juvenile density and adult fish are matched by changes in provision of certain cultural ecosystem services associated with Atlantic salmon, data were accessed from the Environment Agency's Salmonid & Freshwater Fisheries Statistics for England & Wales (1993 – 2017) reports. Actual ES use and benefits to people were examined in terms of 1) annual rod catches of Atlantic salmon (actual ES use), 2) angling effort in terms of the reported number of days fished for salmon and migratory trout per license return (actual ES use), and 3) regional sales of rod fish licences for salmon and sea trout (benefits to people).

Changes in how rod catch licence return data are collected (a national reminder system was initiated in 1994 that led to a significant increase in returns) meant that data on rod catches and angling effort were accessed for the period 1994 – 2017. Annual numbers of rod caught salmon recorded from licence returns are available for the 'principal' salmon rivers across

England and Wales, while effort data are available for a subset of the licence returns and expressed as the ‘number of days fished per return’, with CPUE the ‘catch per licence day’.

While the temporal availability of data on juvenile density varied between catchments (therefore, so did the length of the modelled trends), rod catch data were more consistent with trends created for the whole period between 1994 and 2017 (exceptions being the Rivers Axe and Erme (2002 – 2017), Yealm (1996 – 2017), and Crake (2001 – 2017)). Data on angling effort was available for all ‘principal’ rivers for the entire period between 1994 and 2017. It should be noted that effort data recorded for the licence returns was for both salmon and sea trout, and only provides an indication of changes in angling behaviour.

Statistical analysis

To examine temporal trends in juvenile Atlantic salmon density, generalized additive mixed models (GAMMs) were fitted using the package ‘mgcv’ in R version 3.4.1 (R Core Team, 2018; Wood, 2004, 2011). A separate GAMM was fitted to density data for each of the eight regions (Fig. 2). The GAMMs used a thin plate regression spline for the smoother and were fitted with restricted maximum likelihood. The maximum effective degrees of freedom (EDF) was set to seven to capture the overall trend and multiyear fluctuations (Fewster, Buckland, Siriwardena, Baillie, & Wilson, 2000); however, in this instance, the EDF was also allowed to take lower values resulting in simpler trends (e.g., straight lines). Models were fitted using the Gaussian family (Table S1). Prior to the analysis, the density data were natural log transformed with the addition of a constant (0.001) to account for zero densities. Given the aim to examine differences between Atlantic salmon stocks at the level of an individual river catchment, a base model of the interaction between year and salmon catchment allowing the non-linear relationship between density and time to take a different shape across individual catchments was used. For the Humber and Thames regions, suitable data were only available for a single salmon catchment; therefore, the interaction term was not required and a single smoother was fitted (Table S1). Extending the base model to include a random effect for sampling site was assessed using a likelihood ratio test as the models were nested (Zuur, Ieno, Walker, Saveliev, & Smith, 2009). Model validation was undertaken by creating histograms of the normalized residuals, and plotting the normalized residuals against the fitted values and the covariates, year and catchment (Zuur, Ieno, & Elphick, 2010; Zuur, Saveliev, & Ieno, 2014). The residual plots suggested heteroscedasticity was an issue; therefore, the inclusion of different variance structures was tested (Zuur et al., 2009) and the structure producing the lowest AIC retained. A structure that allowed different variances across the catchments resulted in the lowest AIC value for the majority of regions (Table S1). The models were further extended to correct for temporal autocorrelation by testing model fit when incorporating an auto-regressive moving average (ARMA) structure with lags of 1, 2 or 3; however, in a few instances the full set of models could not be tested as certain models would not converge (Table S1). With the exception of the Thames region, for which the model was rejected, the model validation procedure was repeated and the inclusion of variance and/or autocorrelation structures significantly improved the appearance of the residual plots. Spatial autocorrelation was assessed by using bubble plots from the 'sp' package (Bivand, Pebesma, & Gomez-Rubio, 2013; Pebesma & Bivand, 2005) to plot the normalized residuals in space, with the residuals scaled for size. No patterns were detected.

Trends in returning adult numbers Atlantic salmon numbers were assessed using generalized additive models (GAMs) using the package 'mgcv' in R version 3.4.1 (R Core Team, 2018; Wood, 2004). Given the paucity of data only two GAMs were fitted, one for those rivers where mean annual number of returning adults was <2500 and the other for rivers where mean annual number of returning adults was >4000 (Table S1). The model setup, in terms of the amount of smoothing and spline used, followed that of the GAMM analysis for juvenile density. A base model was fitted which allowed the shape of the relationship between total catch and year to vary between salmon catchments. Models were fitted using the Poisson family; however, after checking for overdispersion, a negative binomial was used (Table S1). Model assumptions were assessed by examining plots of the Pearson residuals against the fitted values and the covariates, year and catchment (Zuur, 2012; Zuur et al., 2010).

Trends in actual ES use, total rod salmon catches, were assessed using generalized additive models (GAMs) using the package ‘mgcv’ in R version 3.4.1 (R Core Team, 2018; Wood, 2004). A separate GAM was fitted to total catch from each of the eight regions (Fig. 2). The model setup, in terms of the amount of smoothing and spline used, followed that of the GAMM analysis for juvenile density. A base model was fitted to each region which allowed the shape of the relationship between total catch and year to vary between salmon catchments (for the Humber and Thames regions, there was only one catchment with data; therefore, the interaction term was not required). Models were fitted using the Poisson family; however, after checking for overdispersion, a negative binomial was used (Table S1). Model assumptions were assessed using the same approach as for adult Atlantic salmon numbers. The residual plots for the Thames region suggested that the model assumptions had not been met; therefore, the model was not retained. For several regions (Table S1), given large differences in the magnitude of yearly rod catches between catchments in each region, two models (small and large catch catchments) were fitted, improving the appearance of the residuals plots.

Trends in actual ES use, angling effort, were also assessed using GAMs using the package ‘mgcv’ in R version 3.4.1 (R Core Team, 2018; Wood, 2004). Again, a separate model was fitted to the effort data for each region, with a model structure where the shape of the relationship between effort and year varied between salmon catchments. Models were fitted using the Gaussian family. The model setup, in terms of the amount of smoothing, the spline used, model validation and procedures to deal with heteroscedasticity and temporal autocorrelation, followed that of the GAMM analysis (Table S1). For two regions, an outlier was removed following inspection of the model validation plots (Table S1).

Table S1: The structure of the different models across the four Atlantic salmon and ecosystem service datasets.

| | | Distribution | | | | Smoother | | | | Variance structure | | | | Temporal autocorrelation | | |
|---|---|---|---|---|---|---|---|---|---|---|---|---|---|---|---|---|
| Stage | Region | Gaussian | Poisson | Negative Binomial | Data split by magnitude of response | Single smoother | Single smoother + factor | Catchment x smoother interaction | Random Effect | varIdent | varFixed | varPower | varConstPower | Lag 1 | Lag 2 | Lag 3 |
| Juveniles | Northeast | ● | — | — | | | | ● | ● | ● | | | | | | ● |
| | Humber | ● | — | — | | ● | — | — | ● | — | | | | | ● | |
| | Thames | No Model | | | | | | | | | | | | | | |
| | Southern | ● | — | — | | | | ● | ● | ● | | | | ● | | |
| | Southwest | ● | — | — | | | | ● | ● | ● | | | | ● | C | C |
| | Severn | ● | — | — | | | | ● | ● | | | | | | | ● |
| | Wales | ● | — | — | | | | ● | ● | ● | | | | | | ● |
| | Northwest | ● | — | — | | | | ●* | ● | ● | | | | | | ● |
| Adults | | — | | ● | ● | | | ● | — | — | — | — | — | — | — | — |
| Rod Catches | Northeast | — | | ● | ● | | | ● | — | — | — | — | — | — | — | — |
| | Humber | — | | ● | | ● | — | — | — | — | — | — | — | — | — | — |
| | Thames | No Model | | | | | | | | | | | | | | |
| | Southern | — | | ● | | | | ● | — | — | — | — | — | — | — | — |
| | Southwest | — | | ● | | | | ● | — | — | — | — | — | — | — | — |
| | Severn | — | | ● | | | | ● | — | — | — | — | — | — | — | — |
| | Wales | — | | ● | ● | | | ● | — | — | — | — | — | — | — | — |
| | Northwest | — | | ● | ● | | | ● | — | — | — | — | — | — | — | — |

| | | | | | | | | | | | | |
|---|---|---|---|---|---|---|---|---|---|---|---|---|
| Angling Effort | Northeast | • | — | — | ** | | | • | — | • | | |
| | Humber | • | — | — | ** | • | — | — | — | — | | |
| | Thames | • | — | — | | • | — | — | — | — | | |
| | Southern | • | — | — | | | | • | — | • | | |
| | Southwest | • | — | — | | | | • | — | • | • | C |
| | Severn | • | — | — | | | | • | — | • | | |
| | Wales | • | — | — | | | | • | — | • | | |
| | Northwest | • | — | — | | | | • | — | • | • | |

— not applicable to this model; C model couldn't converge therefore excluded, * model didn't result in lowest AIC value but structure retained for consistency, ** extreme value removed.

#### APPENDIX S3: Supplementary Results

Table S2: The 141 catchments identified for analysis of trends in juvenile salmon density, the number of sites within the catchment that have at least 10 years of data (with at least 6 years where salmon were recorded), whether a trend in juvenile salmon density was established, and the area of the catchment.

| ID | Salmon Catchment | Area ($km^2$) | Sites | Ecosystem Organisation Juvenile Density Model | | | | Potential ES Use Returning Adult Model | | | | Actual ES Use Rod Catch Model | | | | Actual ES Use Angling Effort Model | | | |
|---|---|---|---|---|---|---|---|---|---|---|---|---|---|---|---|---|---|---|---|
| | | | | **EDF** | ***F*** | ***p*** | **Trend** | **EDF** | $\chi^2$ | ***p*** | **Trend** | **EDF** | $\chi^2$ | ***p*** | **Trend** | **EDF** | ***F*** | ***p*** | **Trend** |
| **North West** | | | | $R^2 = 0.42$, n = 766 | | | | | | | | Small catches: $R^2 = 0.57$, n = 72<br>Large catches: $R^2 = 0.88$, n = 72 | | | | $R^2 = 0.90$, n = 143 | | | |
| 1 | R. Aln | 248 | 4 | 1.00 | 14.21 | **0.0002** | ↗ | - | - | - | - | 3.83 | 25.16 | **<0.0001** | ↗↘ | 1.91 | 3.70 | **0.02** | ↘→ |
| 2 | R. Coquet | 597 | 7 | 1.00 | 0.10 | 0.75 | - | - | - | - | - | 3.57 | 40.29 | **<0.0001** | ↗↘ | 3.85 | 23.93 | **<0.0001** | ↘→ |
| 3 | R. Wansbeck | 330 | 1 | 1.00 | 1.14 | 0.29 | - | - | - | - | - | - | - | - | - | - | - | - | - |
| 4 | R. Tyne | 2269 | 14 | 1.01 | 0.60 | 0.44 | - | 1.01 | 0.11 | 0.75 | | 2.89 | 48.37 | **<0.0001** | ↗↘ | 4.02 | 91.56 | **<0.0001** | ↘ |
| 5 | R. Derwent | 266 | 0 | - | - | - | - | - | - | - | - | - | - | - | - | - | - | - | - |
| 6 | R. Wear | 1107 | 7 | 1.80 | 2.47 | 0.06 | - | - | - | - | - | 3.55 | 115.60 | **<0.0001** | ↗↘ | 1.79 | 16.65 | **<0.0001** | ↘ |
| 7 | R. Tees | 1533 | 16 | 3.46 | 3.07 | **0.02** | ↘↗↘ | 1.36 | 14.20 | **0.01** | ↗ | 2.45 | 8.46 | **0.04** | ↗↘ | 1.00 | 40.04 | **<0.0001** | ↘ |
| 8 | Y. Esk | 348 | 3 | 3.61 | 2.24 | **0.04** | ↗↘ | - | - | - | - | 3.66 | 17.94 | **0.002** | ↗→ | 1.91 | 0.68 | 0.46 | - |
| **Humber** | | | | $R^2 = 0.00$, n = 215 | | | | | | | | $R^2 = 0.94$, n = 24 | | | | $R^2 = 0.81$, n = 23 | | | |
| 9 | Y. Derwent | 1994 | 0 | - | - | - | - | - | - | - | - | - | - | - | - | - | - | - | - |
| 10 | Upper Ouse | 3860 | 13 | 3.52 | 5.18 | **0.002** | ↗ | - | - | - | - | 6.49 | 400.8 | **<0.0001** | ↗ | 5.75 | 15.46 | **<0.0001** | ↘→ |
| 11 | R. Wharfe & Ouse | 1001 | 0 | - | - | - | - | - | - | - | - | - | - | - | - | - | - | - | - |

| | | | | | | | | | | | | | | | | | | | |
|---|---|---|---|---|---|---|---|---|---|---|---|---|---|---|---|---|---|---|---|
| 12 | R. Trent | 9011 | 0 | - | - | - | - | - | - | - | - | - | - | - | - | - | - | - | - |
| **Thames** | | | | N/A | | | | | | | | N/A | | | | $R^2$ = 0.68, n = 24 | | | |
| 13 | R. Thames | 9953 | 2 | - | - | - | - | 4.60 | 227.53 | **<0.0001** | ↗↘ | - | - | - | - | 1.00 | 49.16 | **<0.0001** | ↘ |
| **Southern** | | | | $R^2$ = 0.13, n = 199 | | | | | | | | $R^2$ = 0.69, n = 120 | | | | $R^2$ = 0.85, n = 120 | | | |
| 14 | R. Meon | 108 | 1 | 1.00 | 0.05 | 0.82 | - | - | - | - | - | - | - | - | - | - | - | - | - |
| 15 | R. Itchen | 464 | 1 | 1.15 | 5.50 | **0.02** | ↘ | 3.38 | 21.38 | **0.0003** | ↘↗ | 1.00 | 1.20 | 0.27 | - | 4.86 | 27.15 | **<0.0001** | ↘↗→ |
| 16 | R. Test | 1200 | 1 | 1.00 | 6.02 | **0.02** | ↘ | 3.29 | 19.34 | **0.0007** | ↘↗ | 2.93 | 6.88 | 0.12 | | 2.64 | 4.83 | **0.02** | ↘→ |
| 17 | H. Avon | 1708 | 6 | 3.26 | 4.61 | **0.003** | ↗↘ | 1.84 | 0.63 | 0.66 | | 1.41 | 7.74 | **0.03** | ↗ | 2.46 | 19.07 | **<0.0001** | ↘ |
| 18 | R. Stour | 1238 | 1 | 1.00 | 18.43 | **<0.0001** | ↗ | - | - | - | - | - | - | - | - | - | - | - | - |
| 19 | R. Piddle | 185 | 2 | 1.00 | 0.71 | 0.40 | - | - | - | - | - | 1.73 | 24.65 | **<0.0001** | ↘ | 1.00 | 4.76 | **0.03** | ↘ |
| 20 | R. Frome | 460 | 4 | 2.83 | 2.74 | 0.12 | - | 2.53 | 27.33 | **<0.0001** | ↘→ | 2.38 | 12.65 | **0.005** | ↘↗ | 3.47 | 35.65 | **<0.0001** | ↘ |
| **South West** | | | | $R^2$ = 0.13, n = 2543 | | | | | | | | $R^2$ = 0.75, n = 366 | | | | $R^2$ = 0.77, n = 384 | | | |
| 21 | R. Axe | 305 | 1 | 1.00 | 0.23 | 0.63 | - | - | - | - | - | 3.92 | 13.59 | **0.05** | ↘↗↘ | 1.00 | 52.32 | **<0.0001** | ↘ |
| 22 | Umbourne Brook | 82 | 1 | 1.00 | 0.60 | 0.44 | - | - | - | - | - | - | - | - | - | - | - | - | - |
| 23 | R. Otter | 207 | 0 | - | - | - | - | - | - | - | - | - | - | - | - | - | - | - | - |
| 24 | R. Clyst | 145 | 0 | - | - | - | - | - | - | - | - | - | - | - | - | - | - | - | - |
| 25 | R. Exe | 1191 | 16 | 3.60 | 6.10 | **0.0004** | →↗↘ | - | - | - | - | 3.25 | 28.30 | **<0.0001** | ↘ | 1.00 | 10.03 | **0.002** | ↘ |
| 26 | R. Teign | 398 | 5 | 1.36 | 5.33 | **0.008** | ↘ | - | - | - | - | 3.41 | 22.60 | **0.0002** | ↘ | 1.00 | 0.65 | 0.42 | - |
| 27 | R. Lemon | 43 | 0 | - | - | - | - | - | - | - | - | - | - | - | - | - | - | - | - |
| 28 | Am Brook | 51 | 0 | - | - | - | - | - | - | - | - | - | - | - | - | - | - | - | - |

| | | | | | | | | | | | | | | | | | | | |
|---|---|---|---|---|---|---|---|---|---|---|---|---|---|---|---|---|---|---|---|
| 29 | R. Dart | 289 | 5 | 2.42 | 13.85 | **<0.0001** | ↘ | - | - | - | - | 3.66 | 40.81 | **<0.0001** | ↘ | 1.34 | 10.34 | **0.001** | ↘ |
| 30 | R. Harbourne | 46 | 0 | - | - | - | - | - | - | - | - | - | - | - | - | - | - | - | - |
| 31 | R. Avon (Devon) | 106 | 2 | 2.30 | 2.43 | 0.06 | - | - | - | - | - | 3.78 | 28.27 | **<0.0001** | ↘↗↘ | 3.24 | 13.08 | **<0.0001** | ↘→ |
| 32 | R. Erme | 62 | 2 | 3.09 | 9.75 | **<0.0001** | ↗↘ | - | - | - | - | 1.42 | 2.45 | 0.28 | - | 1.00 | 0.01 | 0.94 | - |
| 33 | R. Yealm | 58 | 5 | 2.79 | 15.19 | **<0.0001** | ↗↘ | - | - | - | - | 2.88 | 7.11 | 0.08 | - | 1.00 | 54.67 | **<0.0001** | ↘ |
| 34 | Tory Brook | 26 | 0 | - | - | - | - | - | - | - | - | - | - | - | - | - | - | - | - |
| 35 | R. Plym | 97 | 8 | 3.16 | 5.06 | **0.001** | ↘ | - | - | - | - | 3.40 | 26.27 | **<0.0001** | ↘ | 1.00 | 78.31 | **<0.0001** | ↘ |
| 36 | R. Tavy | 206 | 12 | 3.57 | 8.85 | **<0.0001** | ↘↗↘ | - | - | - | - | 2.83 | 7.92 | 0.07 | - | 1.00 | 10.58 | **0.001** | ↘ |
| 37 | R. Tamar | 928 | 38 | 4.51 | 11.93 | **<0.0001** | ↗→↘ | 1.69 | 0.85 | 0.74 | | 3.01 | 9.58 | **0.03** | ↘→ | 1.90 | 7.49 | **0.005** | ↘→ |
| 38 | R. Lynher | 152 | 3 | 3.12 | 5.15 | **0.001** | ↘ | - | - | - | - | 1.00 | 4.13 | **0.04** | ↗ | 3.97 | 6.81 | **<0.0001** | ↘↗ |
| 39 | Tiddy Brook | 37 | 0 | - | - | - | - | - | - | - | - | - | - | - | - | - | - | - | - |
| 40 | East Looe | 49 | 1 | 1.00 | 0.25 | 0.62 | - | - | - | - | - | - | - | - | - | - | - | - | - |
| 41 | West Looe | 45 | 1 | 1.00 | 4.35 | **0.04** | ↘ | - | - | - | - | - | - | - | - | - | - | - | - |
| 42 | R. Lerryn | 24 | 0 | - | - | - | - | - | - | - | - | - | - | - | - | - | - | - | - |
| 43 | R. Fowey | 175 | 20 | 1.00 | 6.17 | **0.01** | ↘ | 2.21 | 3.88 | 0.15 | | 1.19 | 0.35 | 0.59 | - | 3.52 | 6.82 | **0.0003** | ↘↗↘ |
| 44 | R. Camel | 217 | 12 | 4.48 | 7.52 | **<0.0001** | ↗↘ | - | - | - | - | 3.39 | 24.91 | **<0.0001** | ↘↗↘ | 3.26 | 13.02 | **<0.0001** | ↘ |
| 45 | R. Fal | 108 | 0 | - | - | - | - | - | - | - | - | - | - | - | - | - | - | - | - |
| 46 | R. Allen | 53 | 9 | 4.53 | 9.83 | **<0.0001** | ↗↘ | - | - | - | - | - | - | - | - | - | - | - | - |
| 47 | R. Valency | 20 | 0 | - | - | - | - | - | - | - | - | - | - | - | - | - | - | - | - |
| 48 | R. Torridge | 717 | 9 | 3.01 | 2.78 | **0.04** | ↘↗↘ | - | - | - | - | 3.66 | 17.13 | **0.003** | ↘→ | 1.71 | 2.30 | 0.07 | - |

| | | | | | | | | | | | | | | | | | | | |
|---|---|---|---|---|---|---|---|---|---|---|---|---|---|---|---|---|---|---|---|
| 49 | R. Yeo | 61 | 0 | - | - | - | - | - | - | - | - | - | - | - | - | - | - | - | - |
| 50 | R. Taw | 918 | 12 | 1.00 | 4.89 | **0.03** | ↘ | - | - | - | - | 1.00 | 4.01 | **0.05** | ↘ | 2.01 | 1.61 | 0.19 | - |
| 51 | Venn Stream | 37 | 0 | - | - | - | - | - | - | - | - | - | - | - | - | - | - | - | - |
| 52 | Barnstable Yeo | 86 | 0 | - | - | - | - | - | - | - | - | - | - | - | - | - | - | - | - |
| 53 | Bradiford Water | 32 | 0 | - | - | - | - | - | - | - | - | - | - | - | - | - | - | - | - |
| 54 | R. Lyn | 77 | 2 | 2.21 | 4.09 | **0.01** | ↘ | - | - | - | - | 2.32 | 56.56 | **<0.0001** | ↘ | 1.21 | 0.31 | 0.72 | - |
| **Severn** | | | | $R^2$ = 0.23, n = 1465 | | | | | | | | $R^2$ = 0.80, n = 120 | | | | $R^2$ = 0.88, n = 120 | | | |
| 55 | R. Washford | 44 | 0 | - | - | - | | - | - | - | - | - | - | - | - | - | - | - | - |
| 56 | R. Parret | 1590 | 0 | - | - | - | | - | - | - | - | - | - | - | - | - | - | - | - |
| 57 | R. Frome (Bristol) | 228 | 0 | - | - | - | | - | - | - | - | - | - | - | - | - | - | - | - |
| 58 | R. Severn | 10083 | 15 | 1.00 | 5.98 | **0.01** | ↗ | - | - | - | - | 1.94 | 3.04 | 0.33 | - | 3.48 | 29.62 | **<0.0001** | ↘→ |
| 59 | R. Leadon | 329 | 0 | - | - | - | | - | - | - | - | - | - | - | - | - | - | - | - |
| 60 | Westbury Brook | 32 | 0 | - | - | - | | - | - | - | - | - | - | - | - | - | - | - | - |
| 61 | Cinderford Brook | 50 | 1 | - | - | - | | - | - | - | - | - | - | - | - | - | - | - | - |
| 62 | R. Wye | 4057 | 35 | 4.75 | 3.15 | **0.04** | →↗↘ | - | - | - | - | 3.02 | 13.25 | **0.01** | ↘→ | 3.59 | 17.69 | **<0.0001** | ↘→ |
| 63 | R. Usk | 1119 | 35 | 5.89 | 30.48 | **<0.0001** | ↗↘ | - | - | - | - | 1.00 | 0.00 | 0.96 | - | 3.04 | 19.80 | **<0.0001** | ↘→ |
| 64 | Sor Brook | 27 | 0 | - | - | - | | - | - | - | - | - | - | - | - | - | - | - | - |
| 65 | R. Lywd | 106 | 0 | - | - | - | | - | - | - | - | - | - | - | - | - | - | - | - |
| 66 | R. Ebbw/Sirhowy | 220 | 0 | - | - | - | | - | - | - | - | - | - | - | - | - | - | - | - |
| 67 | R. Rhymney | 216 | 1 | 1.50 | 5.19 | **0.007** | ↗ | - | - | - | - | - | - | - | - | - | - | - | - |

| | | | | | | | | | | | | | | | | | | | |
|---|---|---|---|---|---|---|---|---|---|---|---|---|---|---|---|---|---|---|---|
| 68 | R. Taff | 529 | 1 | 5.34 | 3.77 | **0.001** | ↘→ | - | - | - | - | 5.49 | 54.65 | **<0.0001** | ↘↗↘ | 1.00 | 5.77 | **0.02** | ↘ |
| 69 | R. Ely | 167 | 1 | 2.96 | 1.77 | 0.18 | | - | - | - | - | - | - | - | - | - | - | - | - |
| 70 | R. Thaw | 123 | 0 | - | - | - | | - | - | - | - | - | - | - | - | - | - | - | - |
| 71 | R. Ogmore | 269 | 2 | 1.00 | 2.94 | 0.09 | | - | - | - | - | 2.57 | 22.41 | **<0.0001** | ↘ | 3.10 | 7.50 | **0.0001** | ↘↗ |
| **Wales** | | | | $R^2$ = 0.47, n = 1082 | | | | | | | | Small catches: $R^2$ = 0.79, n = 552<br>Large catches: $R^2$ = 0.38, n = 72 | | | | $R^2$ = 0.74, n = 624 | | | |
| 72 | Afan | 91 | 1 | 3.40 | 2.26 | **0.07** | ↗↘ | - | - | - | - | 2.19 | 3.68 | 0.35 | - | 2.13 | 4.11 | **0.01** | ↘→ |
| 73 | R. Neath | 264 | 2 | 2.29 | 2.74 | **0.04** | ↗↘ | - | - | - | - | 2.78 | 13.41 | **0.06** | ↗↘ | 1.11 | 2.52 | 0.10 | - |
| 74 | R. Tawe | 267 | 2 | 2.14 | 4.45 | **0.01** | ↘ | - | - | - | - | 4.06 | 25.02 | **0.0001** | ↘↗↘ | 3.91 | 7.67 | **<0.0001** | ↘ |
| 75 | R. Llan | 76 | 0 | - | - | - | | - | - | - | - | - | - | - | - | - | - | - | - |
| 76 | R. Loughor | 173 | 1 | 1.00 | 0.19 | 0.67 | | - | - | - | - | 3.64 | 35.60 | **<0.0001** | ↗↘ | 1.00 | 32.87 | **<0.0001** | ↘ |
| 77 | Gwili (Gower) | 32 | 0 | - | - | - | | - | - | - | - | - | - | - | - | - | - | - | - |
| 78 | Morlais | 24 | 0 | - | - | - | | - | - | - | - | - | - | - | - | - | - | - | - |
| 79 | Afan Goch | 77 | 0 | - | - | - | | - | - | - | - | - | - | - | - | - | - | - | - |
| 80 | Gwendraeth Fawr | 86 | 1 | 1.00 | 0.07 | 0.80 | | - | - | - | - | - | - | - | - | - | - | - | - |
| 81 | Tywi | 1114 | 8 | 2.74 | 2.52 | 0.10 | | - | - | - | - | 4.47 | 26.49 | **0.0001** | ↘↗↘ | 3.44 | 7.96 | **<0.0001** | ↘→ |
| 82 | Gwili | 151 | 1 | 1.54 | 3.62 | **0.02** | ↘ | - | - | - | - | - | - | - | - | - | - | - | - |
| 83 | Cywyn | 78 | 1 | 1.00 | 1.20 | 0.27 | | - | - | - | - | - | - | - | - | - | - | - | - |
| 84 | Cynin | 103 | 1 | 2.66 | 8.28 | **0.0001** | ↘ | - | - | - | - | - | - | - | - | - | - | - | - |
| 85 | Taf | 246 | 2 | 2.47 | 3.98 | **0.02** | ↗↘ | - | - | - | - | 3.16 | 12.80 | **0.01** | ↘↗↘ | 1.63 | 4.99 | **0.007** | ↘ |

| | | | | | | | | | | | | | | | | | | | |
|---|---|---|---|---|---|---|---|---|---|---|---|---|---|---|---|---|---|---|---|
| 86 | Eastern Cleddau | 207 | 2 | 2.53 | 4.86 | **0.01** | ↘ | - | - | - | - | 2.29 | 1.59 | 0.54 | - | 3.99 | 7.48 | **<0.0001** | ↘→↘ |
| 87 | Cartlett Brook | 36 | 0 | - | - | - | | - | - | - | - | - | - | - | - | - | - | - | - |
| 88 | Western Cleddau | 217 | 2 | 3.03 | 13.74 | **<0.0001** | →↘ | - | - | - | - | - | - | - | - | - | - | - | - |
| 89 | Solva | 35 | 0 | - | - | - | | - | - | - | - | - | - | - | - | - | - | - | - |
| 90 | Gwaun | 46 | 0 | - | - | - | | - | - | - | - | - | - | - | - | - | - | - | - |
| 91 | Nevern | 110 | 2 | 2.17 | 9.86 | **<0.0001** | →↘ | - | - | - | - | 1.00 | 0.17 | 0.68 | - | 2.80 | 6.90 | **0.0008** | ↘→ |
| 92 | Teifi | 951 | 12 | 3.19 | 9.50 | **<0.0001** | ↗↘ | - | - | - | - | 2.78 | 25.90 | **<0.0001** | →↘ | 2.44 | 4.75 | **0.008** | ↘↗ |
| 93 | Aeron | 162 | 2 | 3.18 | 7.48 | **<0.0001** | →↘ | - | - | - | - | 2.39 | 8.51 | **0.03** | →↘ | 2.70 | 8.82 | **<0.0001** | ↘→ |
| 94 | Ystwyth | 192 | 2 | 1.00 | 1.05 | 0.31 | | - | - | - | - | 4.28 | 32.49 | **<0.0001** | ↘ | 1.00 | 3.35 | 0.07 | - |
| 95 | Rheidol | 183 | 0 | - | - | - | | - | - | - | - | 3.34 | 23.31 | **0.0001** | ↘→↘ | 2.53 | 12.00 | **<0.0001** | →↘ |
| 96 | Dyfi | 476 | 3 | 2.17 | 1.72 | 0.19 | | - | - | - | - | 3.70 | 22.17 | **0.0003** | ↘↗↘ | 1.00 | 7.69 | **0.006** | ↘ |
| 97 | Dysynni | 126 | 0 | - | - | - | | - | - | - | - | 1.99 | 8.90 | **0.02** | ↘→ | 2.65 | 11.20 | **<0.0001** | ↘↗ |
| 98 | Wnion | 116 | 1 | 1.00 | 0.40 | 0.53 | | - | - | - | - | - | - | - | - | - | - | - | - |
| 99 | Mawddach | 165 | 0 | - | - | - | | - | - | - | - | 1.01 | 18.33 | **<0.0001** | ↘ | 3.06 | 4.71 | **0.003** | ↘↗ |
| 100 | Artro | 55 | 0 | - | - | - | | - | - | - | - | 1.00 | 9.12 | **0.003** | ↘ | 1.00 | 20.25 | **<0.0001** | ↘ |
| 101 | Dwyryd | 81 | 0 | - | - | - | | - | - | - | - | 4.34 | 33.99 | **<0.0001** | ↘↗↘ | 1.63 | 1.87 | 0.10 | - |
| 102 | Glaslyn | 150 | 1 | 2.41 | 8.80 | **<0.0001** | ↗ | - | - | - | - | 3.76 | 15.39 | **0.007** | ↘↗↘ | 3.90 | 15.19 | **<0.0001** | ↘↗↘ |
| 103 | Dwyfawr | 63 | 0 | - | - | - | | - | - | - | - | 2.94 | 17.94 | **0.001** | ↘ | 1.00 | 0.10 | 0.75 | - |
| 104 | Dwyfach | 47 | 0 | - | - | - | | - | - | - | - | - | - | - | - | - | - | - | - |
| 105 | Erch | 56 | 1 | 3.06 | 8.19 | **<0.0001** | ↗↘ | - | - | - | - | - | - | - | - | - | - | - | - |

| | | | | | | | | | | | | | | | | | | | |
|---|---|---|---|---|---|---|---|---|---|---|---|---|---|---|---|---|---|---|---|
| 106 | Rhyd-Hir | 74 | 0 | - | - | - | | - | - | - | - | - | - | - | - | - | - | - | - |
| 107 | Llyfni | 51 | 0 | - | - | - | | - | - | - | - | 1.08 | 75.03 | **<0.0001** | ↘ | 1.00 | 31.62 | **<0.0001** | ↘ |
| 108 | Gwyrfai | 54 | 0 | - | - | - | | - | - | - | - | 2.95 | 6.97 | 0.15 | - | 1.58 | 0.41 | 0.63 | - |
| 109 | Seiont | 80 | 1 | 1.00 | 0.07 | 0.79 | | - | - | - | - | 3.40 | 40.34 | **<0.0001** | ↗↘ | 1.00 | 0.22 | 0.64 | - |
| 110 | Ogwen | 82 | 1 | 1.63 | 0.27 | 0.65 | | - | - | - | - | 1.97 | 7.26 | **0.04** | →↘ | 1.00 | 8.04 | **0.005** | ↘ |
| 111 | Conway | 520 | 7 | 1.00 | 1.41 | 0.24 | | - | - | - | - | 1.65 | 3.39 | 0.18 | - | 2.76 | 3.36 | **0.01** | ↘↗ |
| 112 | Clwyd | 695 | 3 | 3.31 | 9.59 | **<0.0001** | ↗↘ | - | - | - | - | 6.21 | 62.67 | **<0.0001** | →↗↘ | 2.59 | 27.74 | **<0.0001** | ↘ |
| 113 | Dee | 1802 | 11 | 2.14 | 3.12 | **0.04** | ↗↘ | 2.51 | 17.56 | **0.0007** | →↘ | 3.47 | 15.05 | **0.006** | ↘↗↘ | 1.00 | 0.04 | 0.84 | - |
| **North West** | | | | $R^2$ = 0.31, n = 1227 | | | | | | | | Small catches: $R^2$ = 0.78, n = 233<br>Large catches: $R^2$ = 0.63, n = 96 | | | | $R^2$ = 0.78, n = 312 | | | |
| 114 | R. Mersey | 2016 | 0 | - | - | - | - | - | - | - | - | - | - | - | - | - | - | - | - |
| 115 | R. Ribble | 1147 | 12 | 4.52 | 5.07 | **0.0004** | ↘↗↘ | - | - | - | - | 3.43 | 17.35 | **0.002** | ↗↘ | 2.76 | 4.16 | **0.01** | ↘↗ |
| 116 | R. Wyre | 308 | 2 | 1.00 | 3.86 | **0.05** | ↗ | - | - | - | - | 1.17 | 2.98 | 0.16 | - | 4.31 | 17.78 | **<0.0001** | ↘↗ |
| 117 | R. Lune | 1007 | 26 | 5.39 | 7.76 | **<0.0001** | →↗↘ | 4.09 | 33.20 | **<0.0001** | →↗↘ | 3.25 | 40.70 | **<0.0001** | →↘ | 2.02 | 6.30 | **0.002** | ↘ |
| 118 | R. Keer | 56 | 2 | 1.00 | 0.50 | 0.48 | - | - | - | - | - | - | - | - | - | - | - | - | - |
| 119 | R. Bela | 140 | 0 | - | - | - | - | - | - | - | - | - | - | - | - | - | - | - | - |
| 120 | R. Kent | 225 | 2 | 1.00 | 0.07 | 0.79 | - | 1.77 | 2.17 | 0.36 | | 2.53 | 17.28 | **0.0007** | ↘ | 3.01 | 6.96 | **0.0002** | ↘↗ |
| 121 | R. Gilpin | 60 | 0 | - | - | - | - | - | - | - | - | - | - | - | - | - | - | - | - |
| 122 | R. Winster | 55 | 0 | - | - | - | - | - | - | - | - | - | - | - | - | - | - | - | - |
| 123 | R. Eea | 31 | 0 | - | - | - | - | - | - | - | - | - | - | - | - | - | - | - | - |

| | | | | | | | | | | | | | | | | | | | |
|---|---|---|---|---|---|---|---|---|---|---|---|---|---|---|---|---|---|---|---|
| 124 | R. Leven | 256 | 6 | 1.00 | 12.66 | **0.0004** | ↗ | 1.00 | 1.92 | 0.17 | - | 4.74 | 29.86 | **<0.0001** | ↘↗↘ | 3.52 | 6.43 | **0.0001** | ↘↗ |
| 125 | Rusland Pool | 44 | 0 | - | - | - | - | - | - | - | - | - | - | - | - | - | - | - | - |
| 126 | R. Crake | 93 | 1 | 1.00 | 0.00 | 0.99 | - | - | - | - | - | 2.44 | 6.54 | 0.09 | - | - | - | - | - |
| 127 | Kirby Pool | 39 | 0 | - | - | - | - | - | - | - | - | - | - | - | - | - | - | - | - |
| 128 | R. Lickle | 21 | 1 | 2.42 | 2.92 | 0.09 | | - | - | - | - | - | - | - | - | - | - | - | - |
| 129 | R. Duddon | 88 | 1 | 1.00 | 0.06 | 0.81 | - | - | - | - | - | 2.93 | 23.71 | **0.0001** | ↗→ | 1.00 | 1.19 | 0.28 | **-** |
| 130 | Black Beck | 13 | 0 | - | - | - | - | - | - | - | - | - | - | - | - | - | - | - | - |
| 131 | Haverigg Pool | 36 | 0 | - | - | - | - | - | - | - | - | - | - | - | - | - | - | - | - |
| 132 | R. Esk (Cumbria) | 75 | 0 | - | - | - | - | - | - | - | - | 2.38 | 3.63 | 0.27 | - | 1.00 | 6.80 | **0.01** | ↘ |
| 133 | R. Irt | 98 | 0 | - | - | - | - | - | - | - | - | 3.04 | 12.10 | **0.01** | ↘↗↘ | 3.65 | 25.88 | **<0.0001** | ↘→ |
| 134 | R. Calder | 55 | 2 | 1.00 | 0.46 | 0.50 | - | - | - | - | - | 4.22 | 35.72 | **<0.0001** | →↘ | 2.44 | 38.88 | **<0.0001** | ↘ |
| 135 | R. Ehen | 156 | 5 | 2.67 | 4.10 | **0.006** | ↗↘ | - | - | - | - | 2.13 | 3.93 | 0.34 | - | 1.00 | 48.52 | **<0.0001** | ↘ |
| 136 | R. Derwent (Cumbria) | 664 | 9 | 3.81 | 2.77 | **0.03** | ↘↗↘ | - | - | - | - | 3.99 | 58.66 | **<0.0001** | ↗↘ | 3.39 | 6.16 | **0.0005** | ↘↗ |
| 137 | R. Ellen | 127 | 0 | - | - | - | - | - | - | - | - | 4.86 | 27.71 | **<0.0001** | ↗↘ | 1.00 | 0.08 | 0.77 | - |
| 138 | R. Waver | 104 | 0 | - | - | - | - | - | - | - | - | - | - | - | - | - | - | - | - |
| 139 | R. Wampool | 154 | 0 | - | - | - | - | - | - | - | - | - | - | - | - | - | - | - | - |
| 140 | R. Eden* | 2298 | 21 | 3.57 | 8.12 | **0.0002** | ↗↘ | 2.99 | 41.71 | **<0.0001** | →↘ | 4.42 | 35.61 | **<0.0001** | ↘↗↘ | 3.78 | 36.16 | **<0.0001** | ↘→ |
| 141 | R. Lyne | 240 | 2 | 1.76 | 0.52 | 0.66 | - | - | - | - | - | - | - | - | - | - | - | - | - |

↘ = Decrease, ↗ = Increase, → = Stable. * Returning adult data from a tributary (River Caldew).

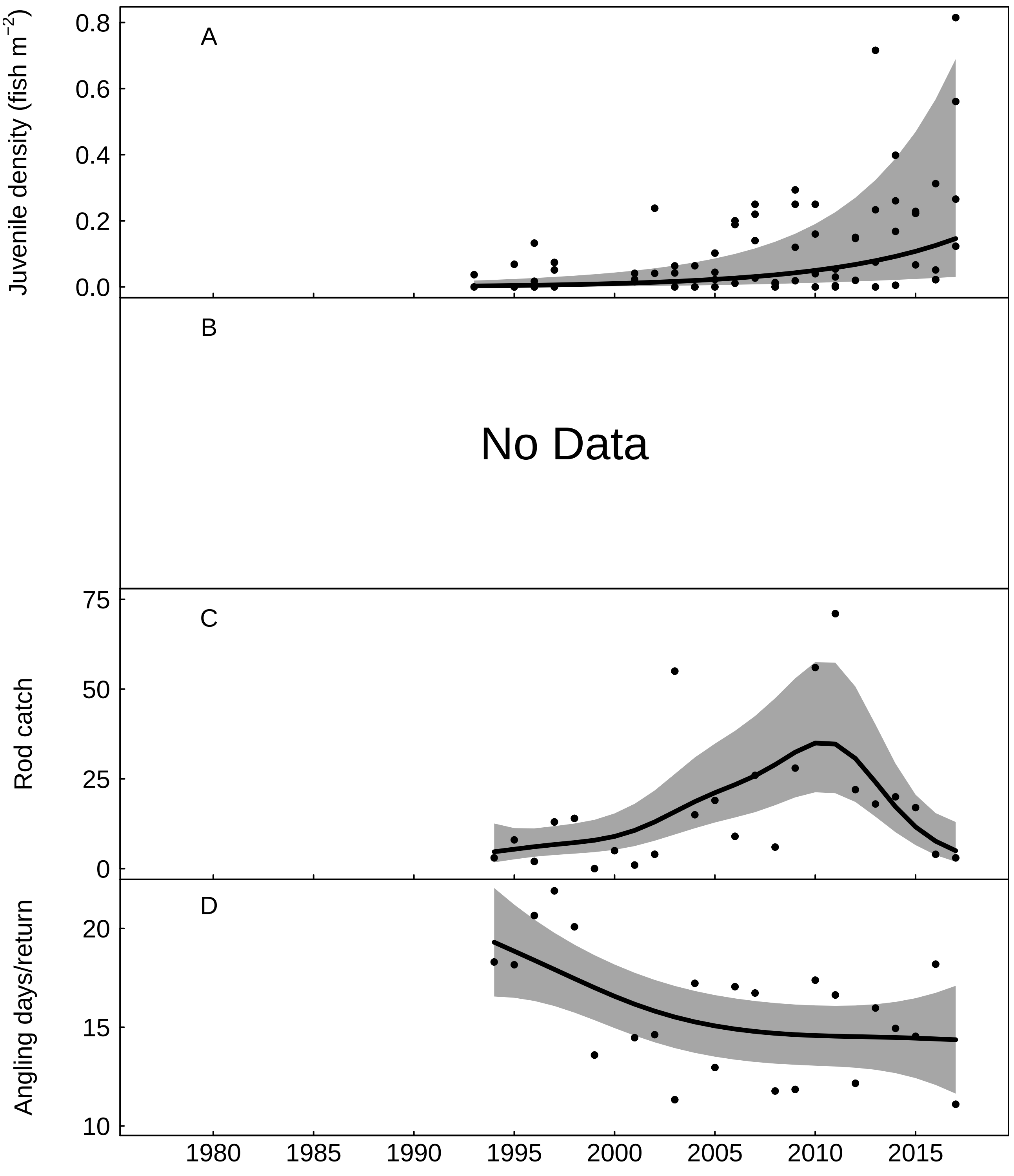


Fig. S1: Trends in the River Aln for A) juvenile salmon density, fitted values from Gaussian additive mixed model, grey areas represent 95% point-wise confidence bands for the smoother, B) No Data, C) rod catch, fitted values from negative binomial generalized additive model, grey areas represent 95% point-wise confidence bands for the smoother, and D) angling effort measured as number of days fished per licence return, fitted values from Gaussian additive model, grey areas represent 95% point-wise confidence bands for the smoother.

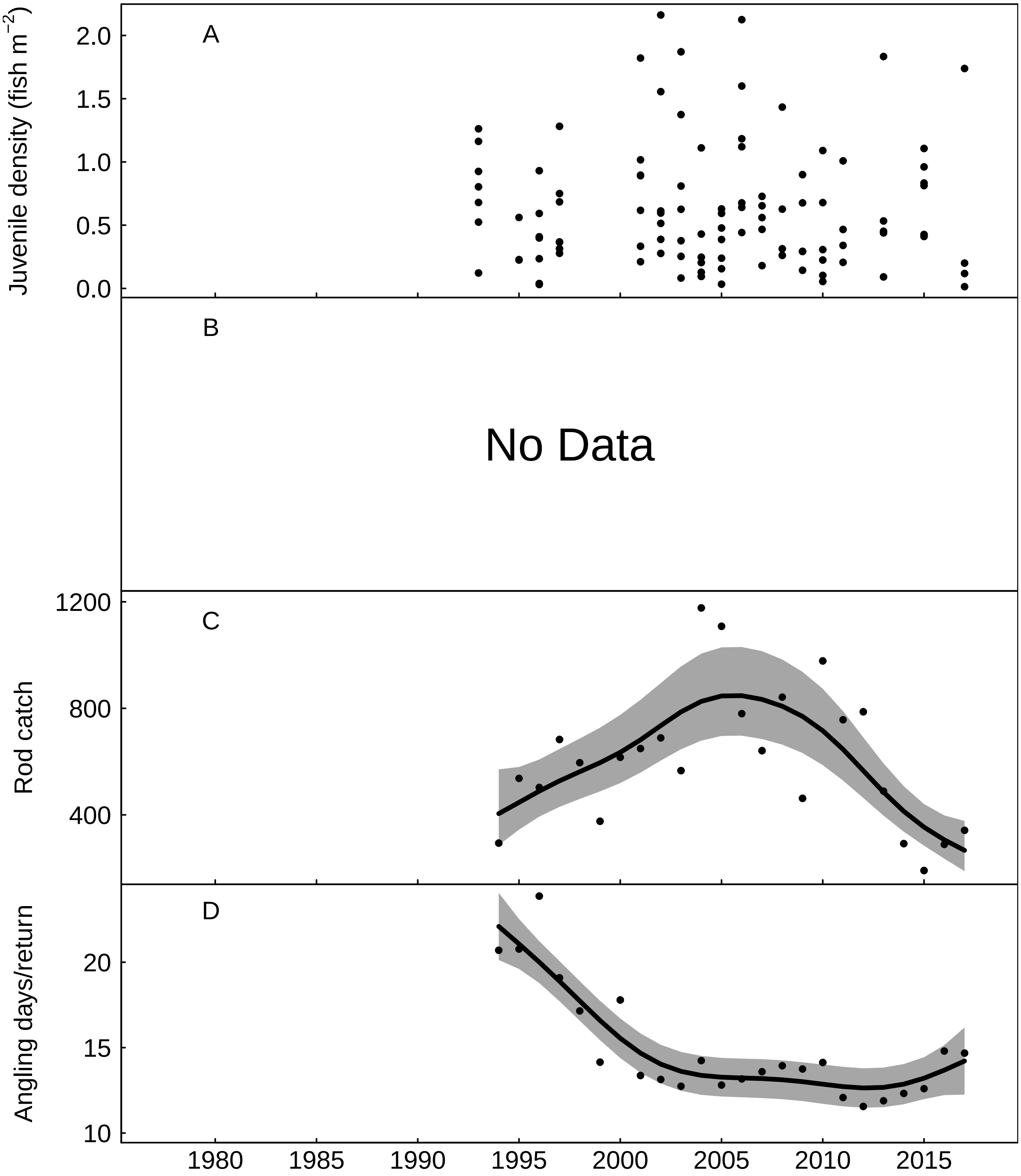


Fig. S2: Trends in the River Coquet for A) juvenile salmon density, B) No Data, C) rod catch, fitted values from negative binomial generalized additive model, grey areas represent 95% point-wise confidence bands for the smoother, and D) angling effort measured as number of days fished per licence return, fitted values from Gaussian additive model, grey areas represent 95% point-wise confidence bands for the smoother.

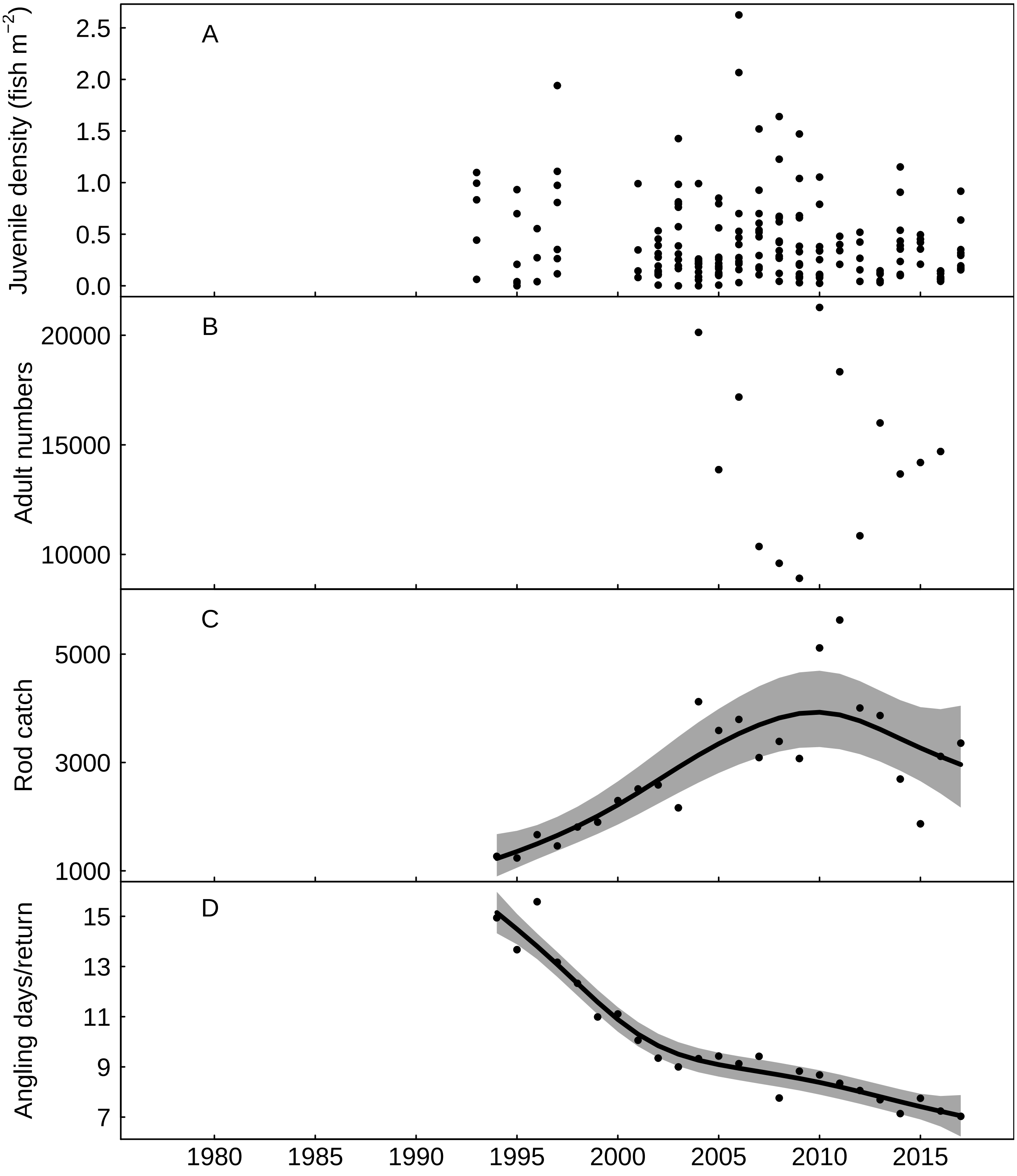


Fig. S3: Trends in the River Tyne for A) juvenile salmon density, B) returning adult numbers, C) rod catch, fitted values from negative binomial generalized additive model, grey areas represent 95% point-wise confidence bands for the smoother, and D) angling effort measured as number of days fished per licence return, fitted values from Gaussian additive model, grey areas represent 95% point-wise confidence bands for the smoother.

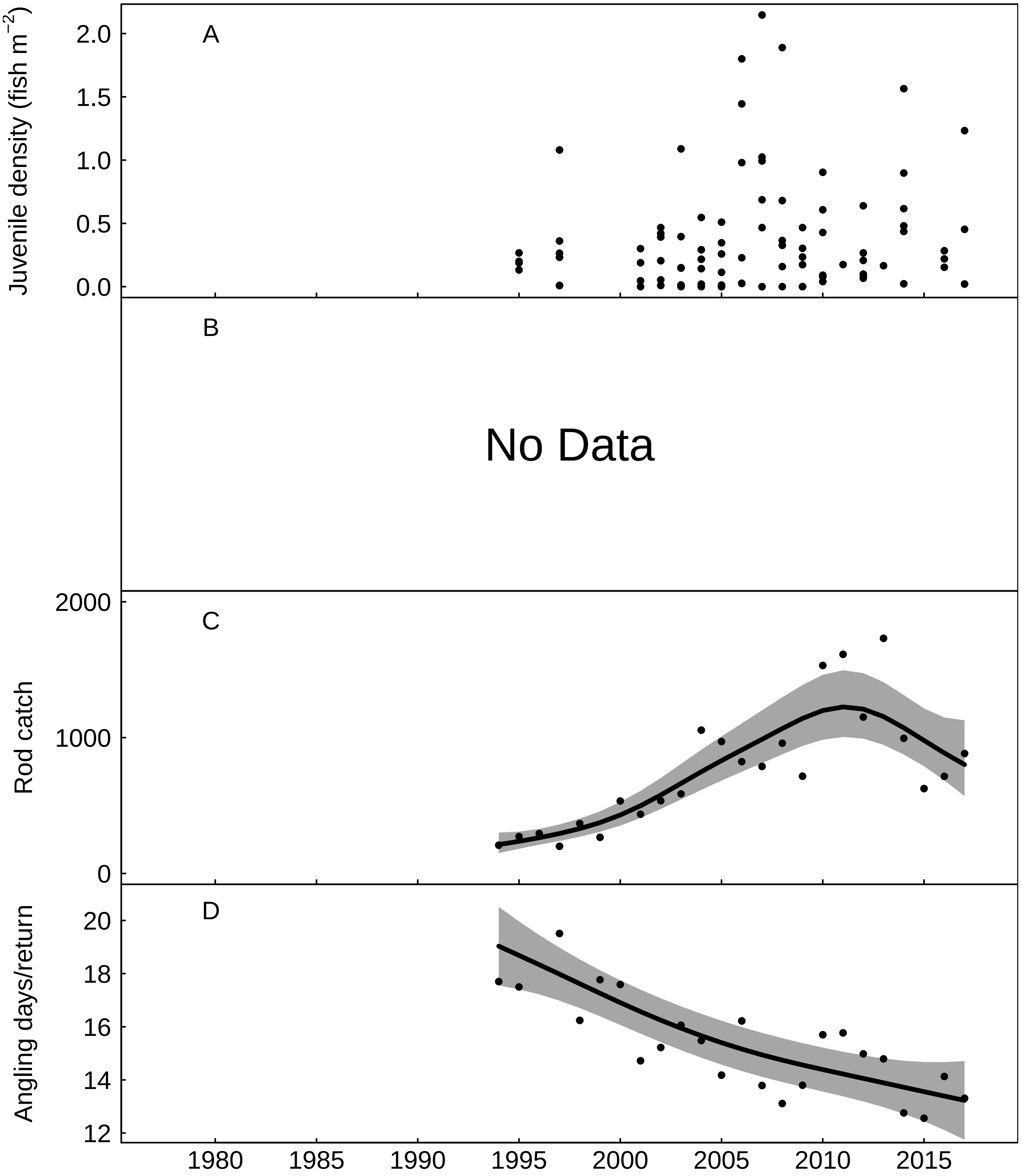


Fig. S4: Trends in the River Wear for A) juvenile salmon density, B) No Data, C) rod catch, fitted values from negative binomial generalized additive model, grey areas represent 95% point-wise confidence bands for the smoother, and D) angling effort measured as number of days fished per licence return, fitted values from Gaussian additive model, grey areas represent 95% point-wise confidence bands for the smoother.

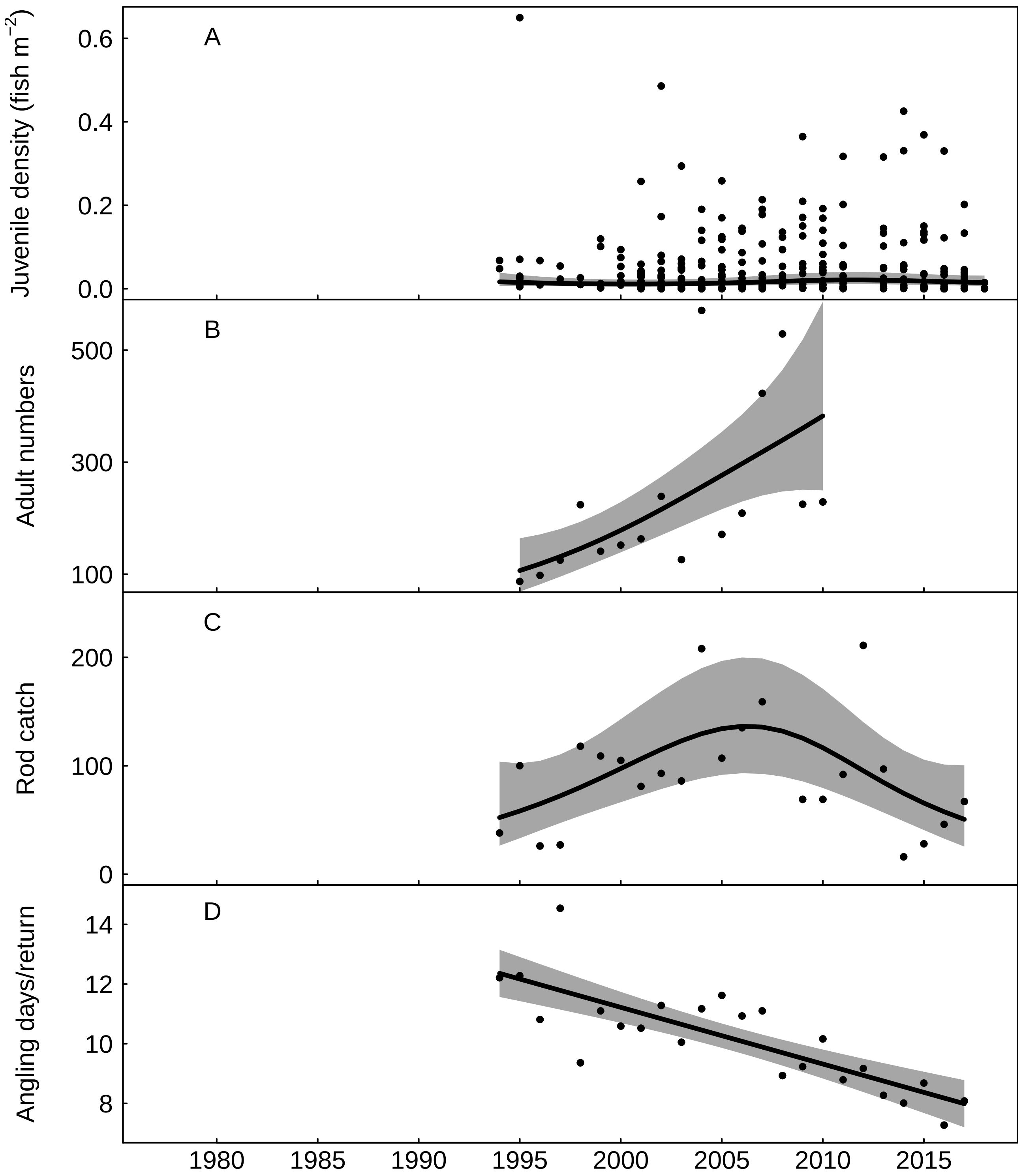


Fig. S5: Trends in the River Tees for A) juvenile salmon density, fitted values from Gaussian additive mixed model, grey areas represent 95% point-wise confidence bands for the smoother, B) returning adult numbers, fitted values from negative binomial generalized additive model, grey areas represent 95% point-wise confidence bands for the smoother, C) rod catch, fitted values from negative binomial generalized additive model, grey areas represent 95% point-wise confidence bands for the smoother, and D) angling effort measured as number of days fished per licence return, fitted values from Gaussian additive model, grey areas represent 95% point-wise confidence bands for the smoother.

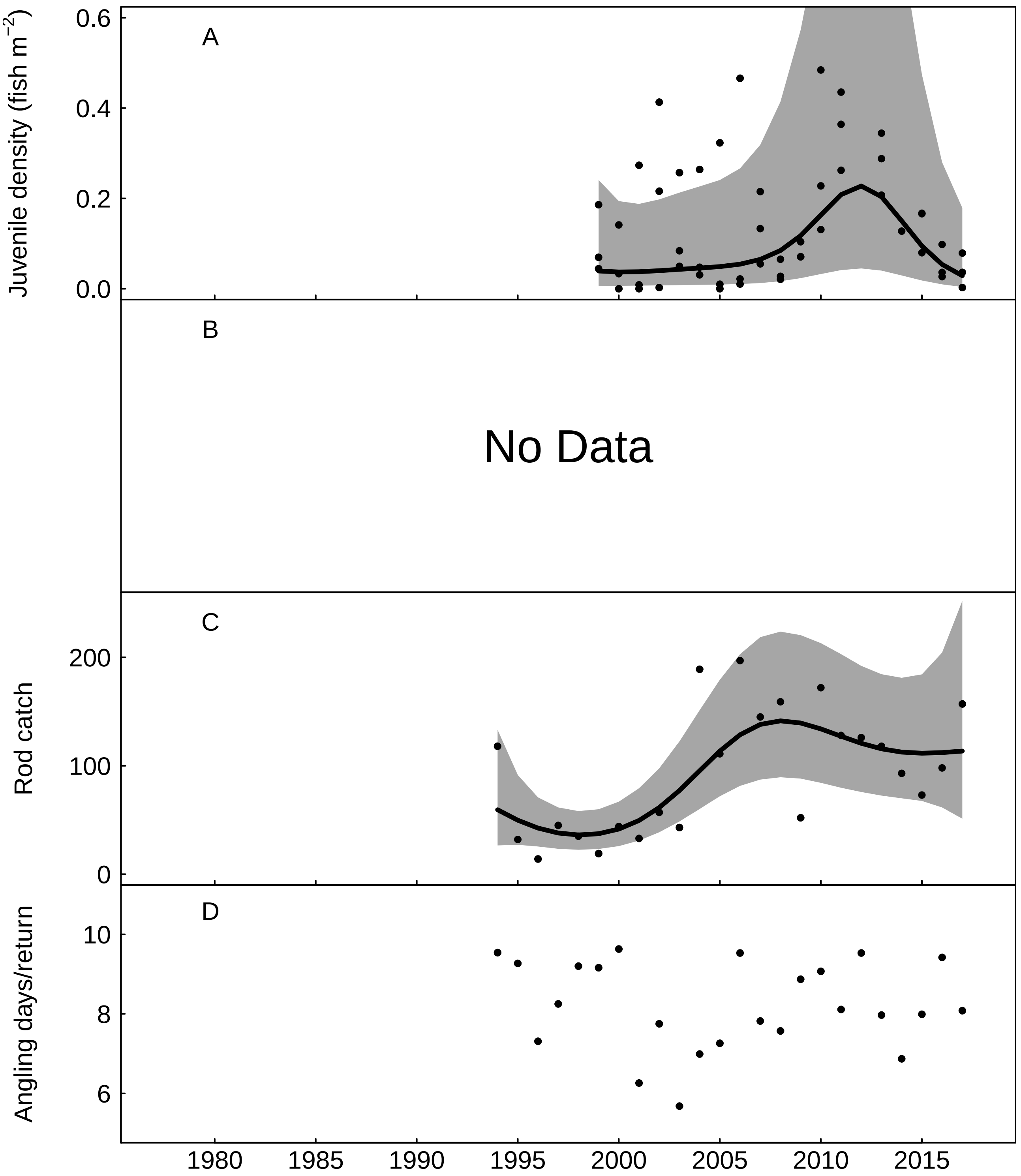


Fig. S6: Trends in the Yorkshire Esk for A) juvenile salmon density, fitted values from Gaussian additive mixed model, grey areas represent 95% point-wise confidence bands for the smoother, B) No Data, C) rod catch, fitted values from negative binomial generalized additive model, grey areas represent 95% point-wise confidence bands for the smoother, and D) angling effort measured as number of days fished per licence return.

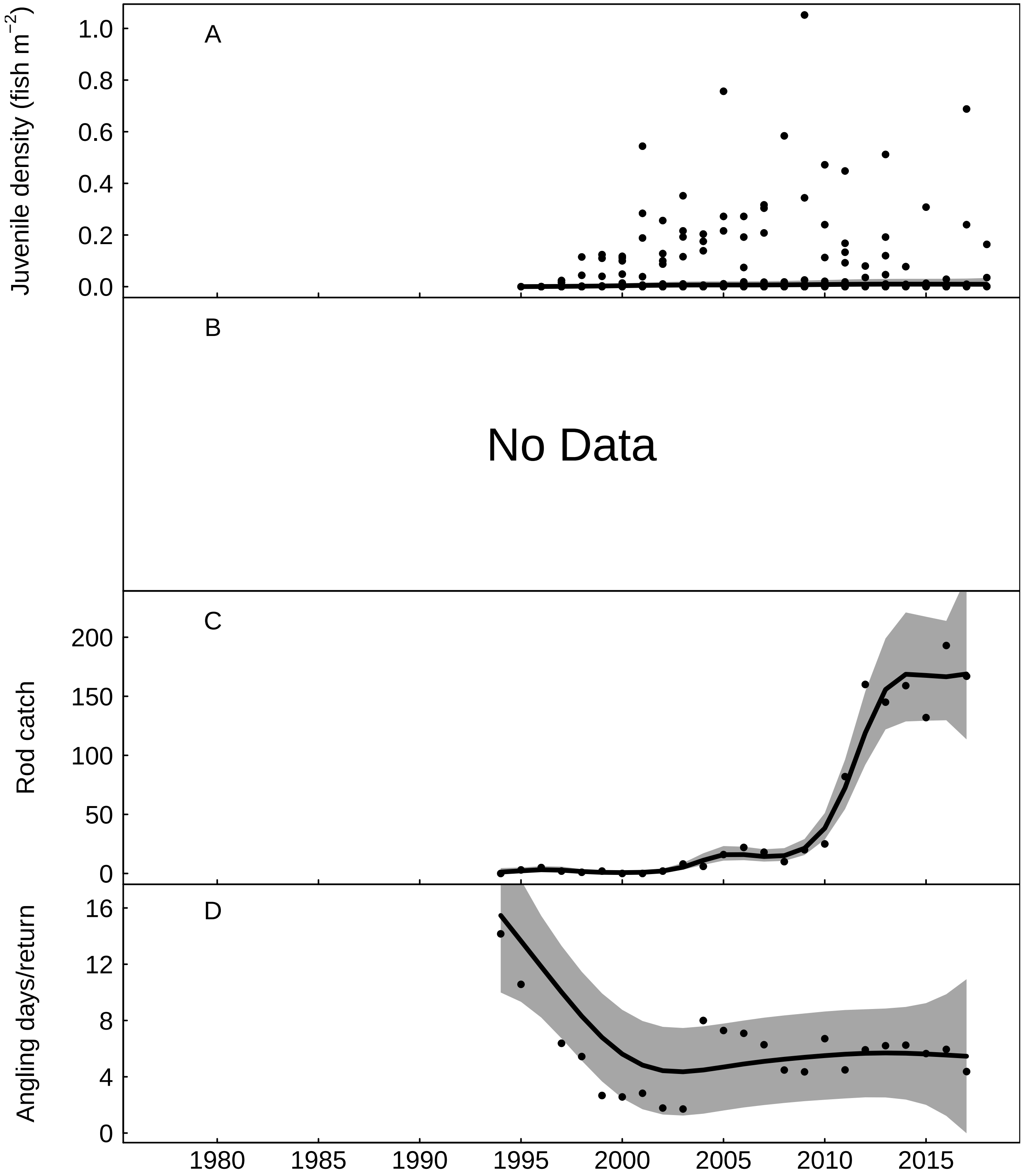


Fig. S7: Trends in the Upper Ouse for A) juvenile salmon density, fitted values from Gaussian additive mixed model, grey areas represent 95% point-wise confidence bands for the smoother, B) No Data, C) rod catch, fitted values from negative binomial generalized additive model, grey areas represent 95% point-wise confidence bands for the smoother, and D) angling effort measured as number of days fished per licence return, fitted values from Gaussian additive model, grey areas represent 95% point-wise confidence bands for the smoother.

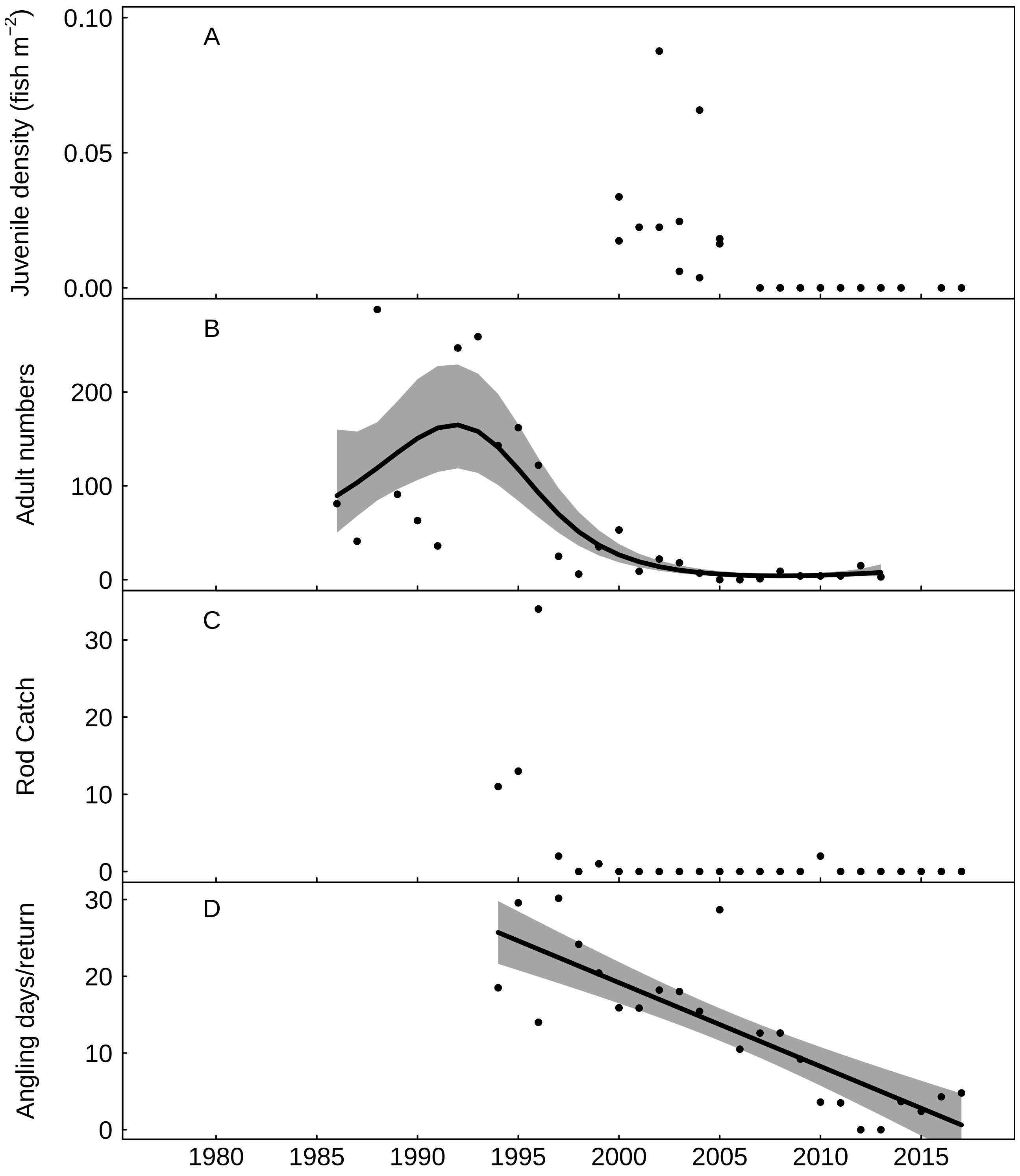


Fig. S8: Trends in the River Thames for A) juvenile salmon density, B) returning adult numbers, fitted values from negative binomial generalized additive model, grey areas represent 95% point-wise confidence bands for the smoother, C) rod catch, and D) angling effort measured as number of days fished per licence return, fitted values from Gaussian additive model, grey areas represent 95% point-wise confidence bands for the smoother.

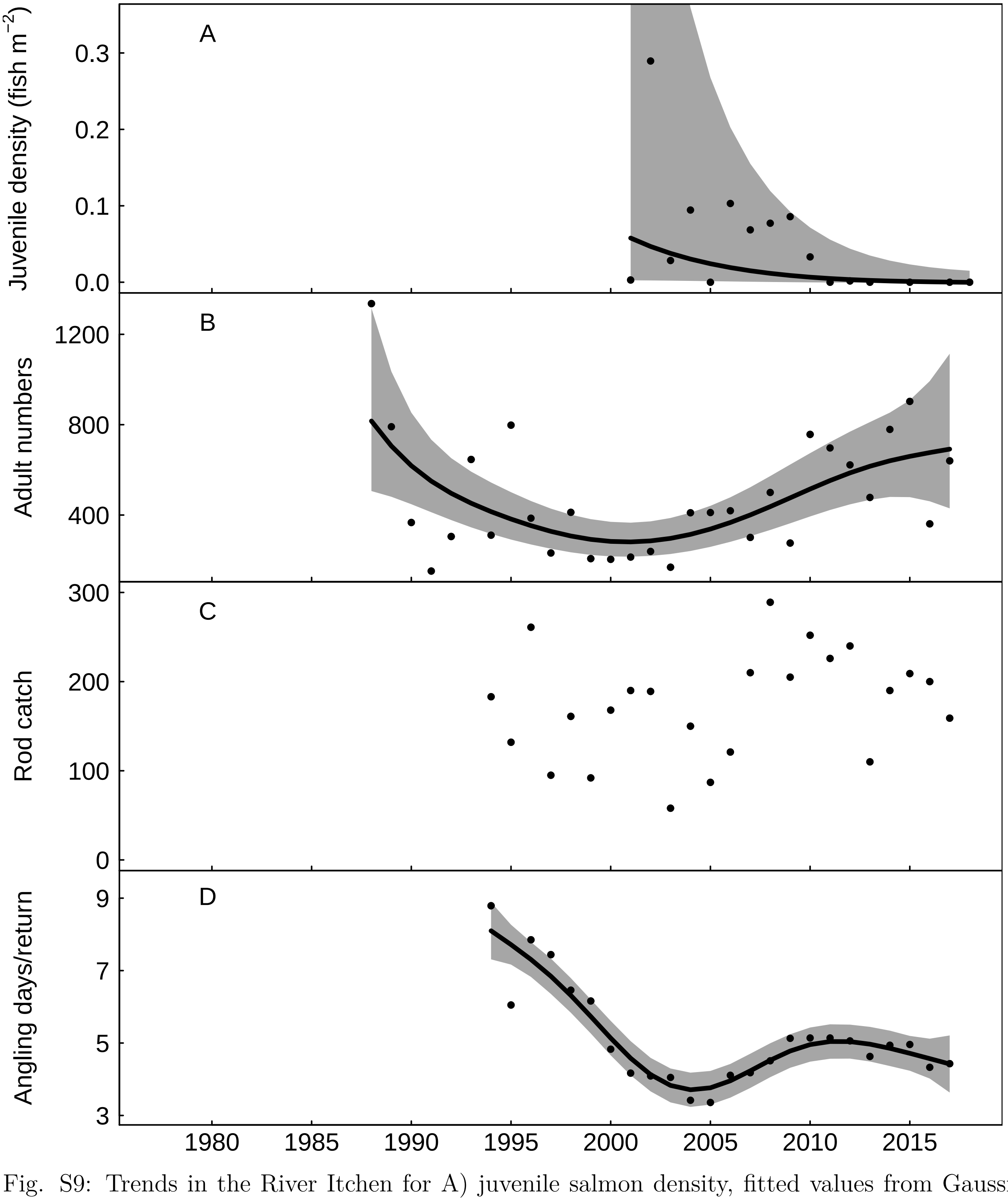


Fig. S9: Trends in the River Itchen for A) juvenile salmon density, fitted values from Gaussian additive mixed model, grey areas represent 95% point-wise confidence bands for the smoother, B) returning adult numbers, fitted values from negative binomial generalized additive model, grey areas represent 95% point-wise confidence bands for the smoother, C) rod catch, and D) angling effort measured as number of days fished per licence return, fitted values from Gaussian additive model, grey areas represent 95% point-wise confidence bands for the smoother.

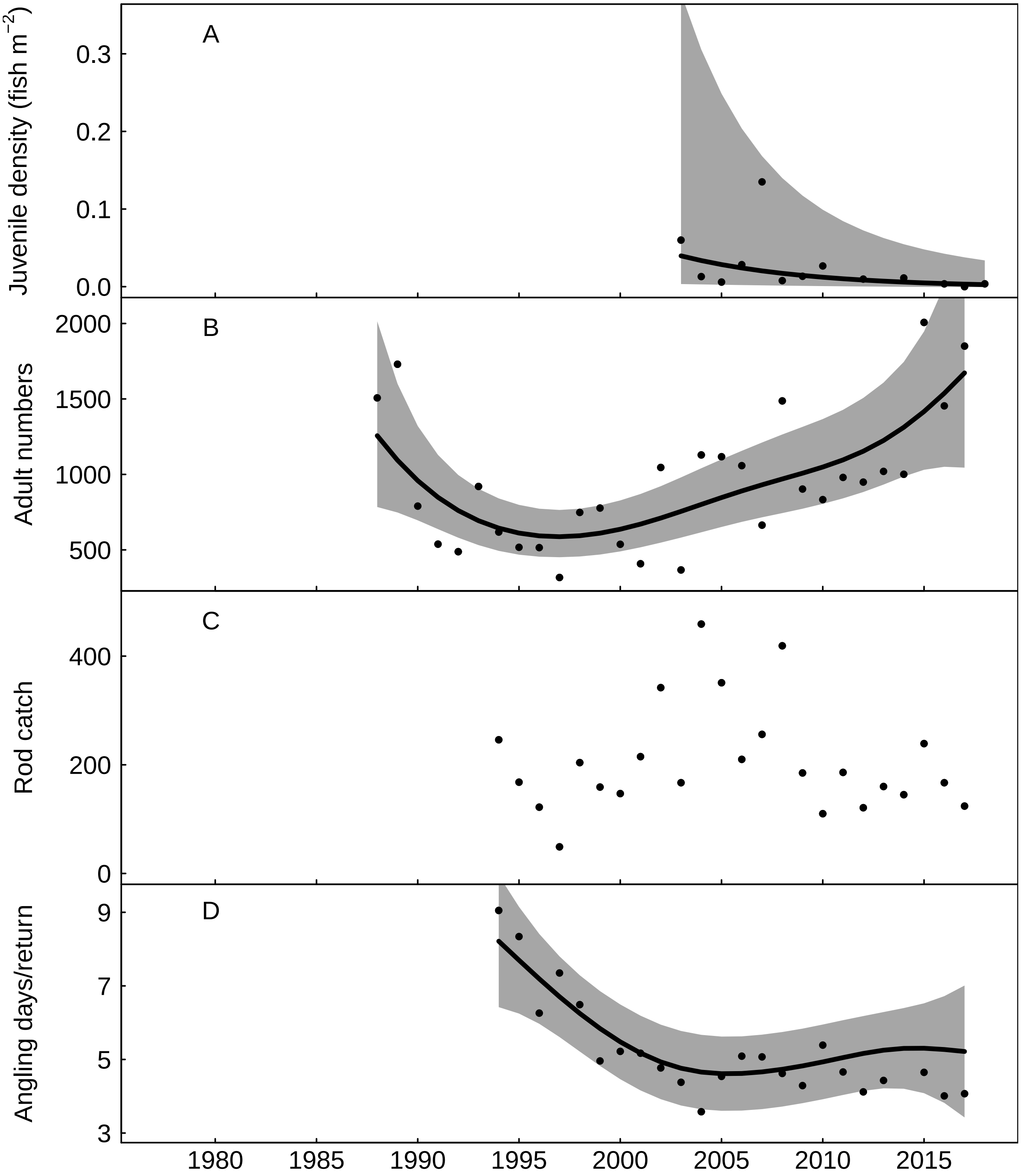


Fig. S10: Trends in the River Test for A) juvenile salmon density, fitted values from Gaussian additive mixed model, grey areas represent 95% point-wise confidence bands for the smoother, B) returning adult numbers, fitted values from negative binomial generalized additive model, grey areas represent 95% point-wise confidence bands for the smoother, C) rod catch, and D) angling effort measured as number of days fished per licence return, fitted values from Gaussian additive model, grey areas represent 95% point-wise confidence bands for the smoother.

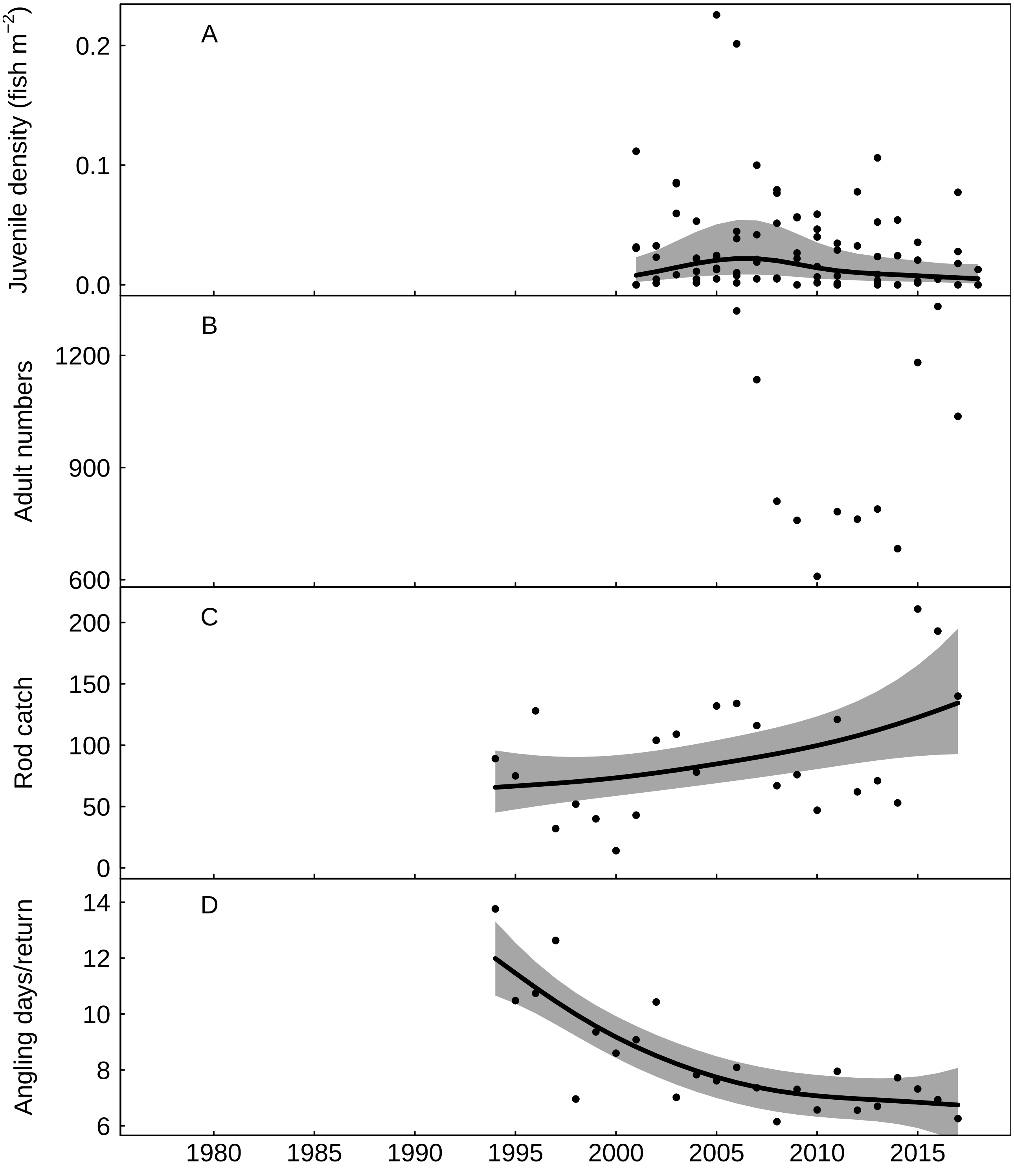


Fig. S11: Trends in the Hampshire Avon for A) juvenile salmon density, fitted values from Gaussian additive mixed model, grey areas represent 95% point-wise confidence bands for the smoother, B) returning adult numbers, C) rod catch, fitted values from negative binomial generalized additive model, grey areas represent 95% point-wise confidence bands for the smoother, and D) angling effort measured as number of days fished per licence return, fitted values from Gaussian additive model, grey areas represent 95% point-wise confidence bands for the smoother.

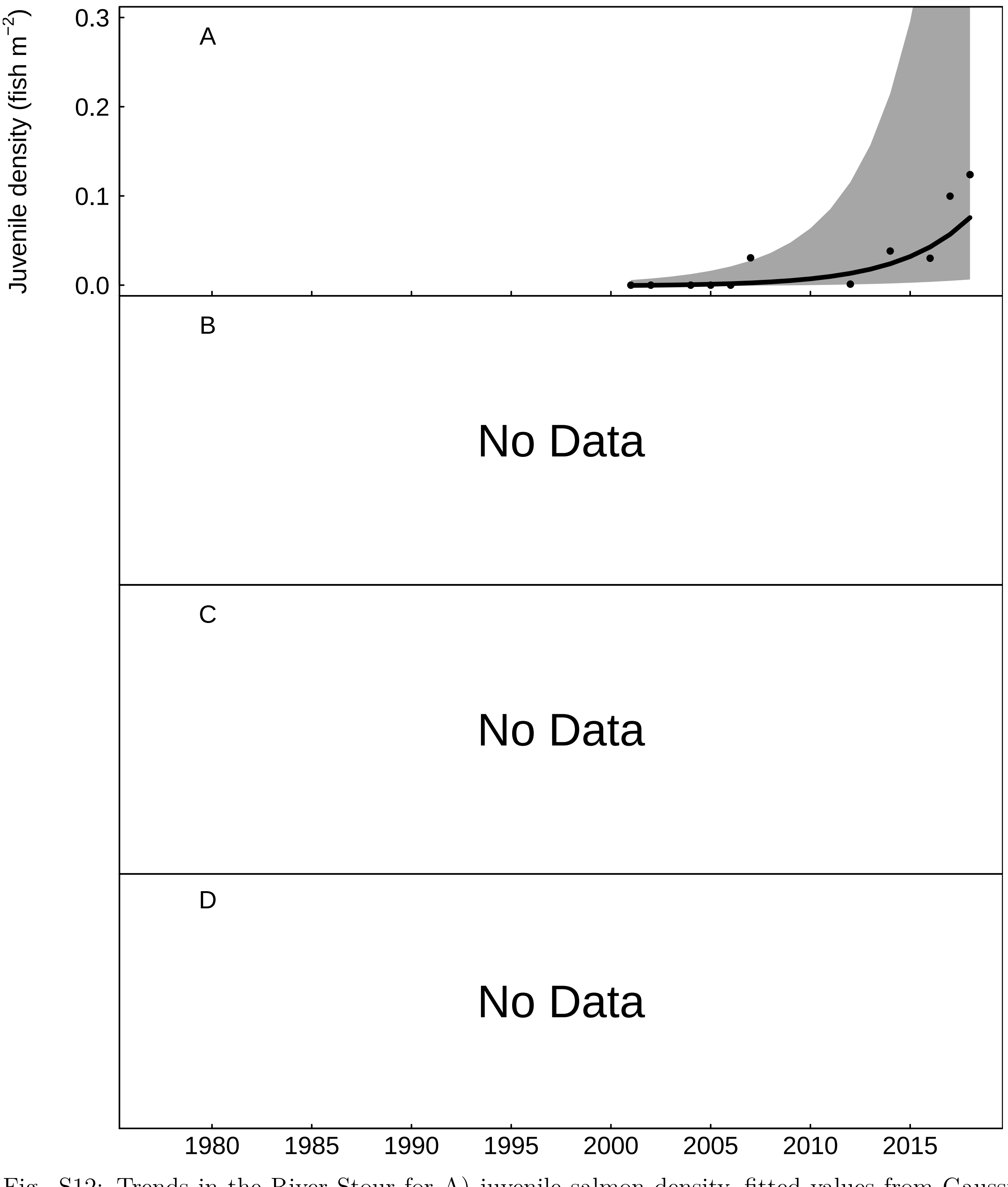


Fig. S12: Trends in the River Stour for A) juvenile salmon density, fitted values from Gaussian additive mixed model, grey areas represent 95% point-wise confidence bands for the smoother, B) No Data, C) No Data, and D) No Data.

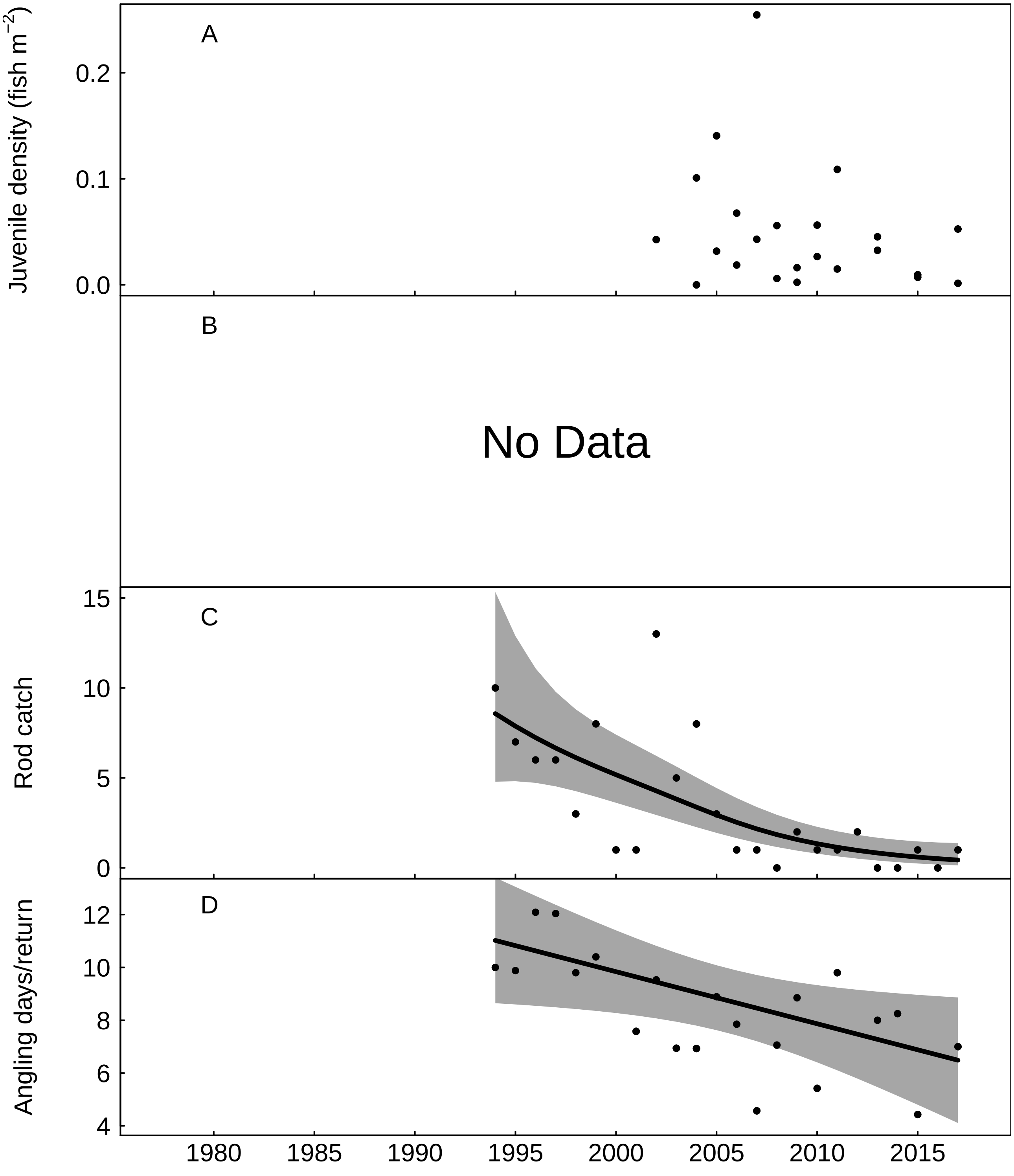


Fig. S13: Trends in the River Piddle for A) juvenile salmon density, B) No Data, C) rod catch, fitted values from negative binomial generalized additive model, grey areas represent 95% point-wise confidence bands for the smoother, and D) angling effort measured as number of days fished per licence return, fitted values from Gaussian additive model, grey areas represent 95% point-wise confidence bands for the smoother.

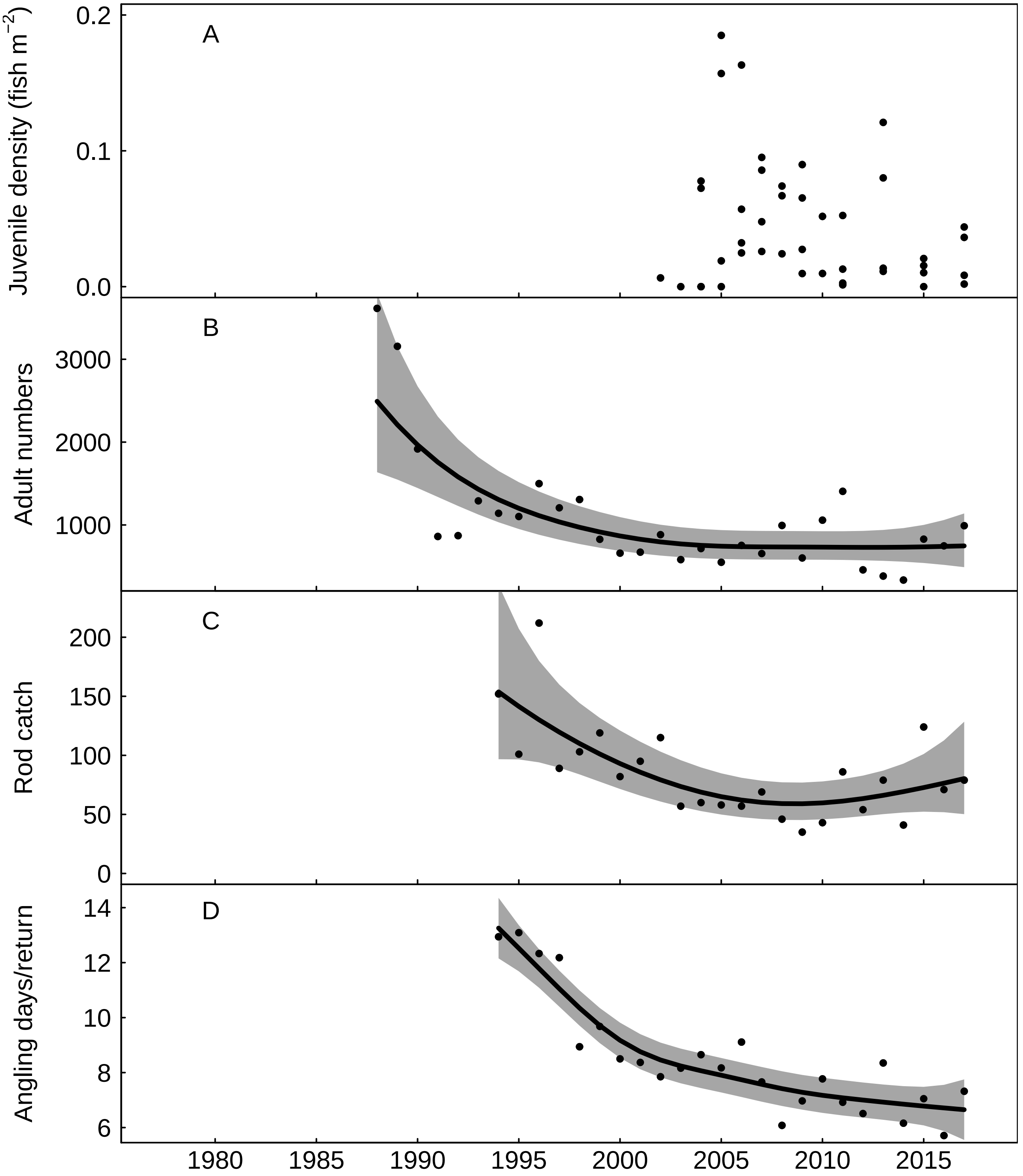


Fig. S14: Trends in the River Frome for A) juvenile salmon density, B) returning adult numbers, fitted values from negative binomial generalized additive model, grey areas represent 95% point-wise confidence bands for the smoother, C) rod catch, fitted values from negative binomial generalized additive model, grey areas represent 95% point-wise confidence bands for the smoother, and D) angling effort measured as number of days fished per licence return, fitted values from Gaussian additive model, grey areas represent 95% point-wise confidence bands for the smoother.

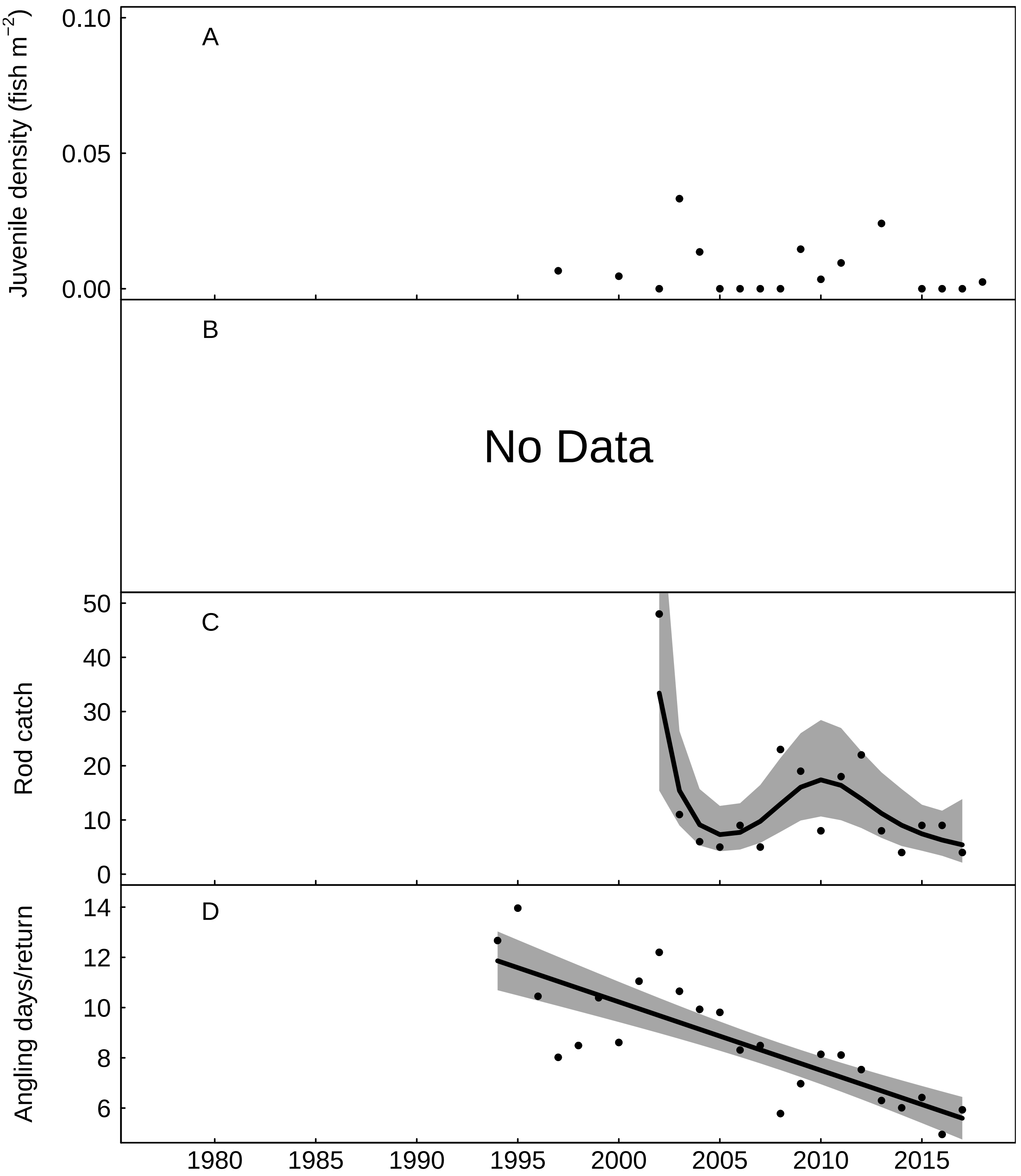


Fig. S15: Trends in the River Axe for A) juvenile salmon density, B) No Data, C) rod catch, fitted values from negative binomial generalized additive model, grey areas represent 95% point-wise confidence bands for the smoother, and D) angling effort measured as number of days fished per licence return, fitted values from Gaussian additive model, grey areas represent 95% point-wise confidence bands for the smoother.

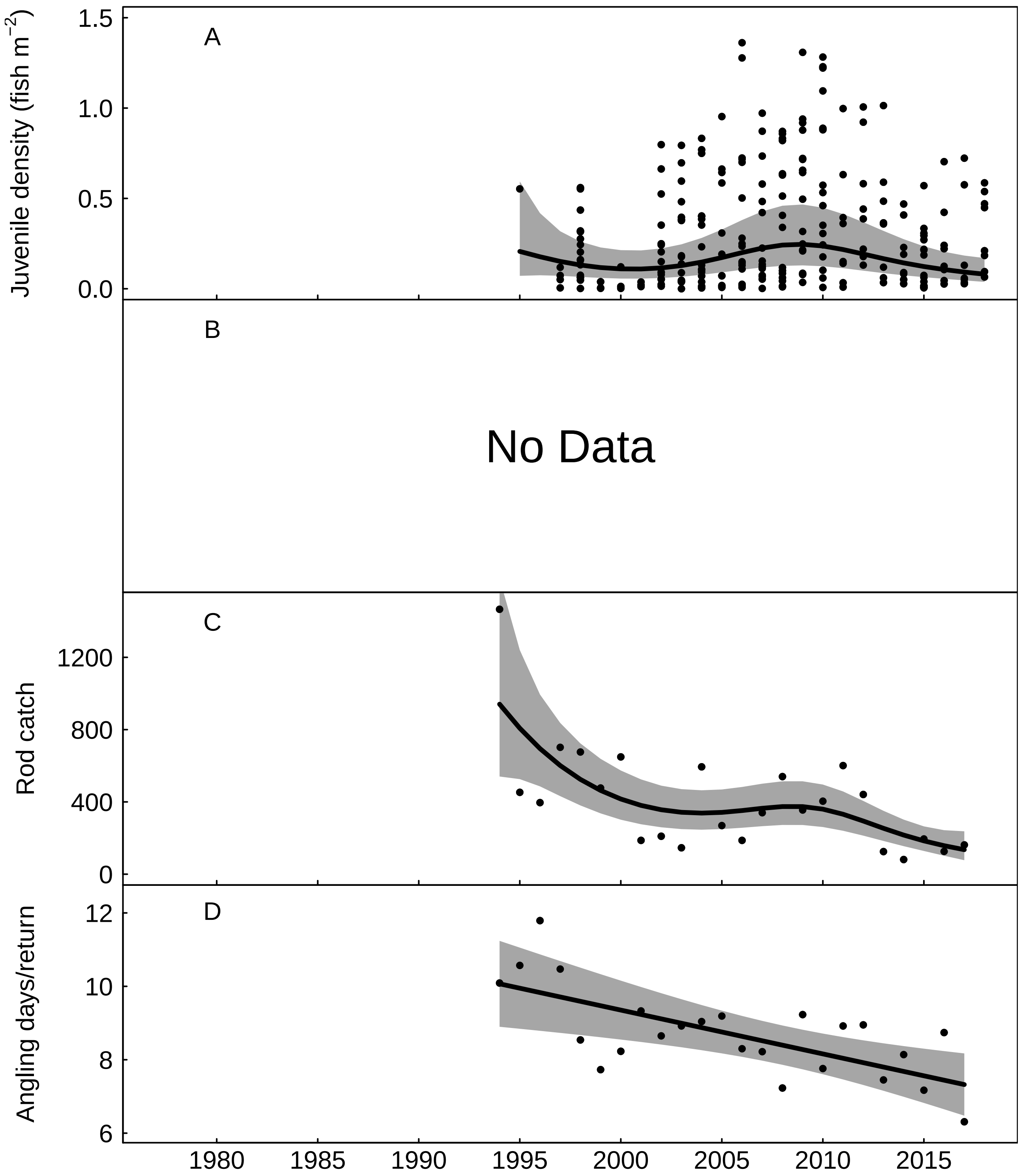


Fig. S16: Trends in the River Exe for A) juvenile salmon density, fitted values from Gaussian additive mixed model, grey areas represent 95% point-wise confidence bands for the smoother, B) No Data, C) rod catch, fitted values from negative binomial generalized additive model, grey areas represent 95% point-wise confidence bands for the smoother, and D) angling effort measured as number of days fished per licence return, fitted values from Gaussian additive model, grey areas represent 95% point-wise confidence bands for the smoother.

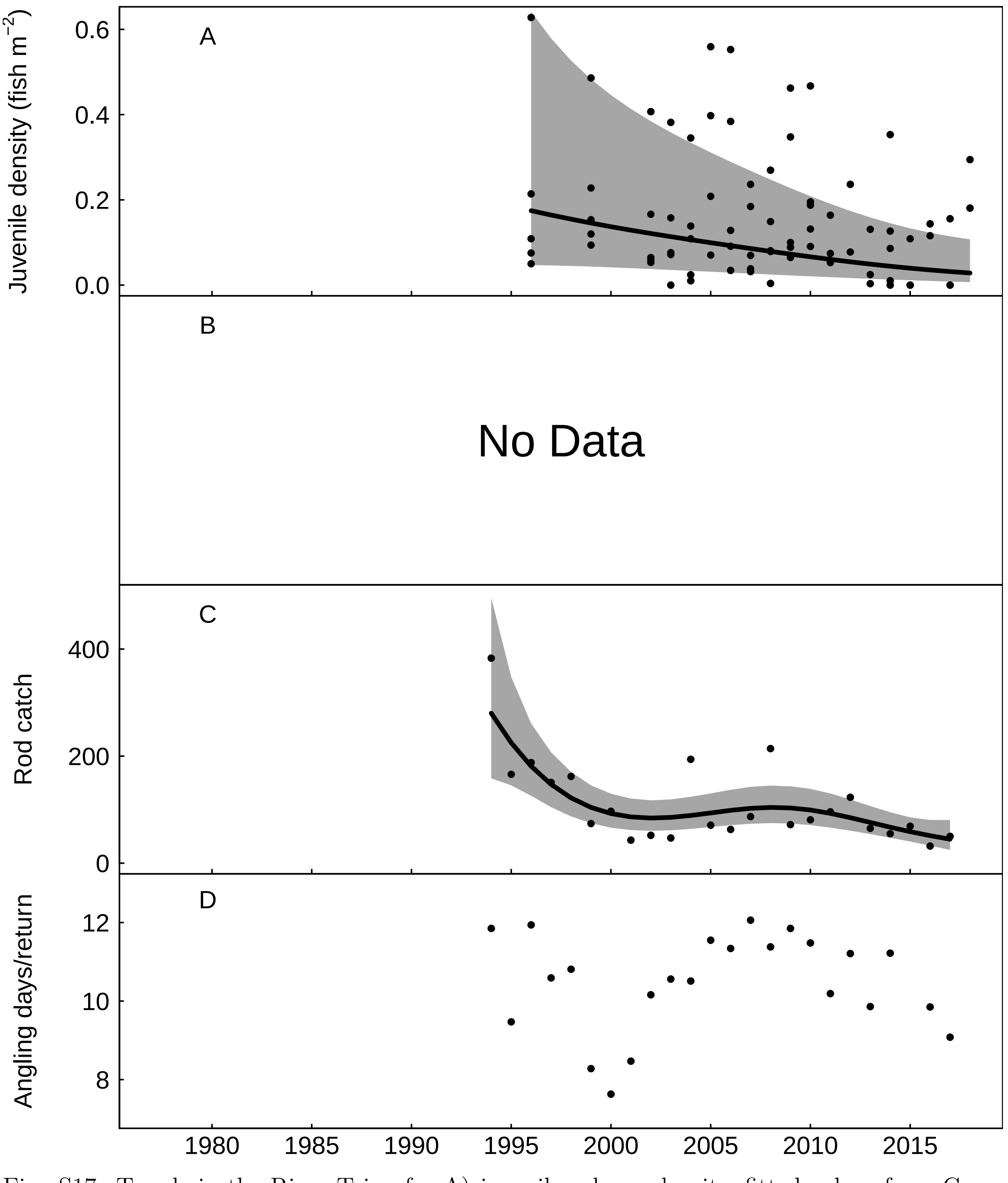


Fig. S17: Trends in the River Teign for A) juvenile salmon density, fitted values from Gaussian additive mixed model, grey areas represent 95% point-wise confidence bands for the smoother, B) No Data, C) rod catch, fitted values from negative binomial generalized additive model, grey areas represent 95% point-wise confidence bands for the smoother, and D) angling effort measured as number of days fished per licence return.

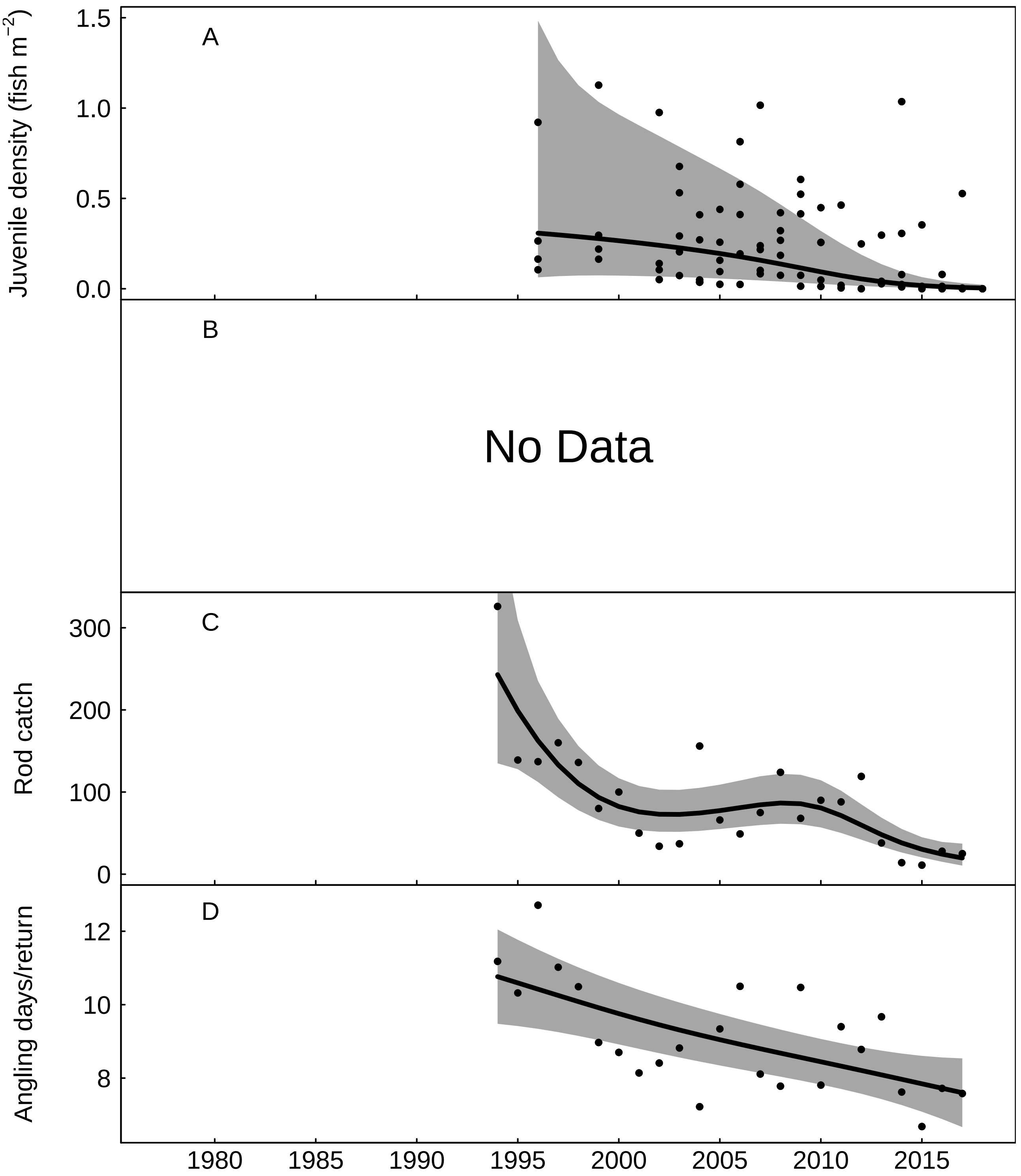


Fig. S18: Trends in the River Dart for A) juvenile salmon density, fitted values from Gaussian additive mixed model, grey areas represent 95% point-wise confidence bands for the smoother, B) No Data, C) rod catch, fitted values from negative binomial generalized additive model, grey areas represent 95% point-wise confidence bands for the smoother, and D) angling effort measured as number of days fished per licence return, fitted values from Gaussian additive model, grey areas represent 95% point-wise confidence bands for the smoother.

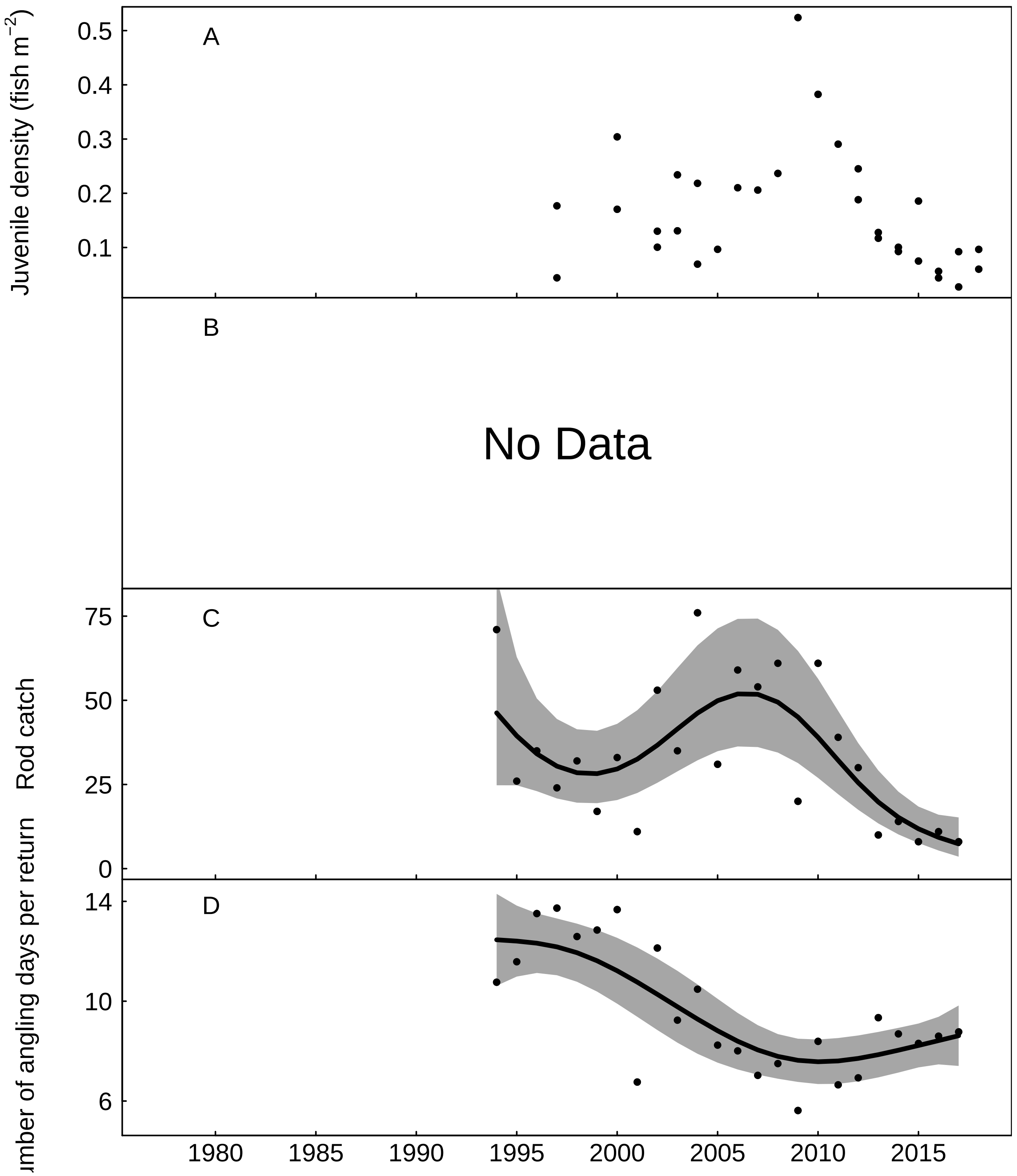


Fig. S19: Trends in the River Avon (Devon) for A) juvenile salmon density, B) No Data, C) rod catch, fitted values from negative binomial generalized additive model, grey areas represent 95% point-wise confidence bands for the smoother, and D) angling effort measured as number of days fished per licence return, fitted values from Gaussian additive model, grey areas represent 95% point-wise confidence bands for the smoother.

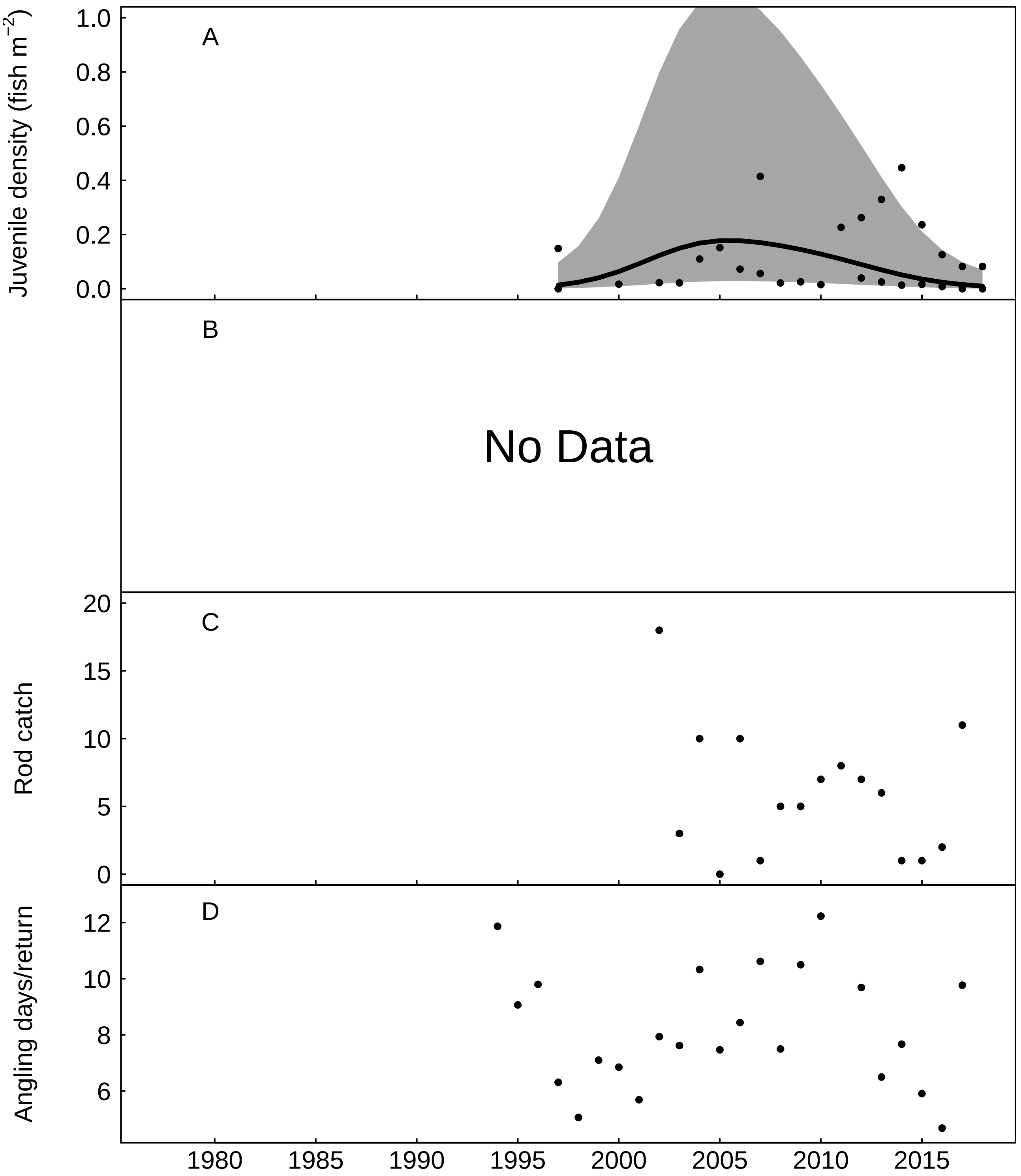


Fig. S20: Trends in the River Erme for A) juvenile salmon density, fitted values from Gaussian additive mixed model, grey areas represent 95% point-wise confidence bands for the smoother, B) No Data, C) rod catch, and D) angling effort measured as number of days fished per licence return.

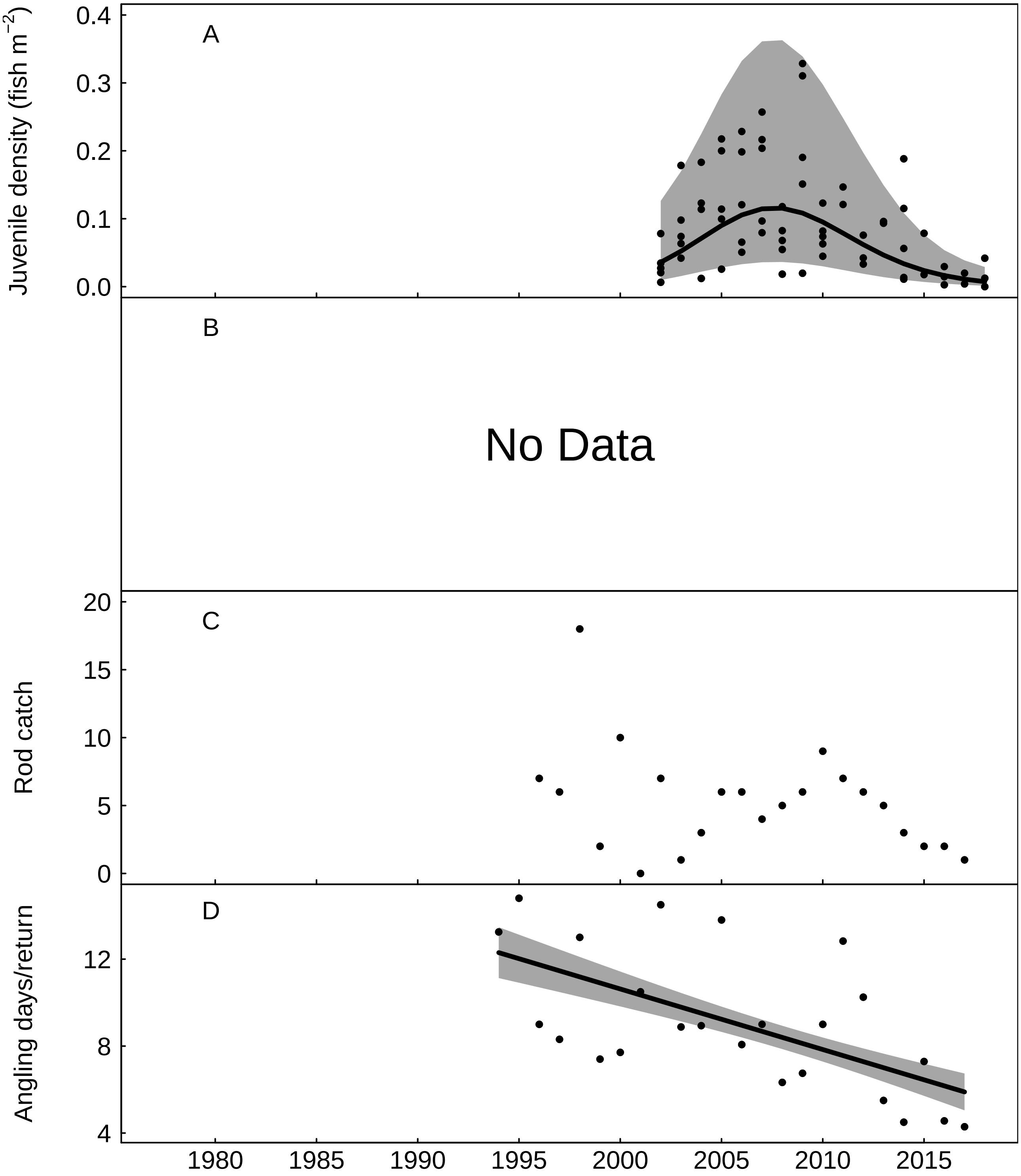


Fig. S21: Trends in the River Yealm for A) juvenile salmon density, fitted values from Gaussian additive mixed model, grey areas represent 95% point-wise confidence bands for the smoother, B) No Data, C) rod catch, and D) angling effort measured as number of days fished per licence return, fitted values from Gaussian additive model, grey areas represent 95% point-wise confidence bands for the smoother.

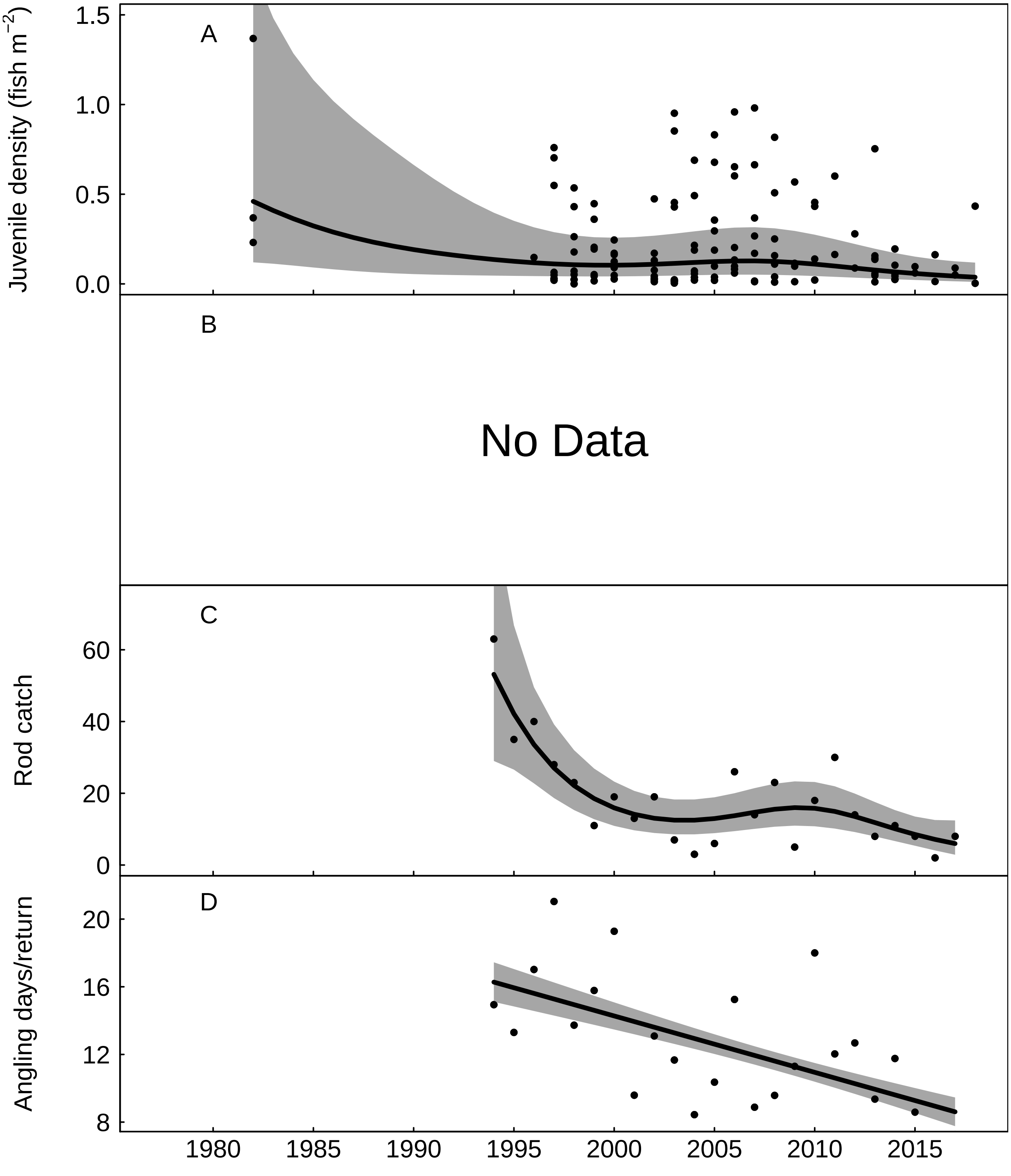


Fig. S22: Trends in the River Plym for A) juvenile salmon density, fitted values from Gaussian additive mixed model, grey areas represent 95% point-wise confidence bands for the smoother, B) No Data, C) rod catch, fitted values from negative binomial generalized additive model, grey areas represent 95% point-wise confidence bands for the smoother, and D) angling effort measured as number of days fished per licence return, fitted values from Gaussian additive model, grey areas represent 95% point-wise confidence bands for the smoother.

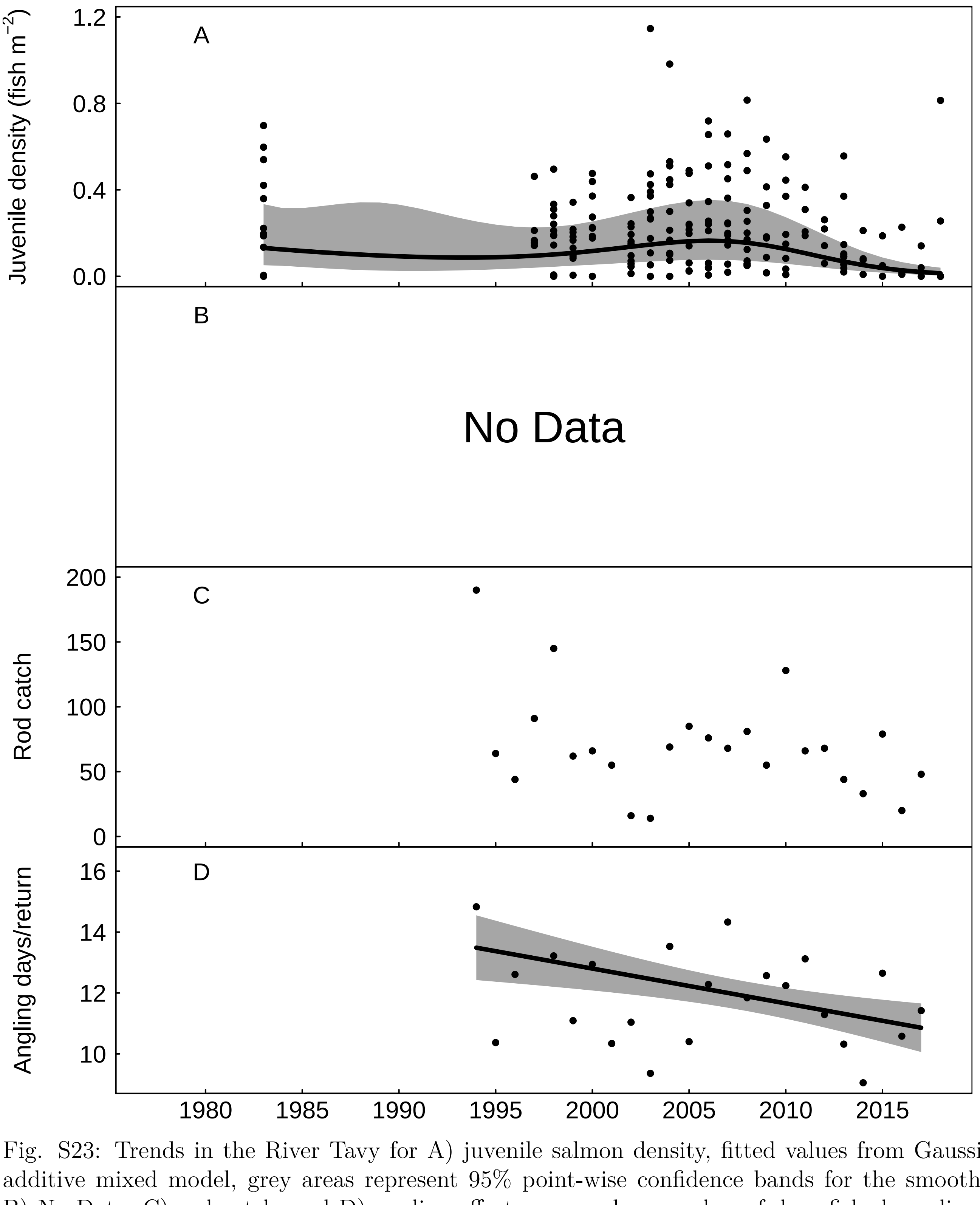


Fig. S23: Trends in the River Tavy for A) juvenile salmon density, fitted values from Gaussian additive mixed model, grey areas represent 95% point-wise confidence bands for the smoother, B) No Data, C) rod catch, and D) angling effort measured as number of days fished per licence return, fitted values from Gaussian additive model, grey areas represent 95% point-wise confidence bands for the smoother.

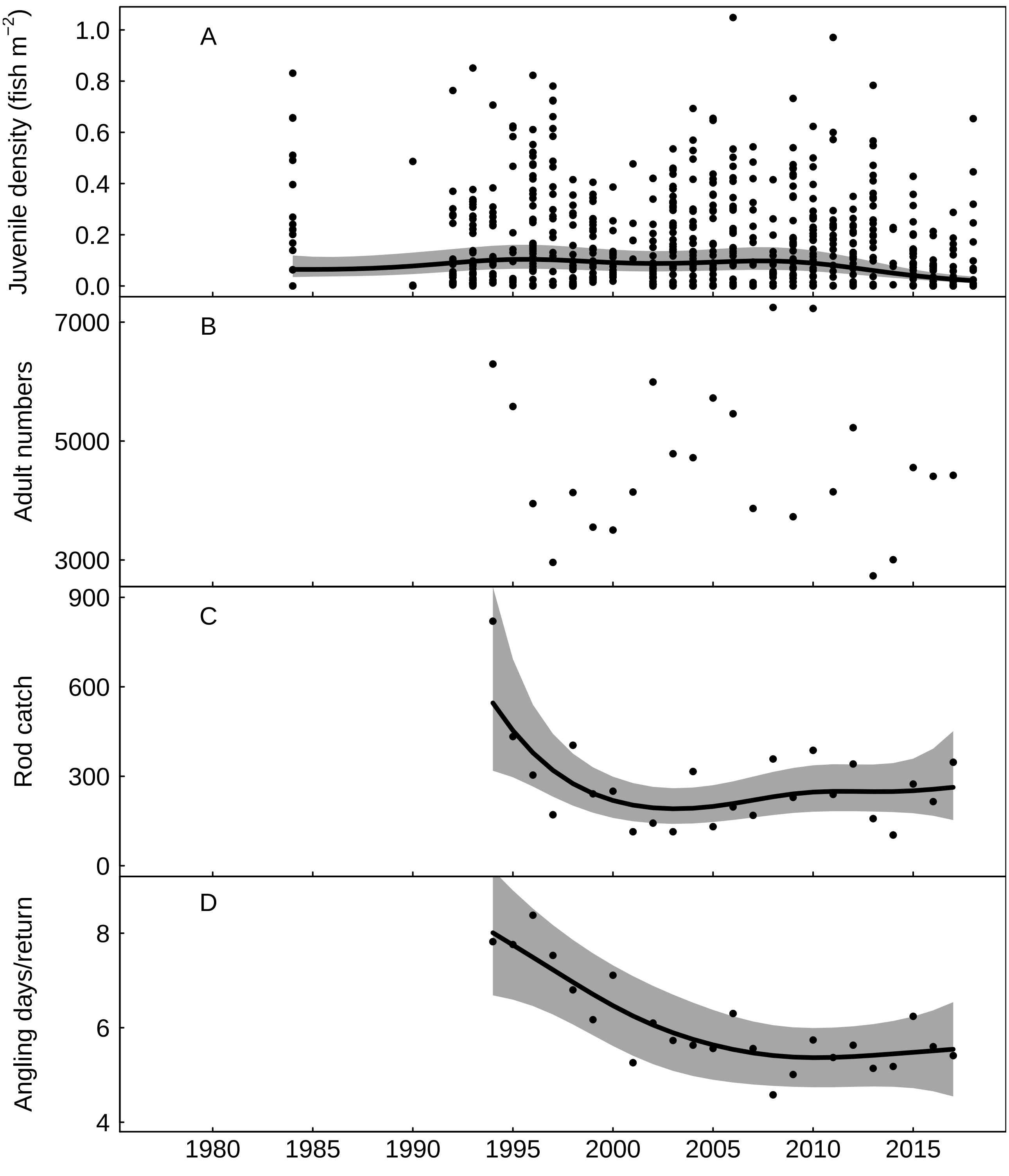


Fig. S24: Trends in the River Tamar for A) juvenile salmon density, fitted values from Gaussian additive mixed model, grey areas represent 95% point-wise confidence bands for the smoother, B) returning adult numbers, C) rod catch, fitted values from negative binomial generalized additive model, grey areas represent 95% point-wise confidence bands for the smoother, and D) angling effort measured as number of days fished per licence return, fitted values from Gaussian additive model, grey areas represent 95% point-wise confidence bands for the smoother.

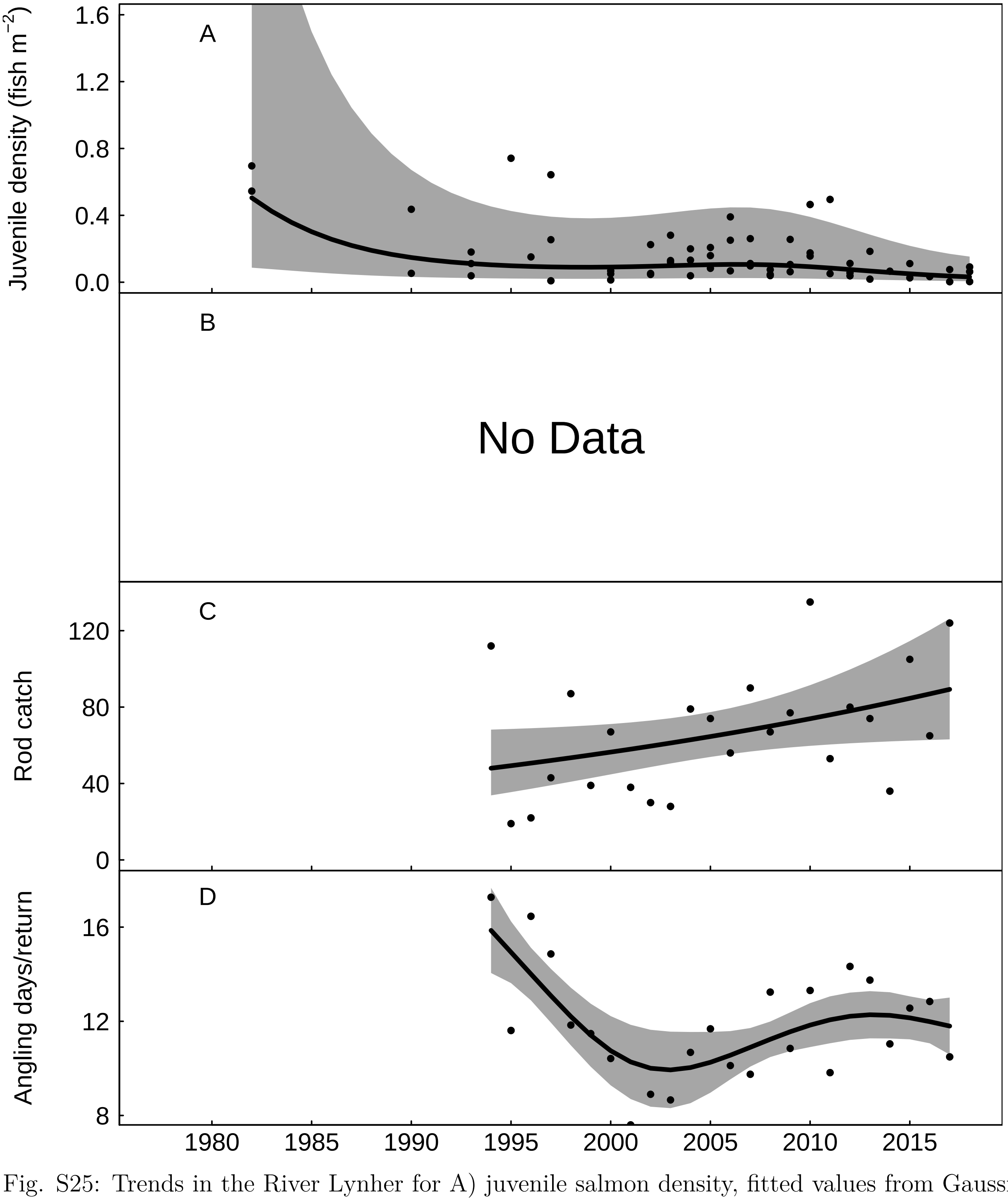


Fig. S25: Trends in the River Lynher for A) juvenile salmon density, fitted values from Gaussian additive mixed model, grey areas represent 95% point-wise confidence bands for the smoother, B) No Data, C) rod catch, fitted values from negative binomial generalized additive model, grey areas represent 95% point-wise confidence bands for the smoother, and D) angling effort measured as number of days fished per licence return, fitted values from Gaussian additive model, grey areas represent 95% point-wise confidence bands for the smoother.

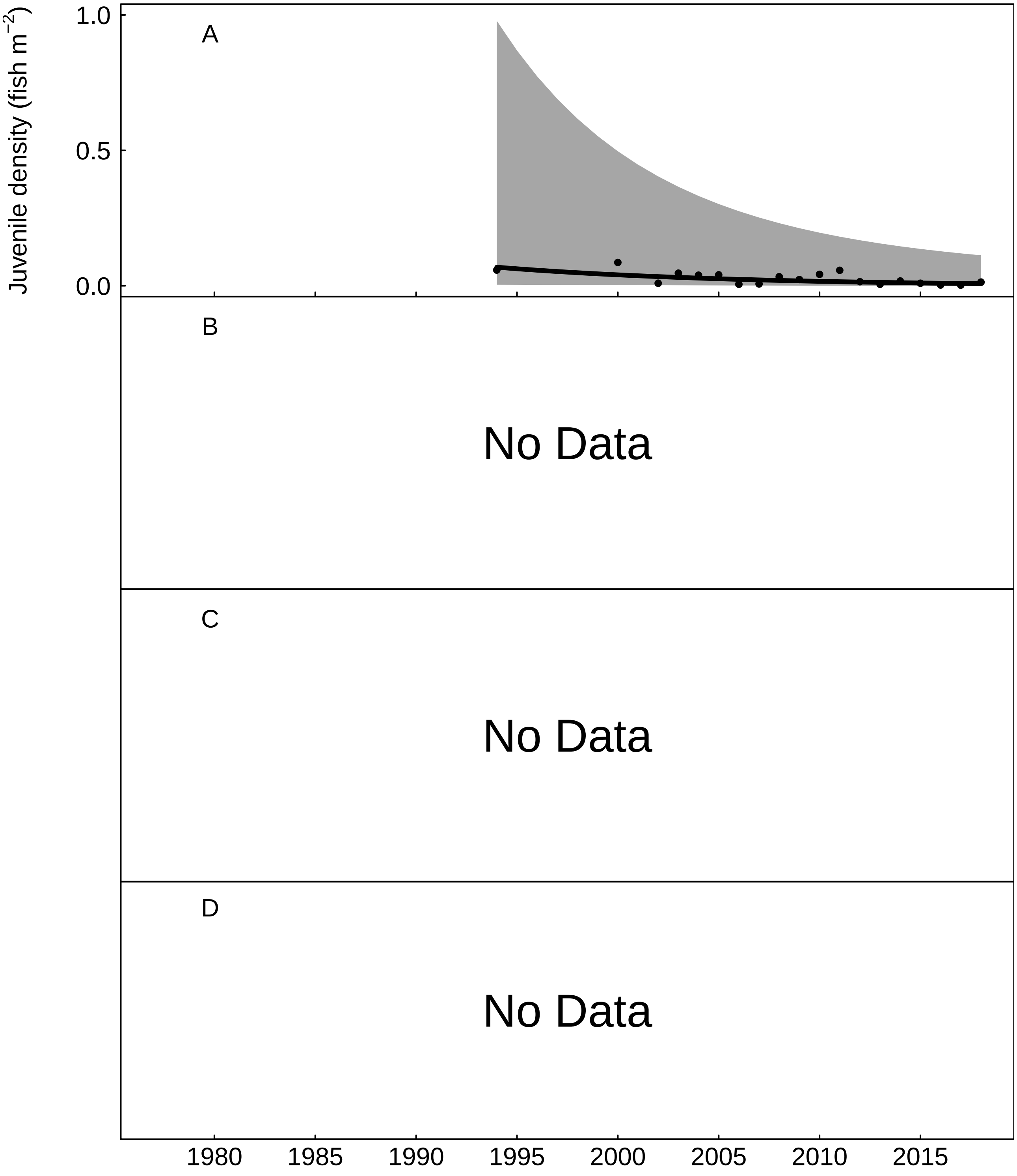


Fig. S26: Trends in the West Looe for A) juvenile salmon density, fitted values from Gaussian additive mixed model, grey areas represent 95% point-wise confidence bands for the smoother, B) No Data, C) No Data, and D) No Data.

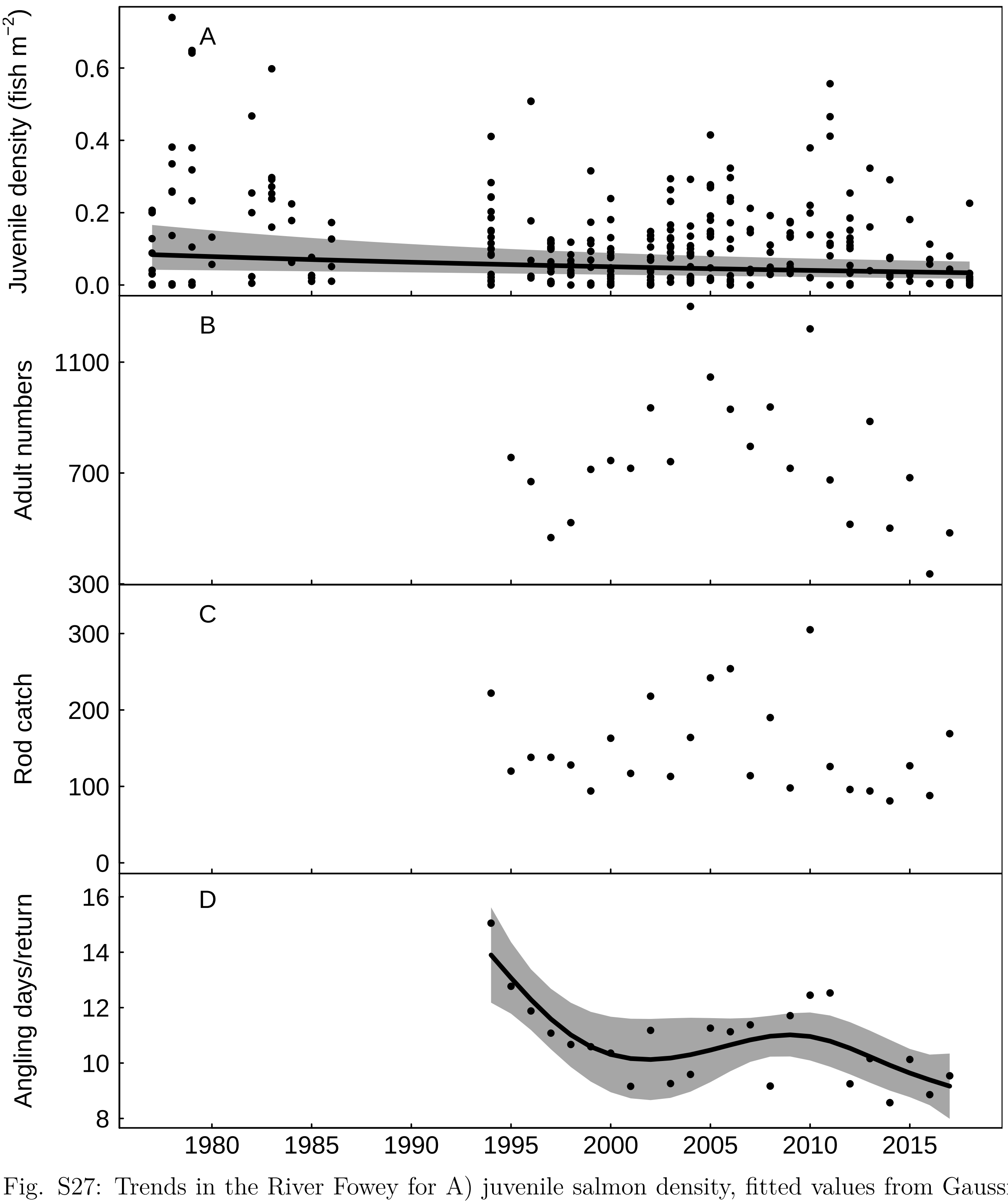


Fig. S27: Trends in the River Fowey for A) juvenile salmon density, fitted values from Gaussian additive mixed model, grey areas represent 95% point-wise confidence bands for the smoother, B) returning adult numbers, C) rod catch, and D) angling effort measured as number of days fished per licence return, fitted values from Gaussian additive model, grey areas represent 95% point-wise confidence bands for the smoother.

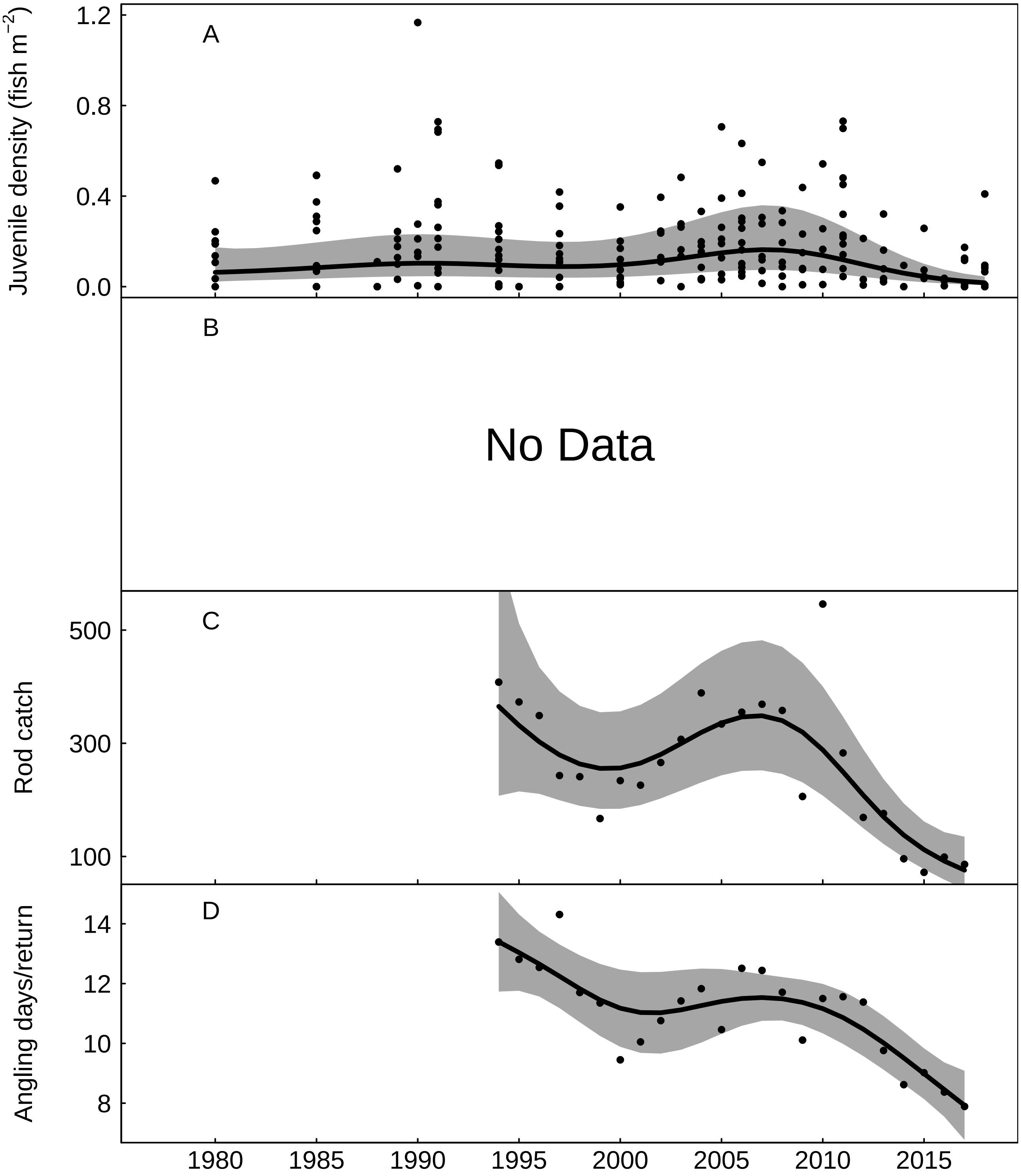


Fig. S28: Trends in the River Camel for A) juvenile salmon density, fitted values from Gaussian additive mixed model, grey areas represent 95% point-wise confidence bands for the smoother, B) No Data, C) rod catch, fitted values from negative binomial generalized additive model, grey areas represent 95% point-wise confidence bands for the smoother, and D) angling effort measured as number of days fished per licence return, fitted values from Gaussian additive model, grey areas represent 95% point-wise confidence bands for the smoother.

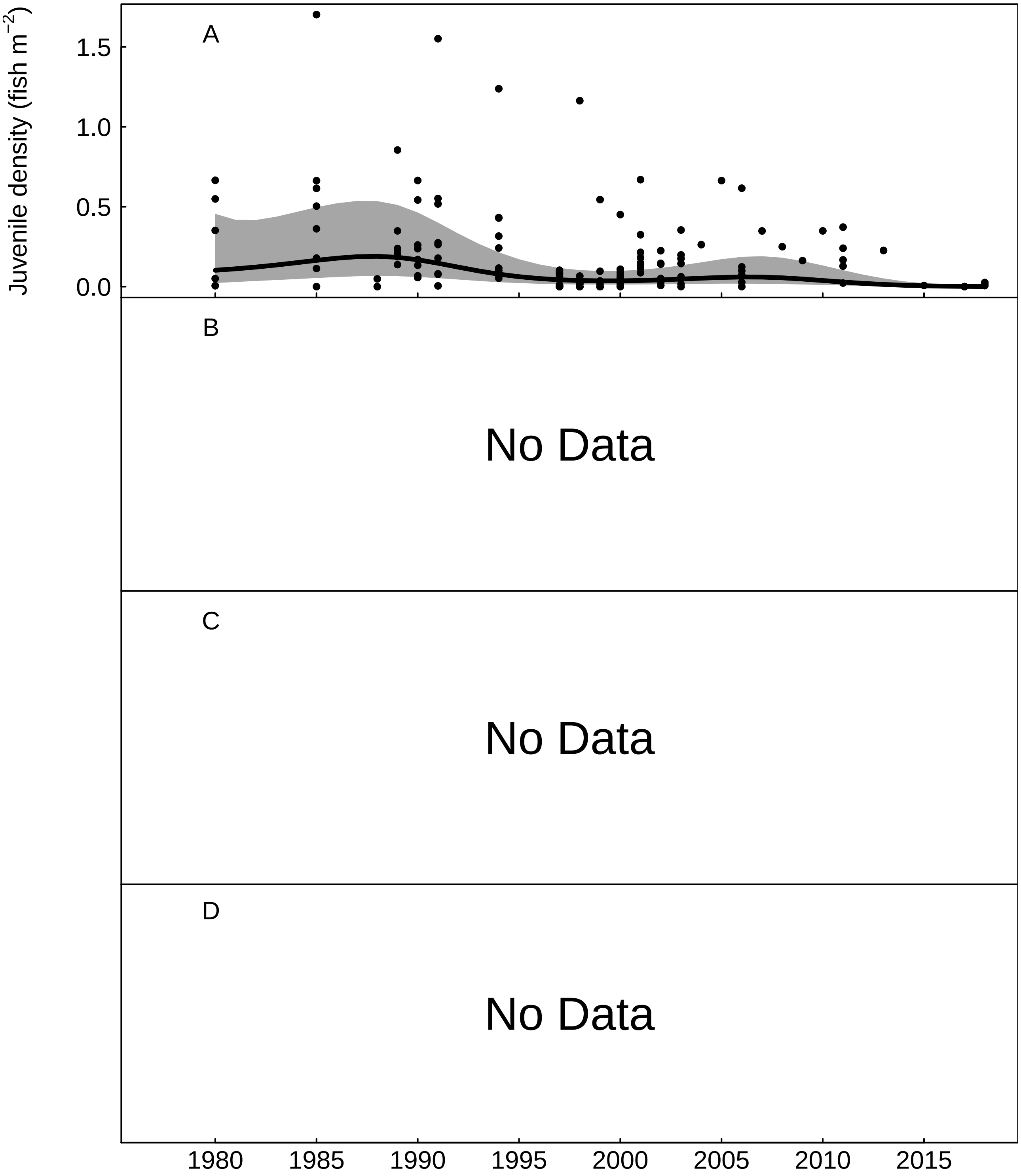


Fig. S29: Trends in the River Allen for A) juvenile salmon density, fitted values from Gaussian additive mixed model, grey areas represent 95% point-wise confidence bands for the smoother, B) No Data, C) No Data, and D) No Data.

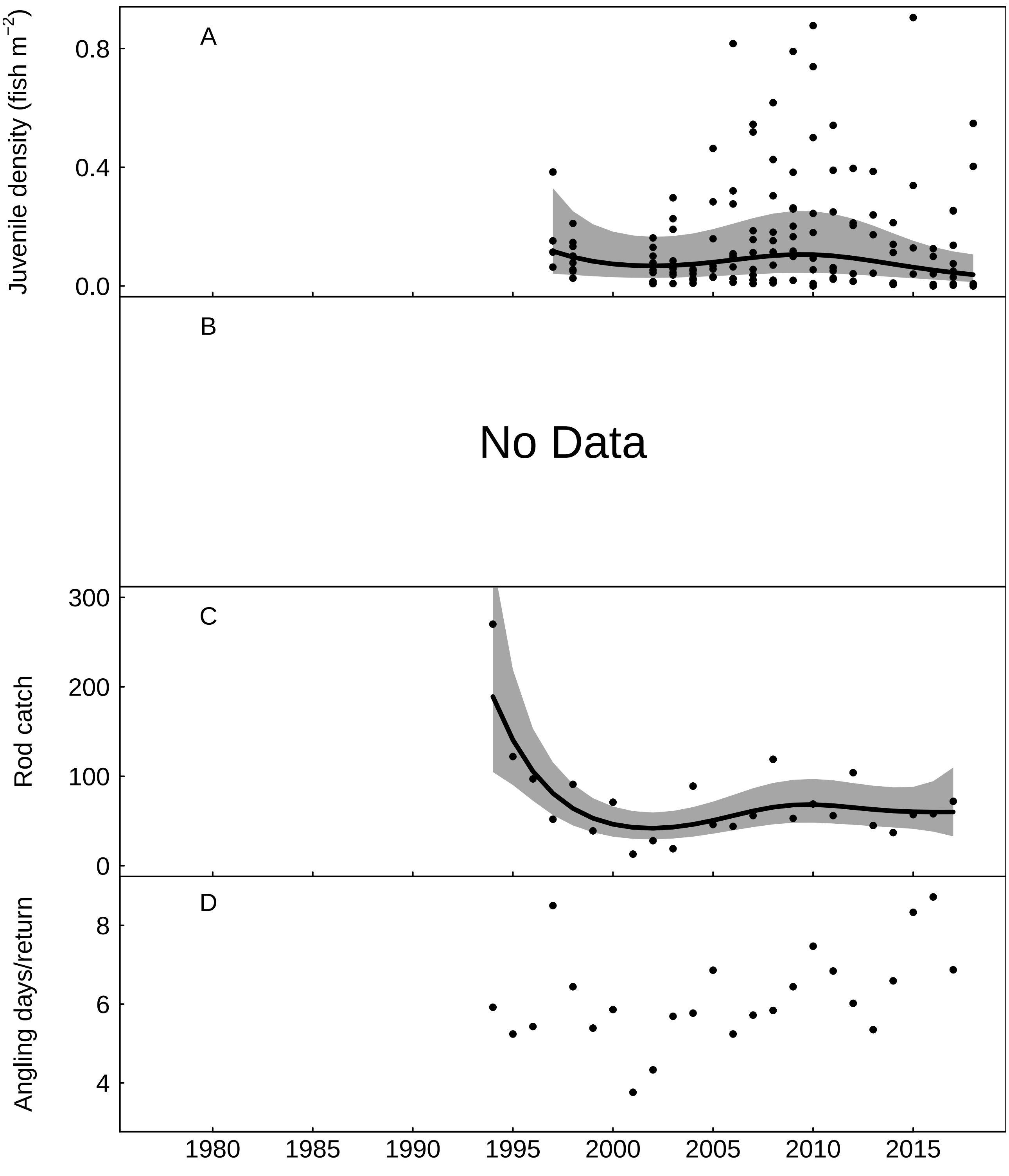


Fig. S30: Trends in the River Torridge for A) juvenile salmon density, fitted values from Gaussian additive mixed model, grey areas represent 95% point-wise confidence bands for the smoother, B) No Data, C) rod catch, fitted values from negative binomial generalized additive model, grey areas represent 95% point-wise confidence bands for the smoother, and D) angling effort measured as number of days fished per licence return.

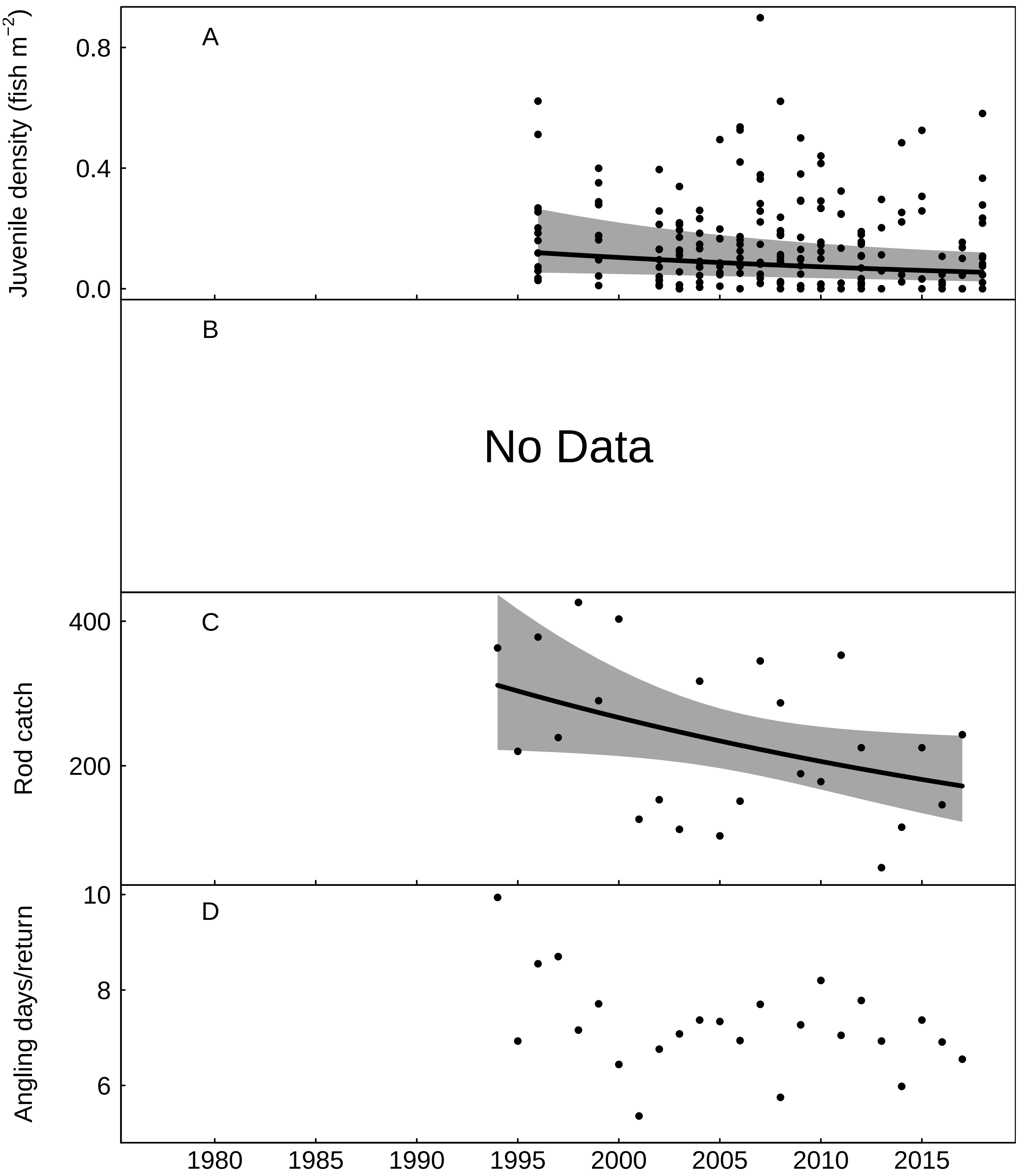


Fig. S31: Trends in the River Taw for A) juvenile salmon density, fitted values from Gaussian additive mixed model, grey areas represent 95% point-wise confidence bands for the smoother, B) No Data, C) rod catch, fitted values from negative binomial generalized additive model, grey areas represent 95% point-wise confidence bands for the smoother, and D) angling effort measured as number of days fished per licence return.

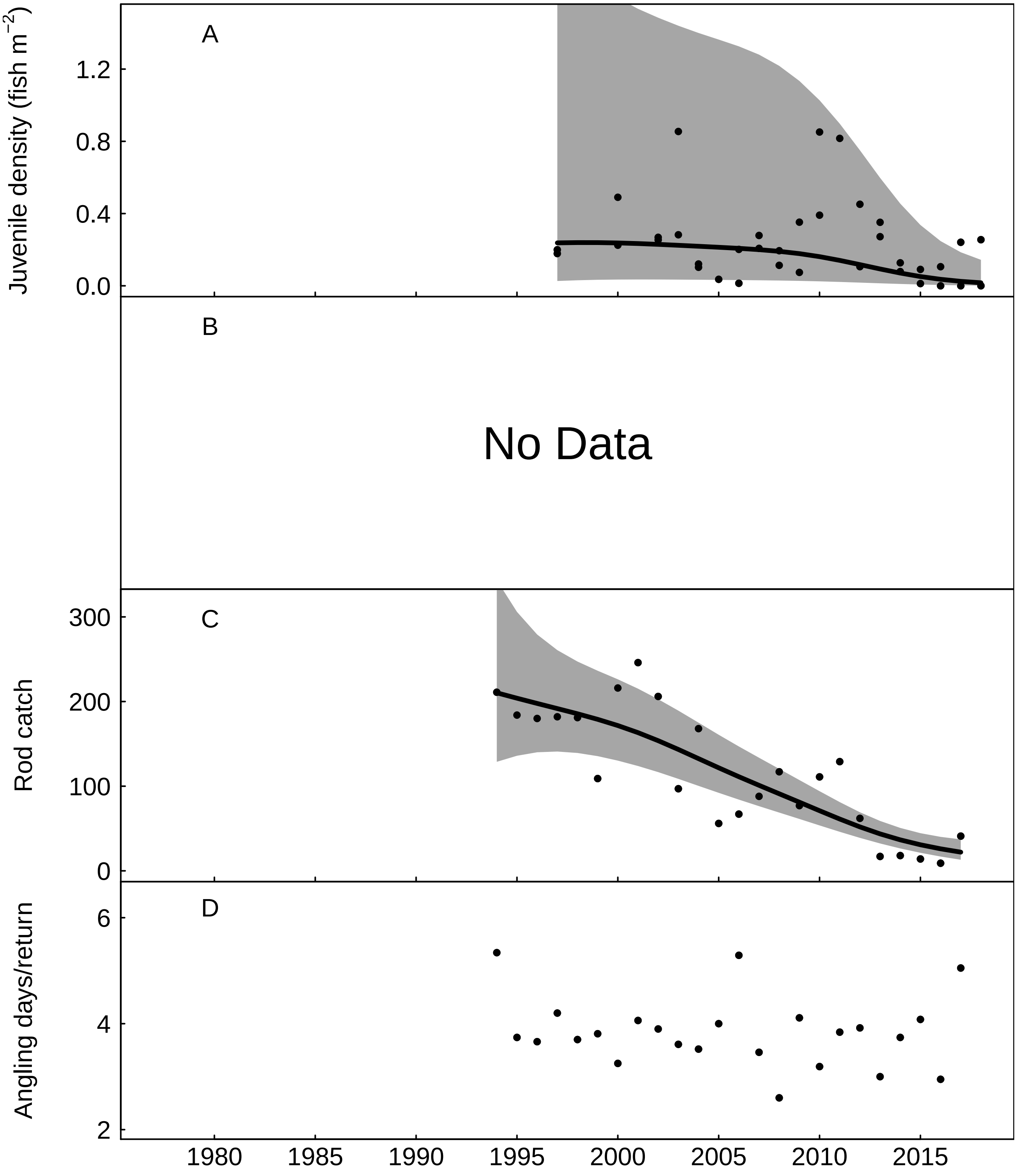


Fig. S32: Trends in the River Lyn for A) juvenile salmon density, fitted values from Gaussian additive mixed model, grey areas represent 95% point-wise confidence bands for the smoother, B) No Data, C) rod catch, fitted values from negative binomial generalized additive model, grey areas represent 95% point-wise confidence bands for the smoother, and D) angling effort measured as number of days fished per licence return.

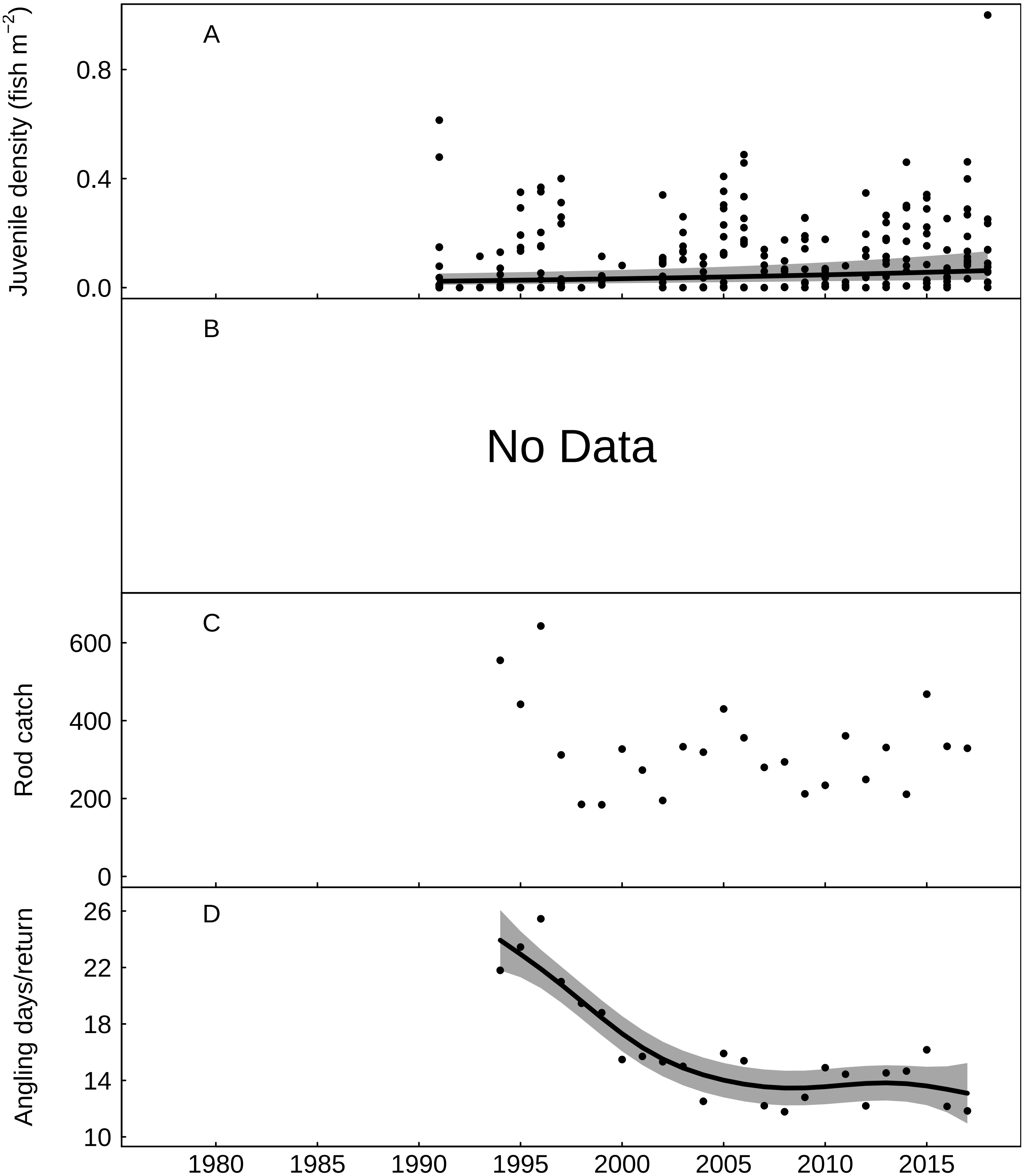


Fig. S33: Trends in the River Severn for A) juvenile salmon density, fitted values from Gaussian additive mixed model, grey areas represent 95% point-wise confidence bands for the smoother, B) No Data, C) rod catch, and D) angling effort measured as number of days fished per licence return, fitted values from Gaussian additive model, grey areas represent 95% point-wise confidence bands for the smoother.

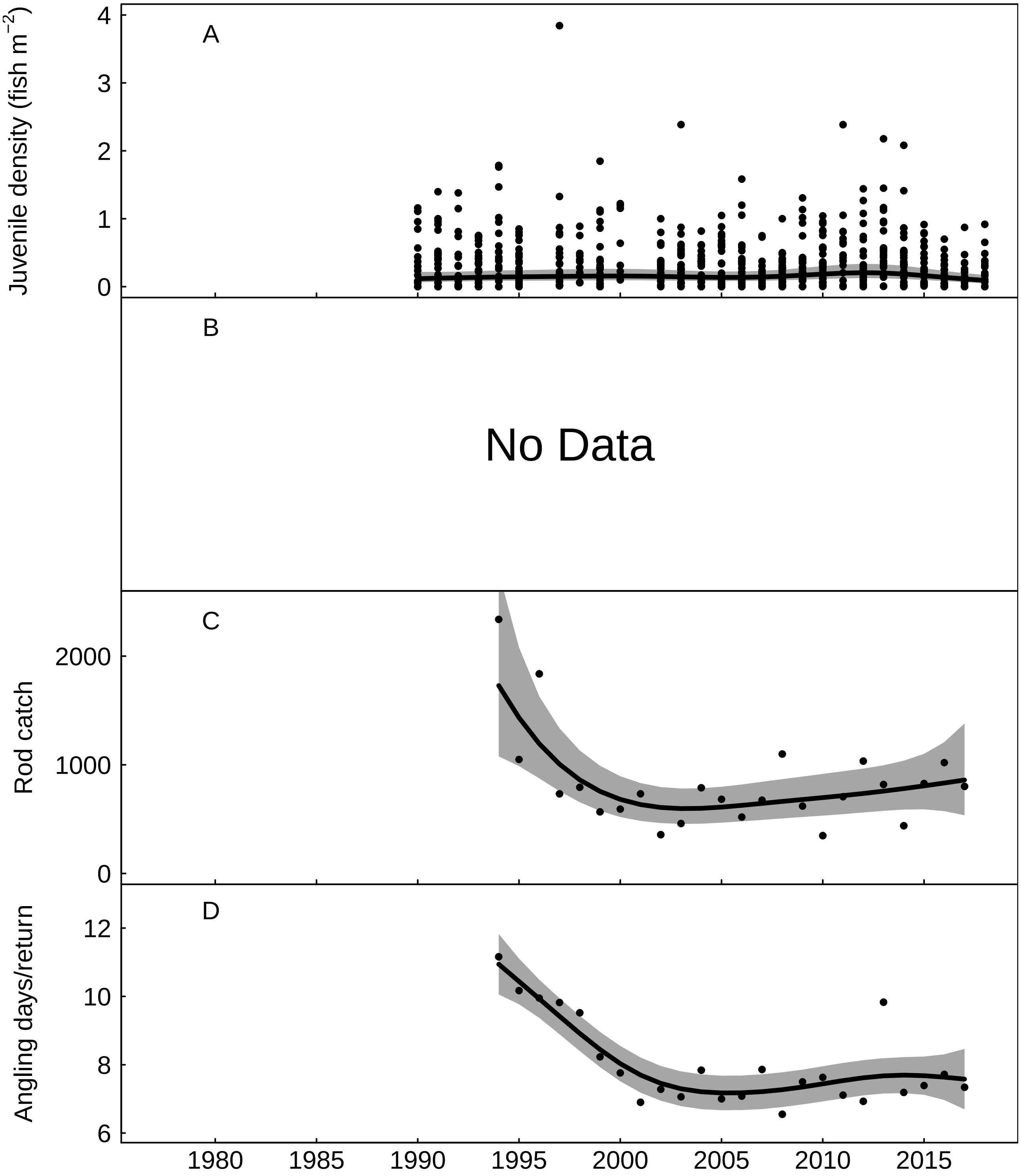


Fig. S34: Trends in the River Wye for A) juvenile salmon density, fitted values from Gaussian additive mixed model, grey areas represent 95% point-wise confidence bands for the smoother, B) No Data, C) rod catch, fitted values from negative binomial generalized additive model, grey areas represent 95% point-wise confidence bands for the smoother, and D) angling effort measured as number of days fished per licence return, fitted values from Gaussian additive model, grey areas represent 95% point-wise confidence bands for the smoother.

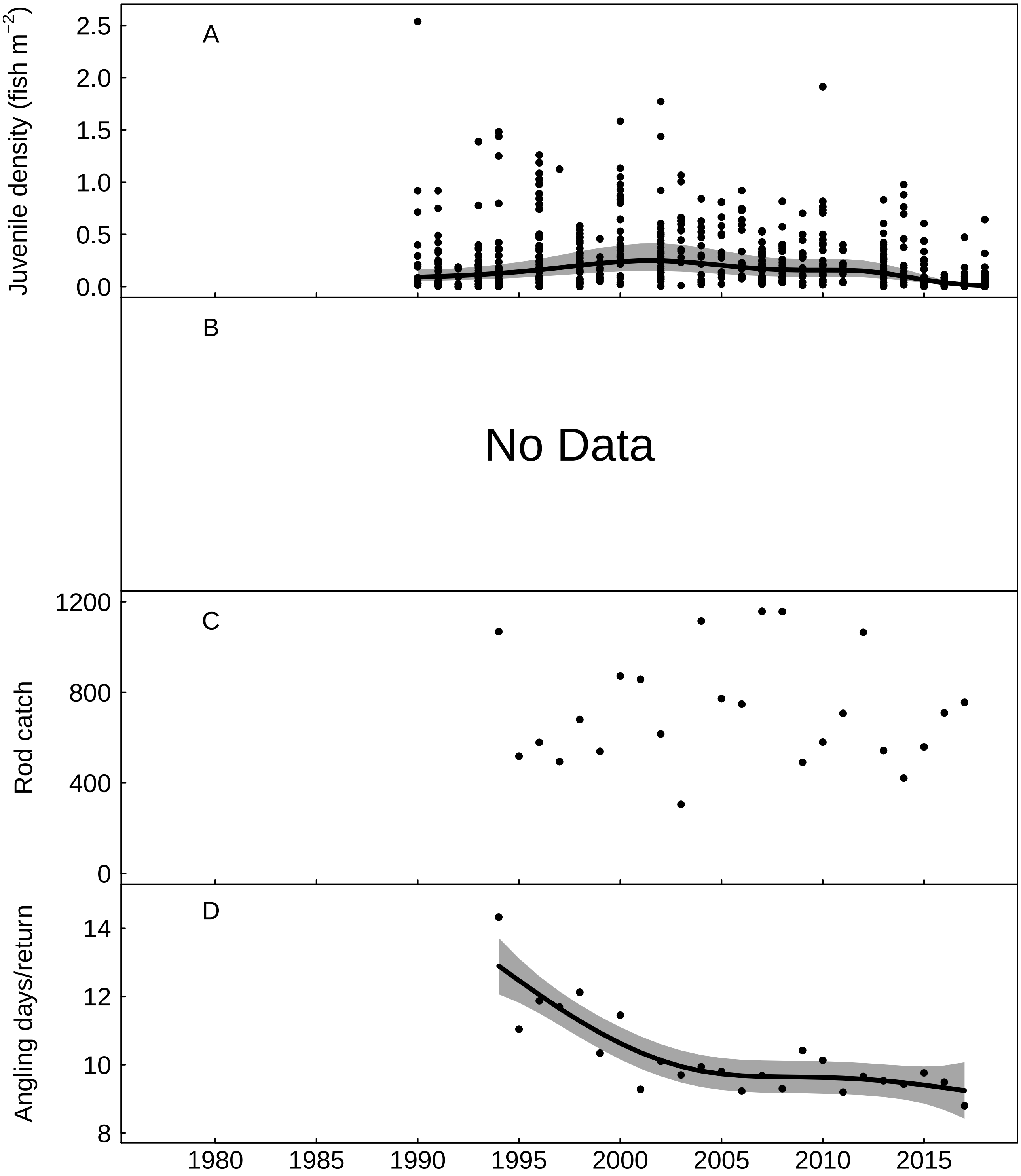


Fig. S35: Trends in the River Usk for A) juvenile salmon density, fitted values from Gaussian additive mixed model, grey areas represent 95% point-wise confidence bands for the smoother, B) No Data, C) rod catch, and D) angling effort measured as number of days fished per licence return, fitted values from Gaussian additive model, grey areas represent 95% point-wise confidence bands for the smoother.

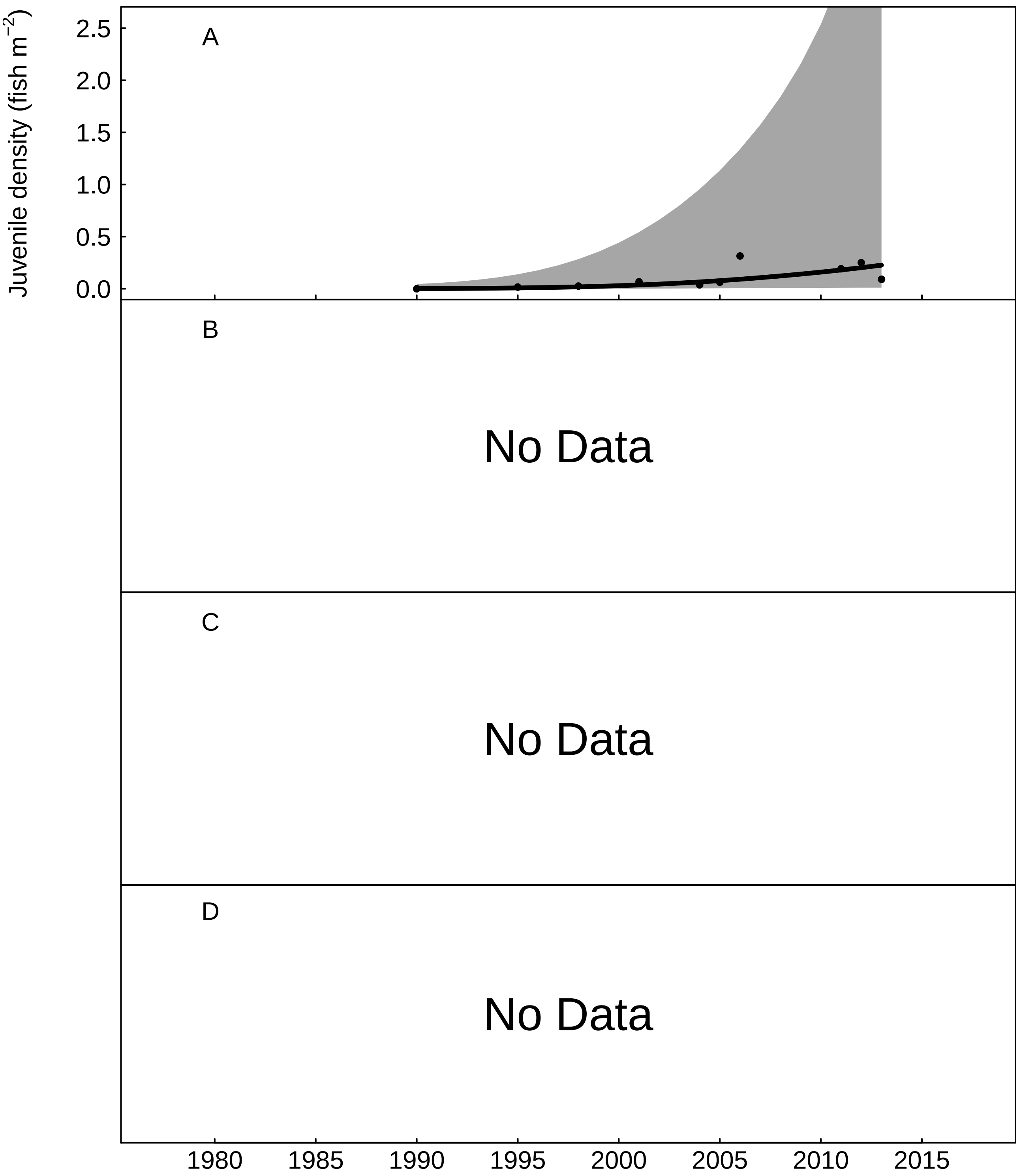


Fig. S36: Trends in the River Rhymney for A) juvenile salmon density, fitted values from Gaussian additive mixed model, grey areas represent 95% point-wise confidence bands for the smoother, B) No Data, C) No Data, and D) No Data.

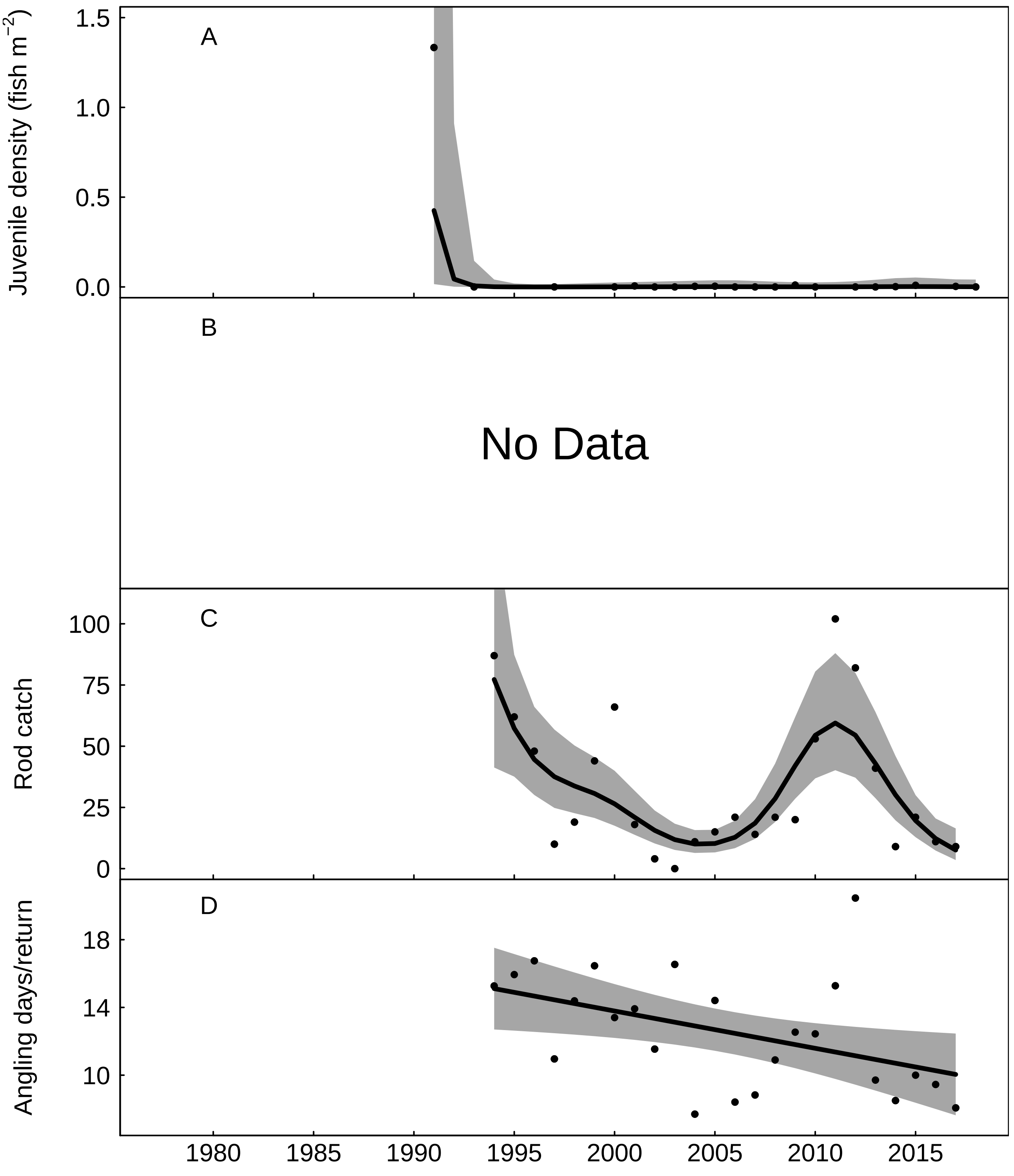


Fig. S37: Trends in the River Taff for A) juvenile salmon density, fitted values from Gaussian additive mixed model, grey areas represent 95% point-wise confidence bands for the smoother, B) No Data, C) rod catch, fitted values from negative binomial generalized additive model, grey areas represent 95% point-wise confidence bands for the smoother, and D) angling effort measured as number of days fished per licence return, fitted values from Gaussian additive model, grey areas represent 95% point-wise confidence bands for the smoother.

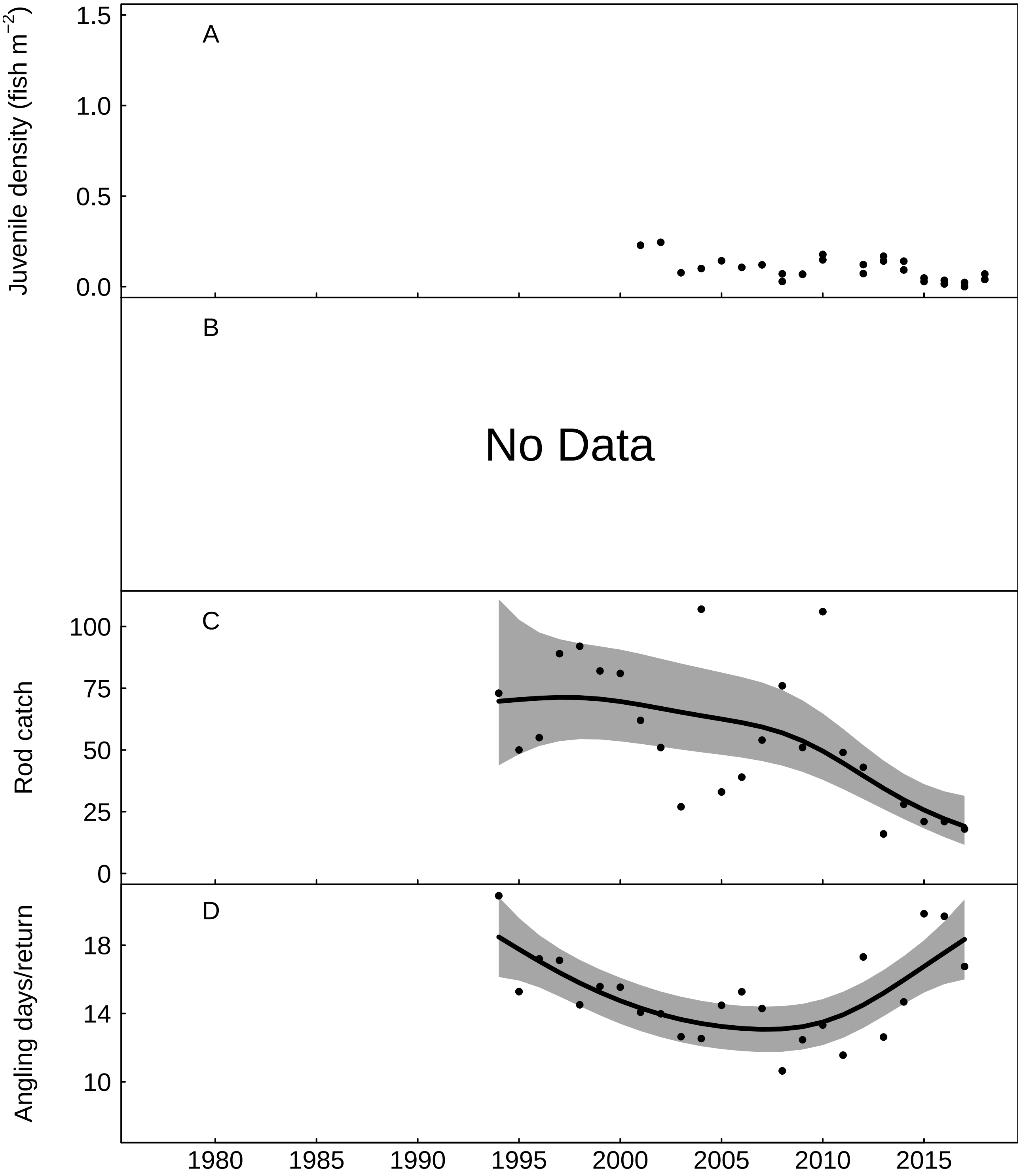


Fig. S38: Trends in the River Ogmore for A) juvenile salmon density, B) No Data, C) rod catch, fitted values from negative binomial generalized additive model, grey areas represent 95% point-wise confidence bands for the smoother, and D) angling effort measured as number of days fished per licence return, fitted values from Gaussian additive model, grey areas represent 95% point-wise confidence bands for the smoother.

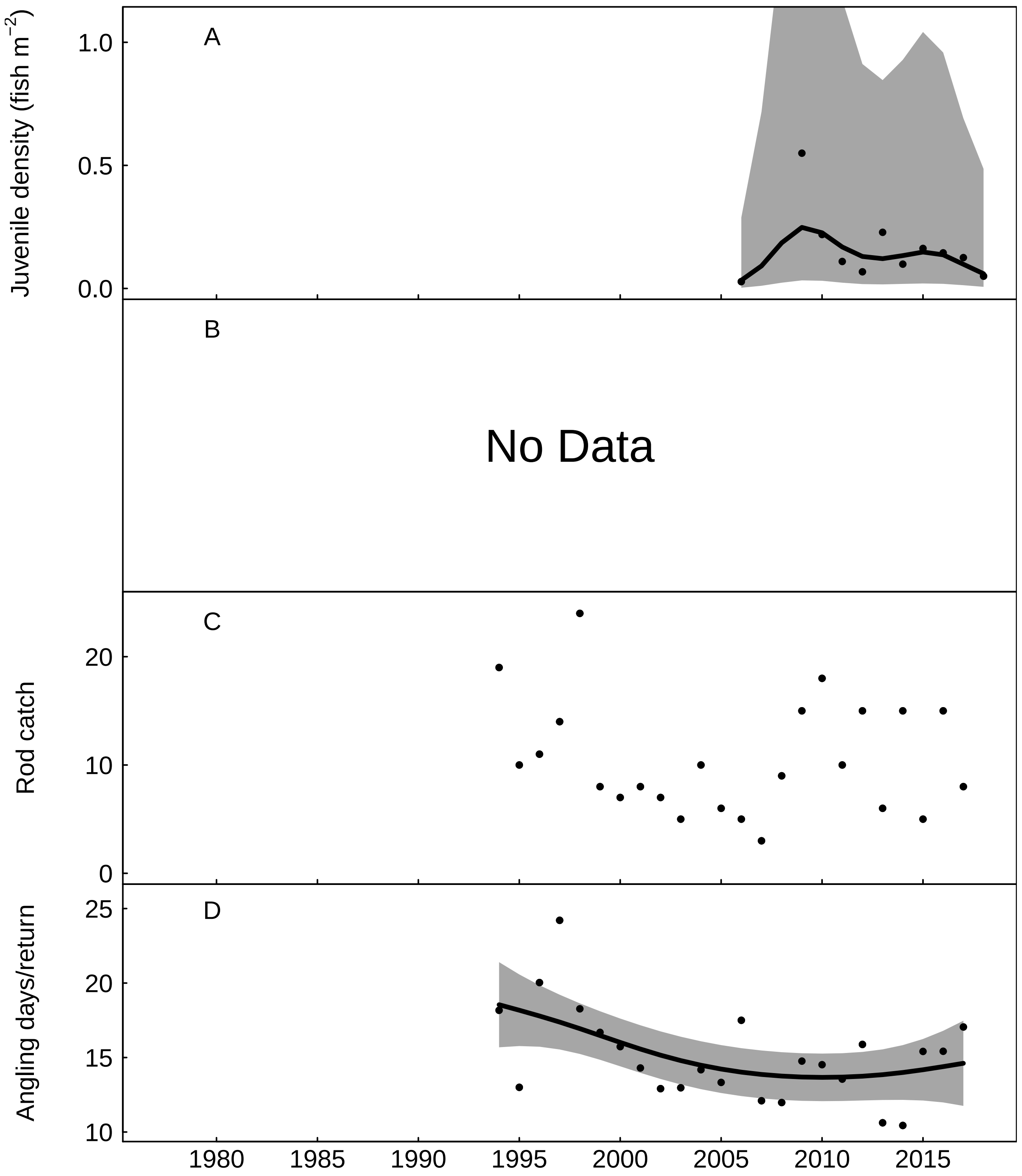


Fig. S39: Trends in the Afan for A) juvenile salmon density, fitted values from Gaussian additive mixed model, grey areas represent 95% point-wise confidence bands for the smoother, B) No Data, C) rod catch, and D) angling effort measured as number of days fished per licence return, fitted values from Gaussian additive model, grey areas represent 95% point-wise confidence bands for the smoother.

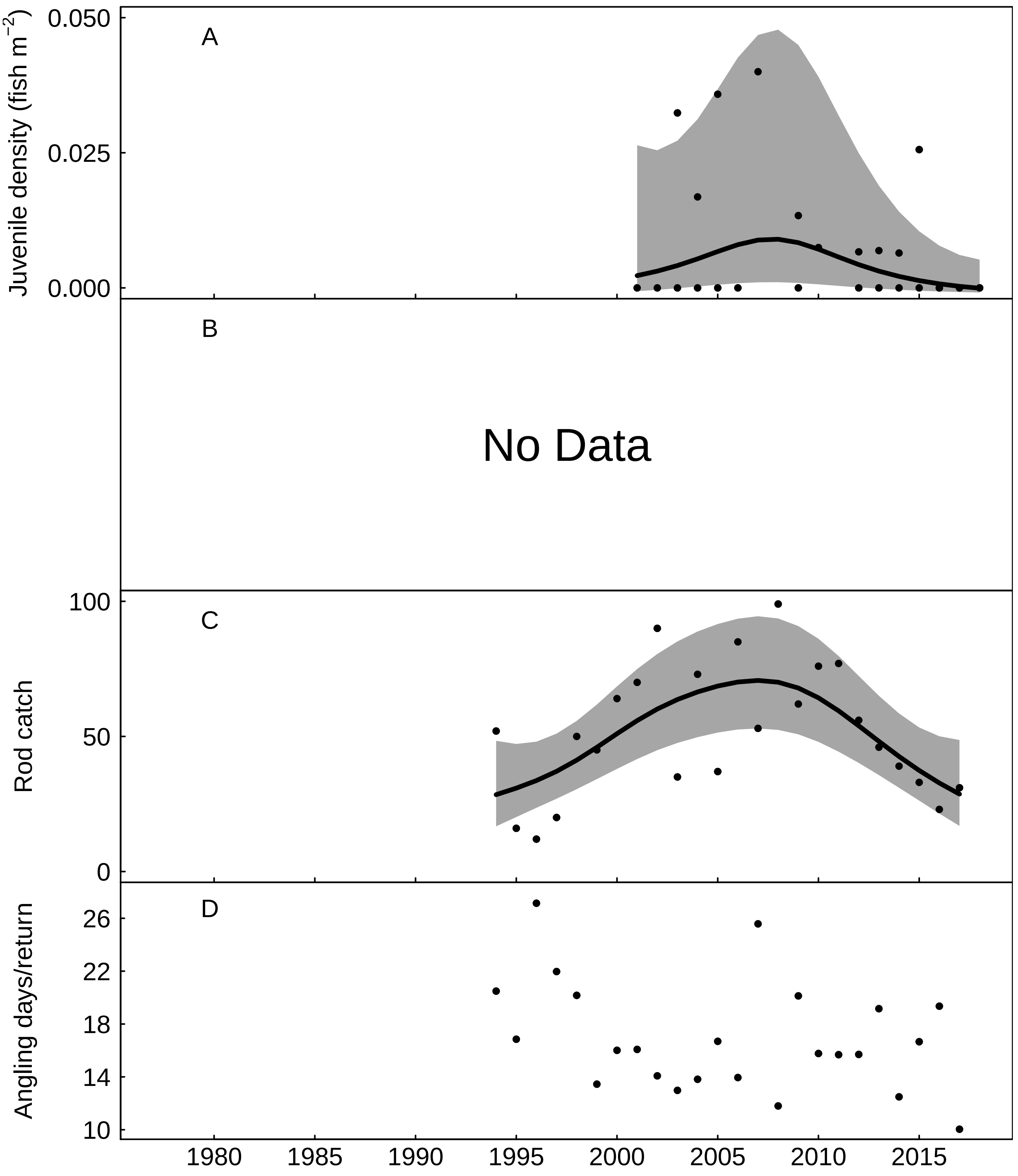


Fig. S40: Trends in the River Neath for A) juvenile salmon density, fitted values from Gaussian additive mixed model, grey areas represent 95% point-wise confidence bands for the smoother, B) No Data, C) rod catch, fitted values from negative binomial generalized additive model, grey areas represent 95% point-wise confidence bands for the smoother, and D) angling effort measured as number of days fished per licence return.

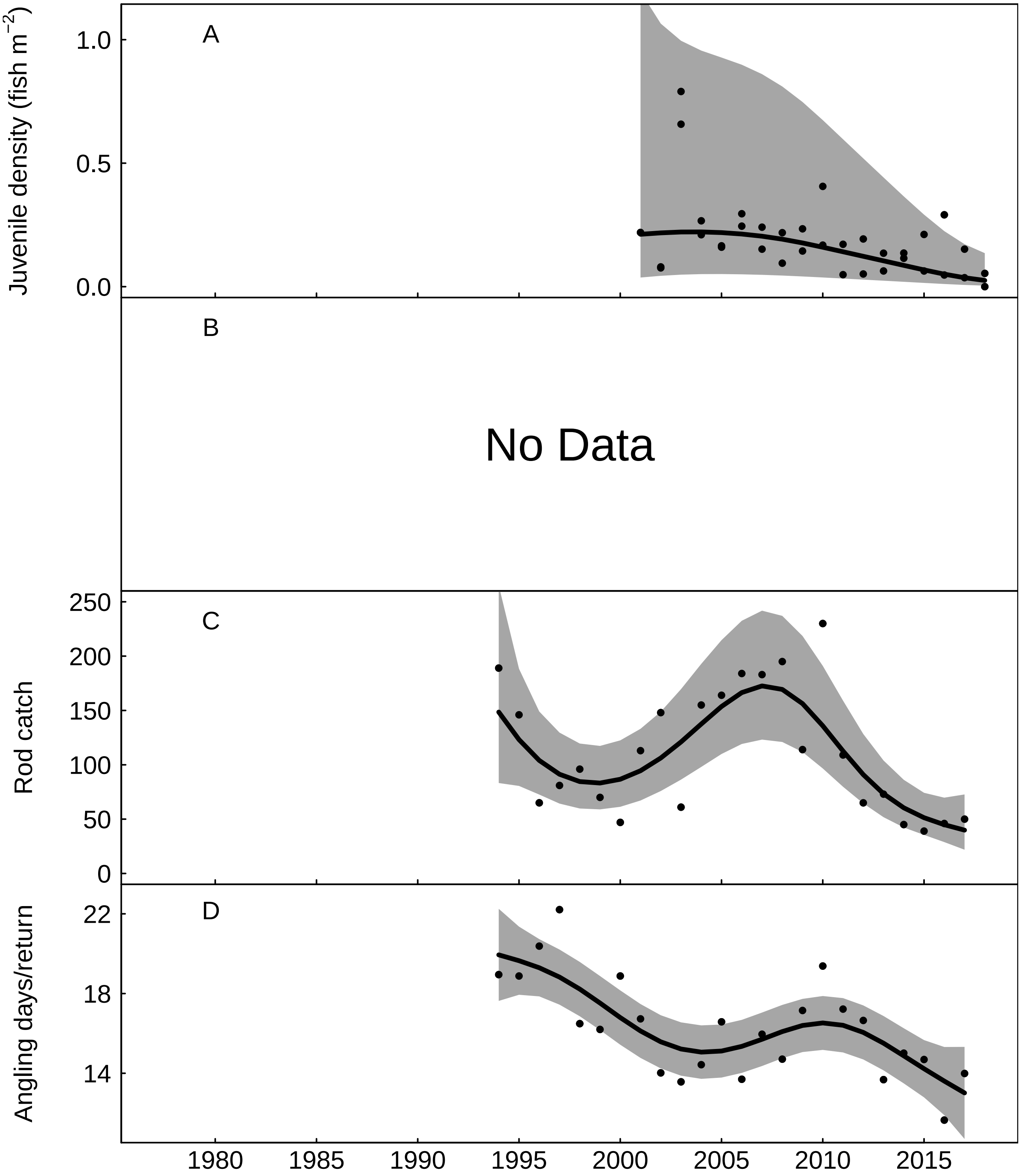


Fig. S41: Trends in the River Tawe for A) juvenile salmon density, fitted values from Gaussian additive mixed model, grey areas represent 95% point-wise confidence bands for the smoother, B) No Data, C) rod catch, fitted values from negative binomial generalized additive model, grey areas represent 95% point-wise confidence bands for the smoother, and D) angling effort measured as number of days fished per licence return, fitted values from Gaussian additive model, grey areas represent 95% point-wise confidence bands for the smoother.

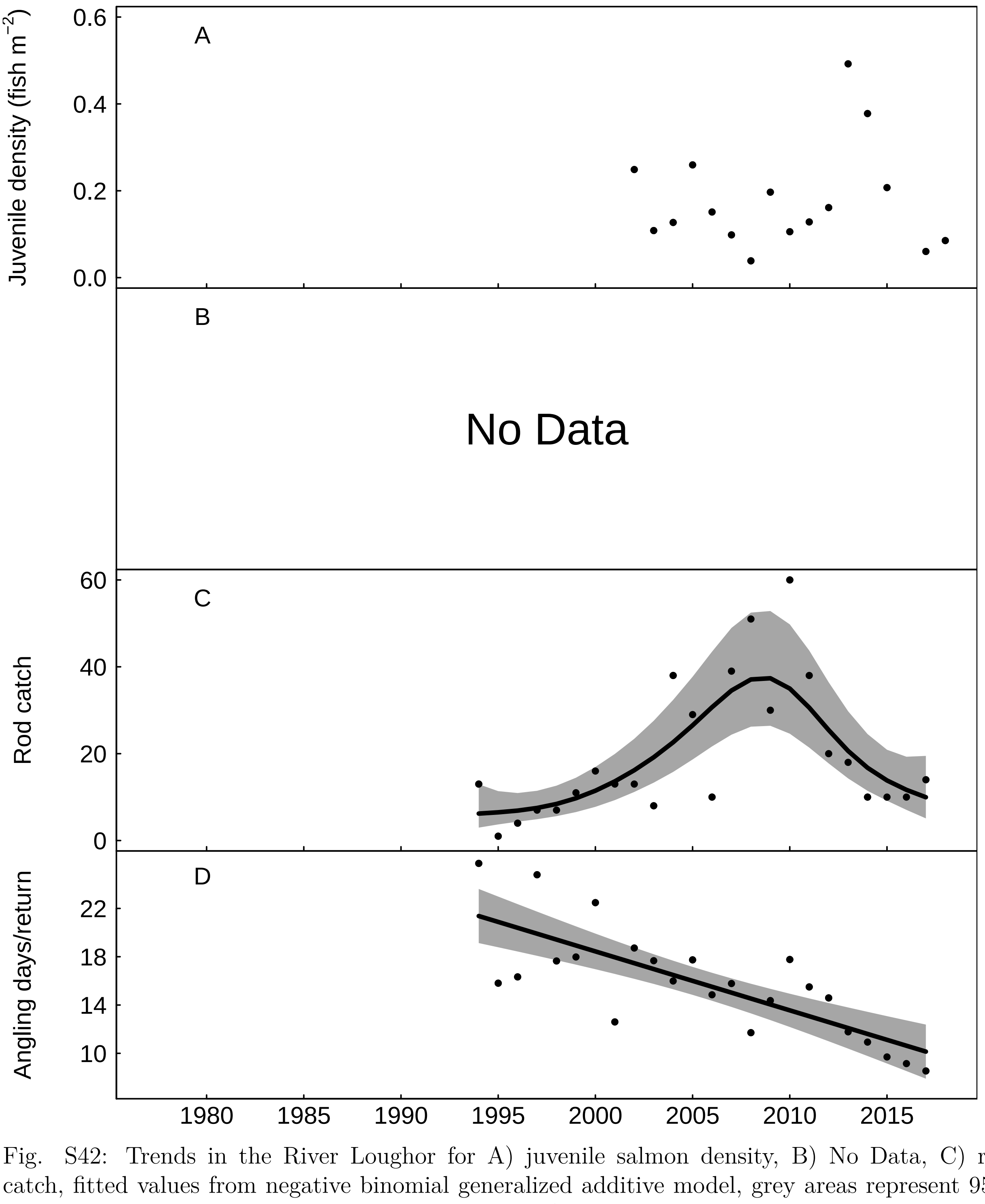


Fig. S42: Trends in the River Loughor for A) juvenile salmon density, B) No Data, C) rod catch, fitted values from negative binomial generalized additive model, grey areas represent 95% point-wise confidence bands for the smoother, and D) angling effort measured as number of days fished per licence return, fitted values from Gaussian additive model, grey areas represent 95% point-wise confidence bands for the smoother.

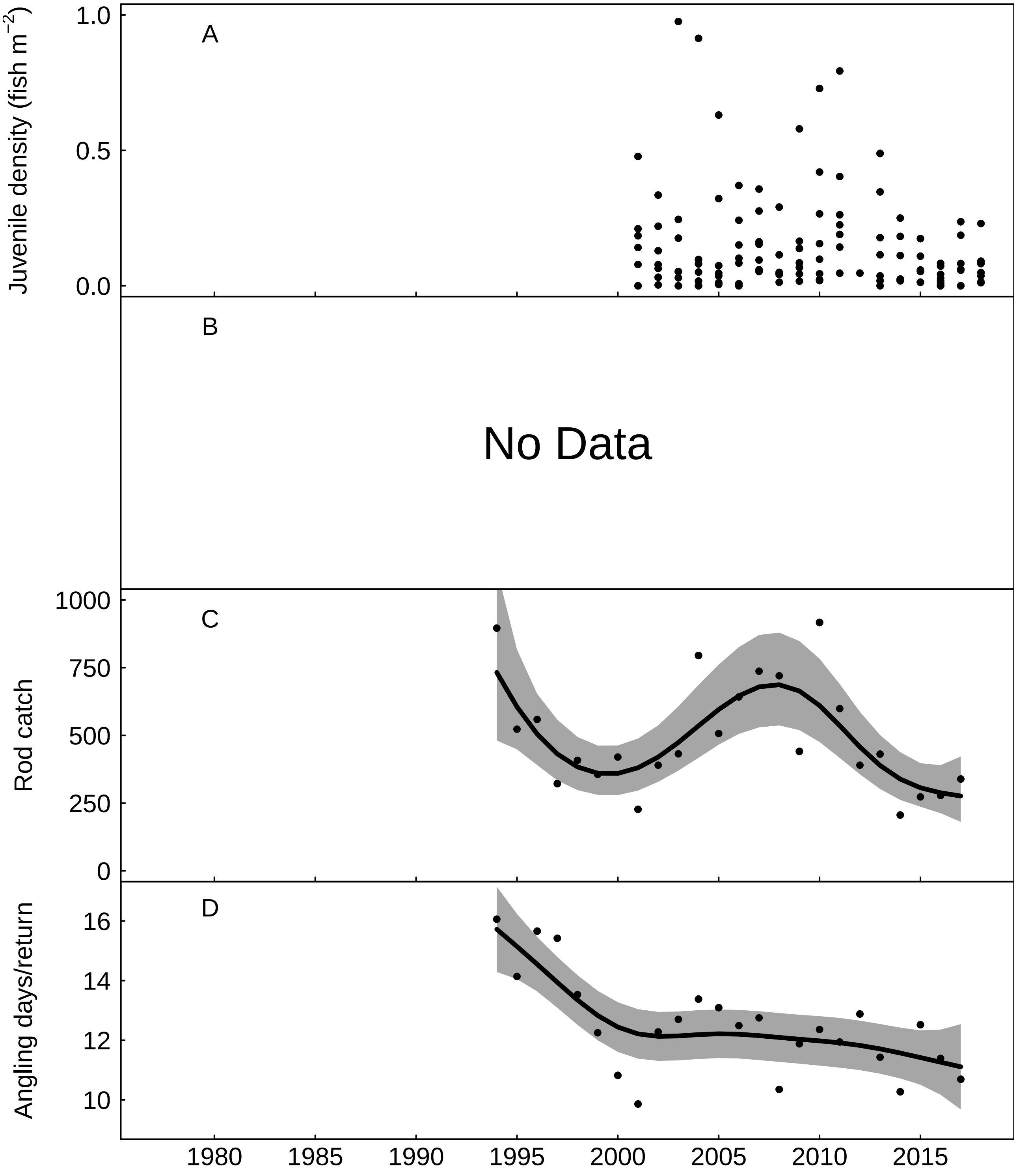


Fig. S43: Trends in the Tywi for A) juvenile salmon density, B) No Data, C) rod catch, fitted values from negative binomial generalized additive model, grey areas represent 95% point-wise confidence bands for the smoother, and D) angling effort measured as number of days fished per licence return, fitted values from Gaussian additive model, grey areas represent 95% point-wise confidence bands for the smoother.

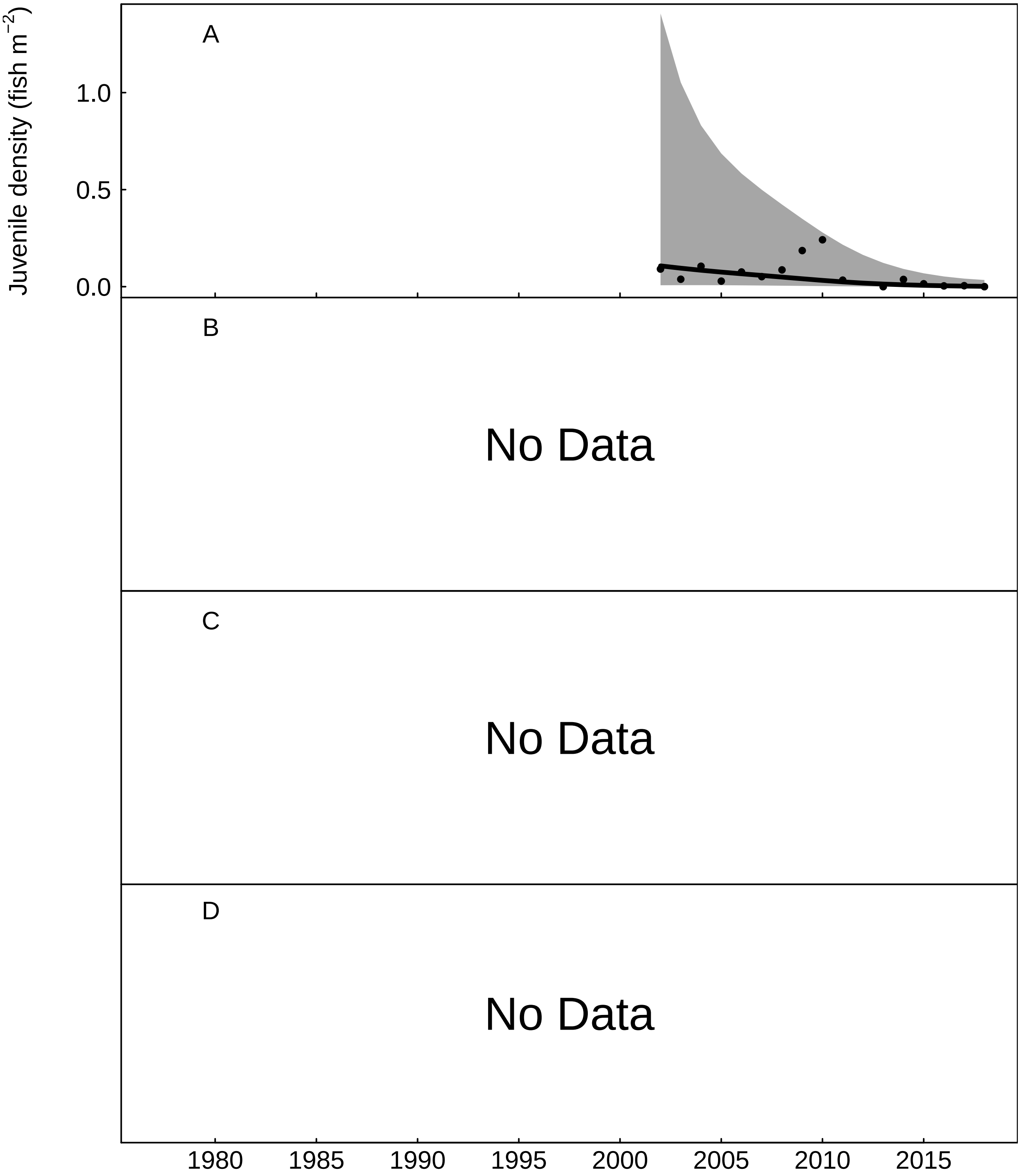


Fig. S44: Trends in the Gwili for A) juvenile salmon density, fitted values from Gaussian additive mixed model, grey areas represent 95% point-wise confidence bands for the smoother, B) No Data, C) No Data, and D) No Data.

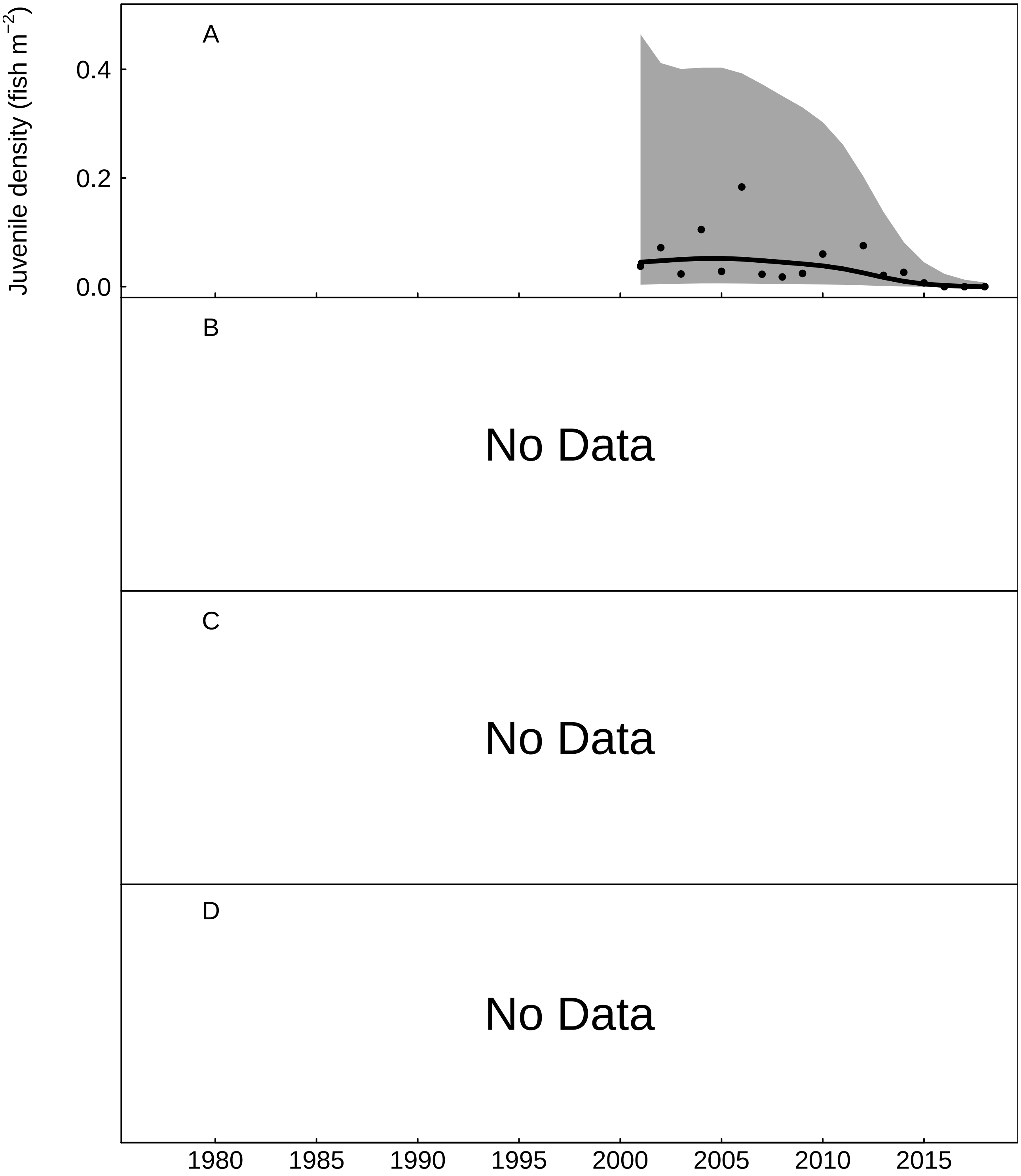


Fig. S45: Trends in the Cynin for A) juvenile salmon density, fitted values from Gaussian additive mixed model, grey areas represent 95% point-wise confidence bands for the smoother, B) No Data, C) No Data, and D) No Data.

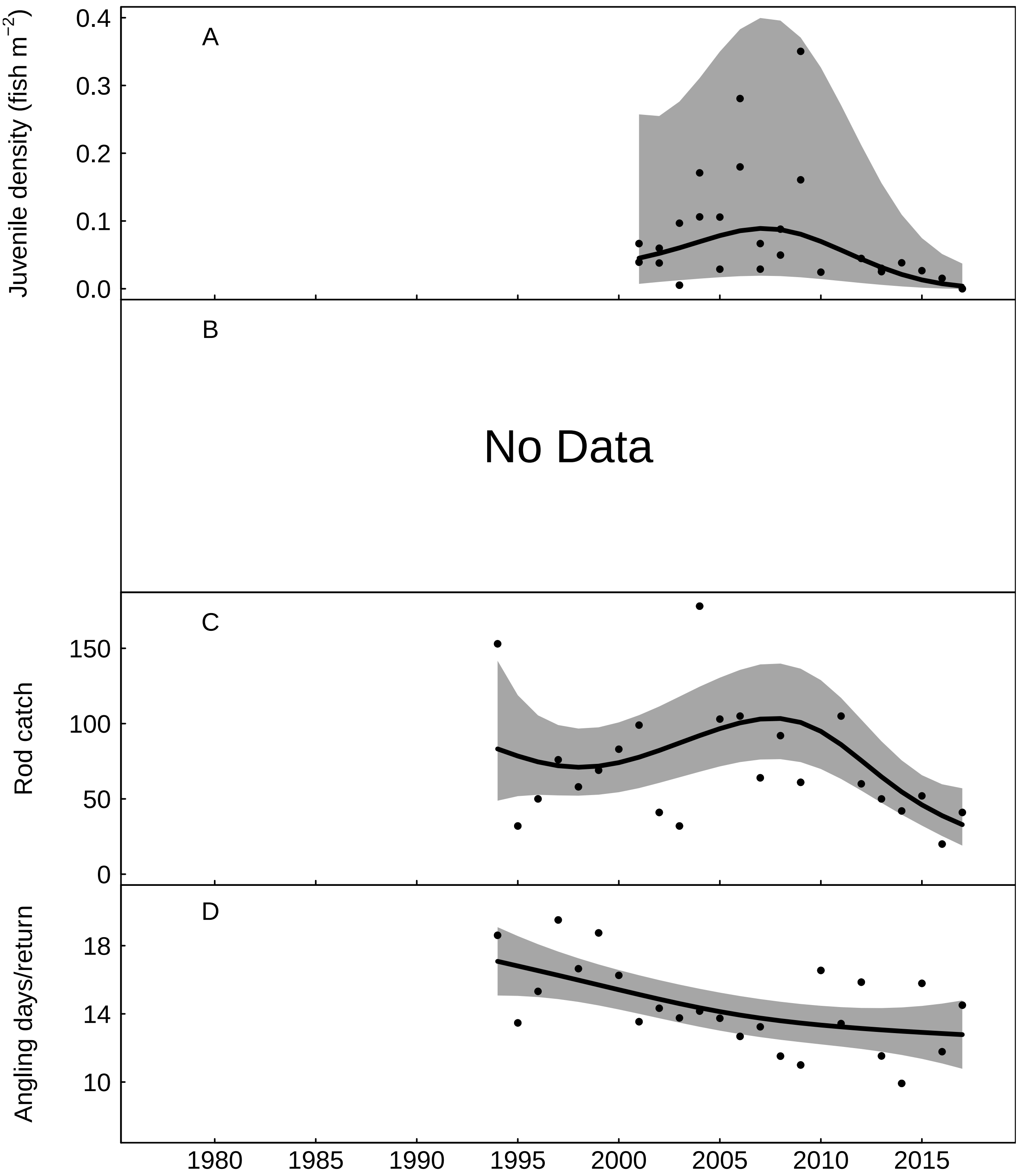


Fig. S46: Trends in the Taf for A) juvenile salmon density, fitted values from Gaussian additive mixed model, grey areas represent 95% point-wise confidence bands for the smoother, B) No Data, C) rod catch, fitted values from negative binomial generalized additive model, grey areas represent 95% point-wise confidence bands for the smoother, and D) angling effort measured as number of days fished per licence return, fitted values from Gaussian additive model, grey areas represent 95% point-wise confidence bands for the smoother.

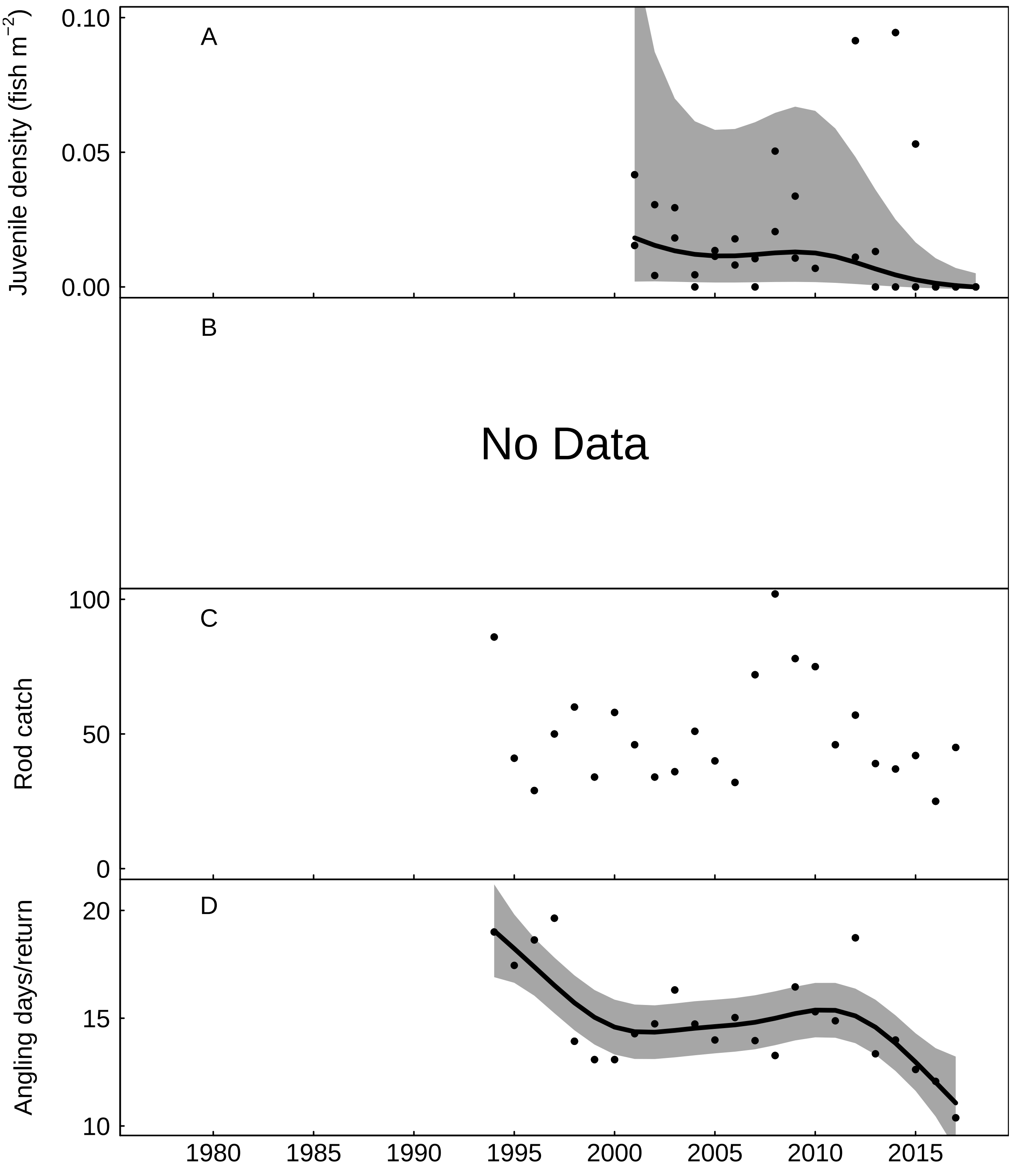


Fig. S47: Trends in the Eastern Cleddau for A) juvenile salmon density, fitted values from Gaussian additive mixed model, grey areas represent 95% point-wise confidence bands for the smoother, B) No Data, C) rod catch, and D) angling effort measured as number of days fished per licence return, fitted values from Gaussian additive model, grey areas represent 95% point-wise confidence bands for the smoother.

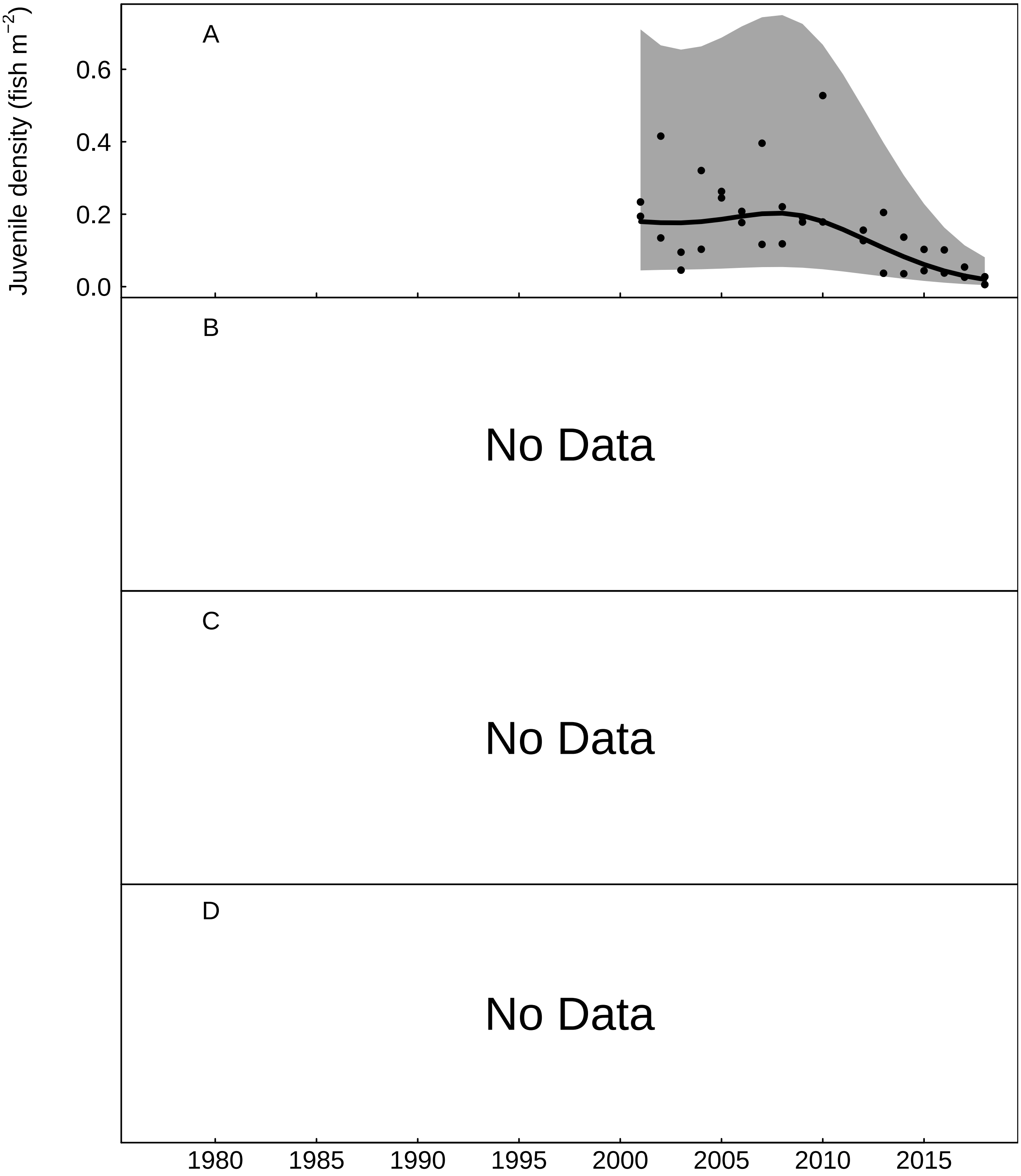


Fig. S48: Trends in the Western Cleddau for A) juvenile salmon density, fitted values from Gaussian additive mixed model, grey areas represent 95% point-wise confidence bands for the smoother, B) No Data, C) No Data, and D) No Data.

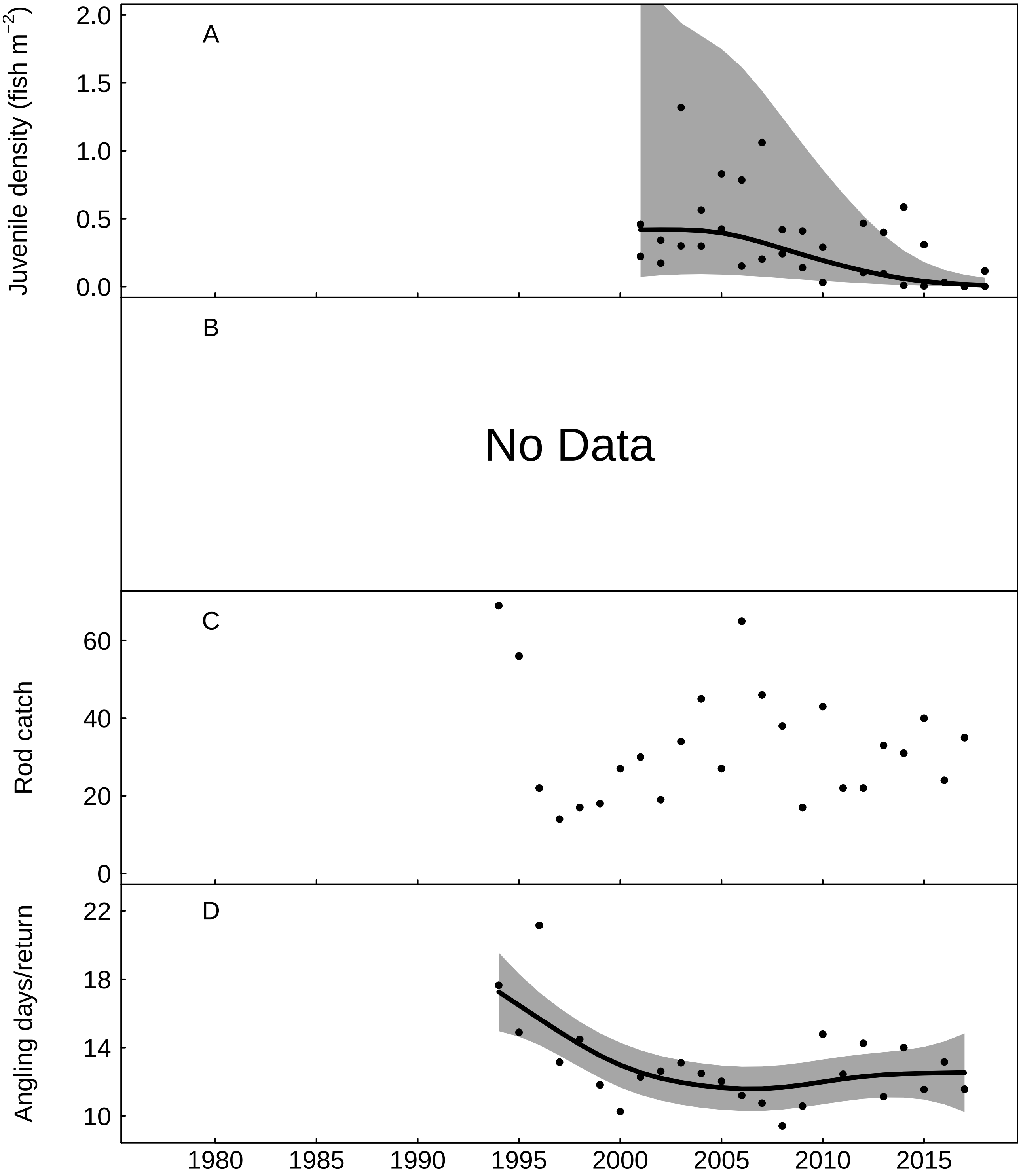


Fig. S49: Trends in the Nevern for A) juvenile salmon density, fitted values from Gaussian additive mixed model, grey areas represent 95% point-wise confidence bands for the smoother, B) No Data, C) rod catch, and D) angling effort measured as number of days fished per licence return, fitted values from Gaussian additive model, grey areas represent 95% point-wise confidence bands for the smoother.

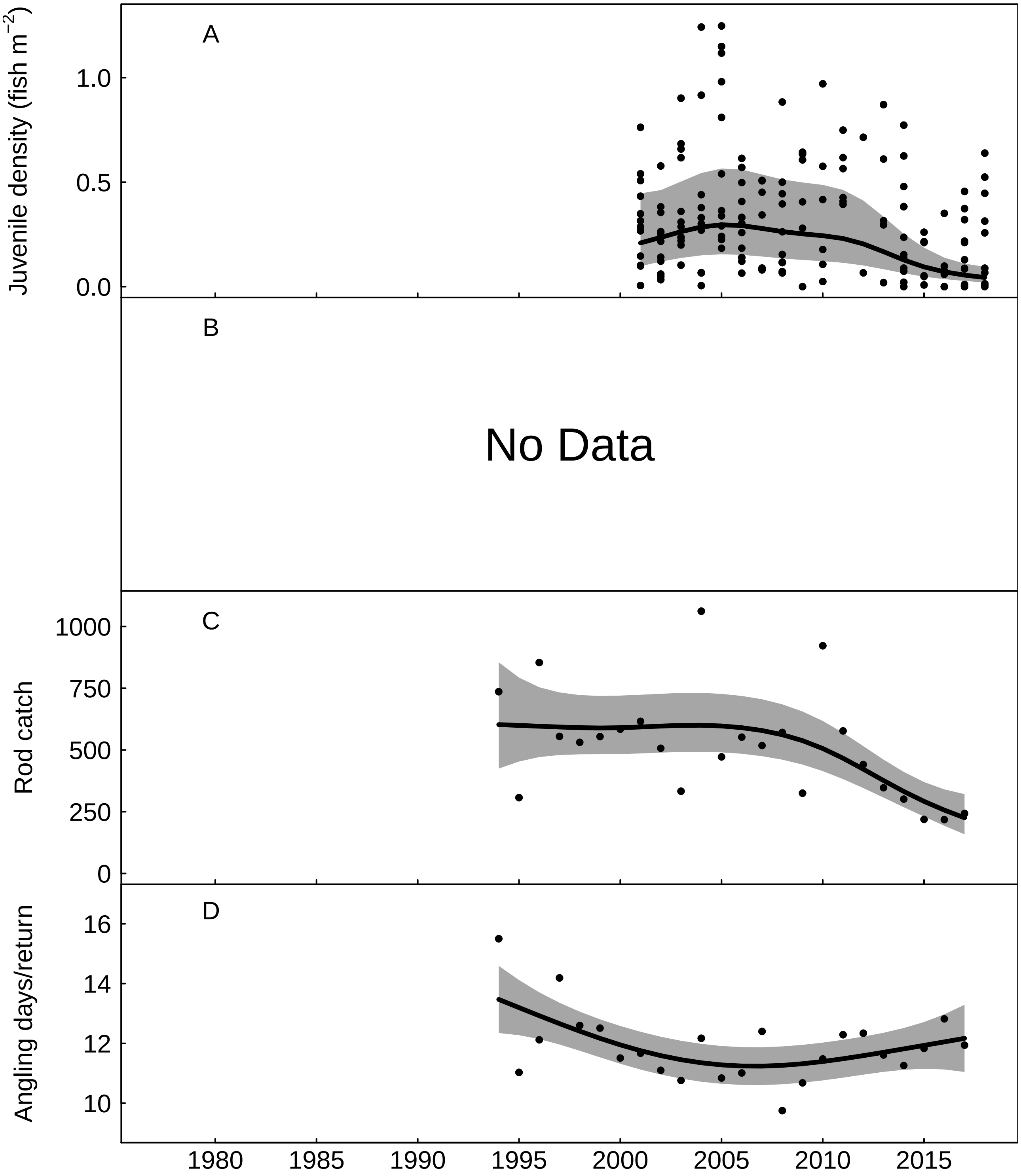


Fig. S50: Trends in the Teifi for A) juvenile salmon density, fitted values from Gaussian additive mixed model, grey areas represent 95% point-wise confidence bands for the smoother, B) No Data, C) rod catch, fitted values from negative binomial generalized additive model, grey areas represent 95% point-wise confidence bands for the smoother, and D) angling effort measured as number of days fished per licence return, fitted values from Gaussian additive model, grey areas represent 95% point-wise confidence bands for the smoother.

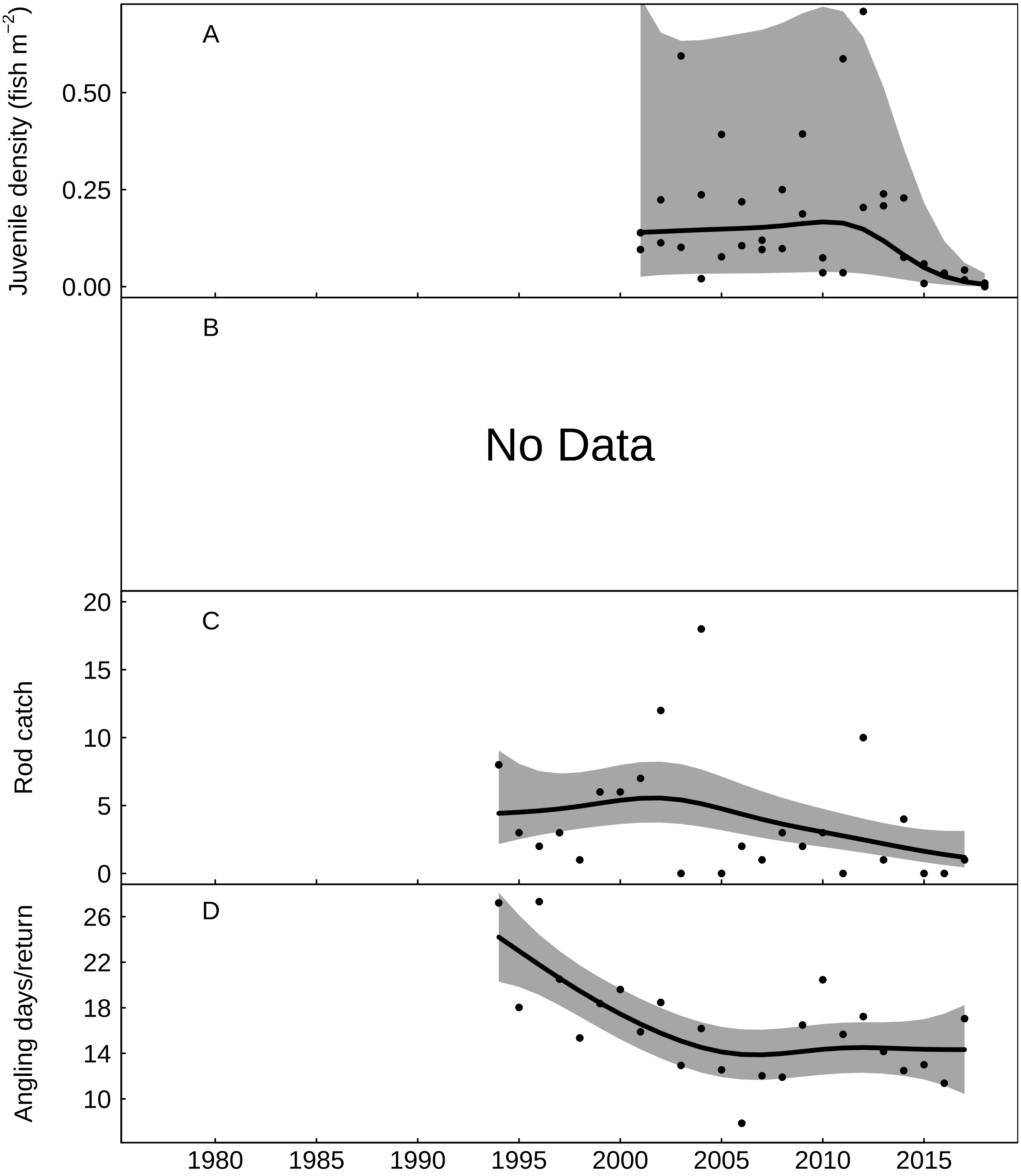


Fig. S51: Trends in the Aeron for A) juvenile salmon density, fitted values from Gaussian additive mixed model, grey areas represent 95% point-wise confidence bands for the smoother, B) No Data, C) rod catch, fitted values from negative binomial generalized additive model, grey areas represent 95% point-wise confidence bands for the smoother, and D) angling effort measured as number of days fished per licence return, fitted values from Gaussian additive model, grey areas represent 95% point-wise confidence bands for the smoother.

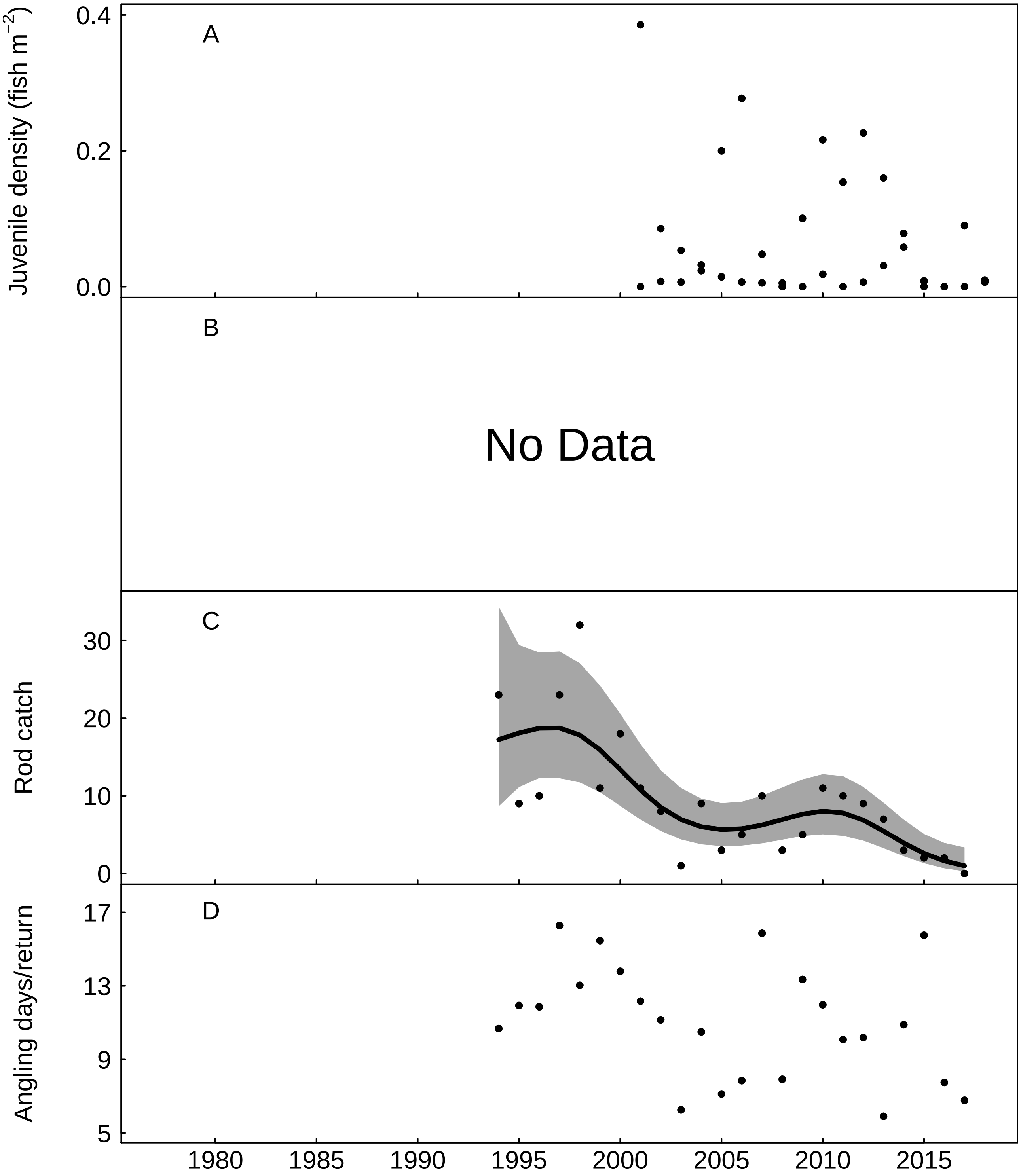


Fig. S52: Trends in the Ystwyth for A) juvenile salmon density, B) No Data, C) rod catch, fitted values from negative binomial generalized additive model, grey areas represent 95% point-wise confidence bands for the smoother, and D) angling effort measured as number of days fished per licence return.

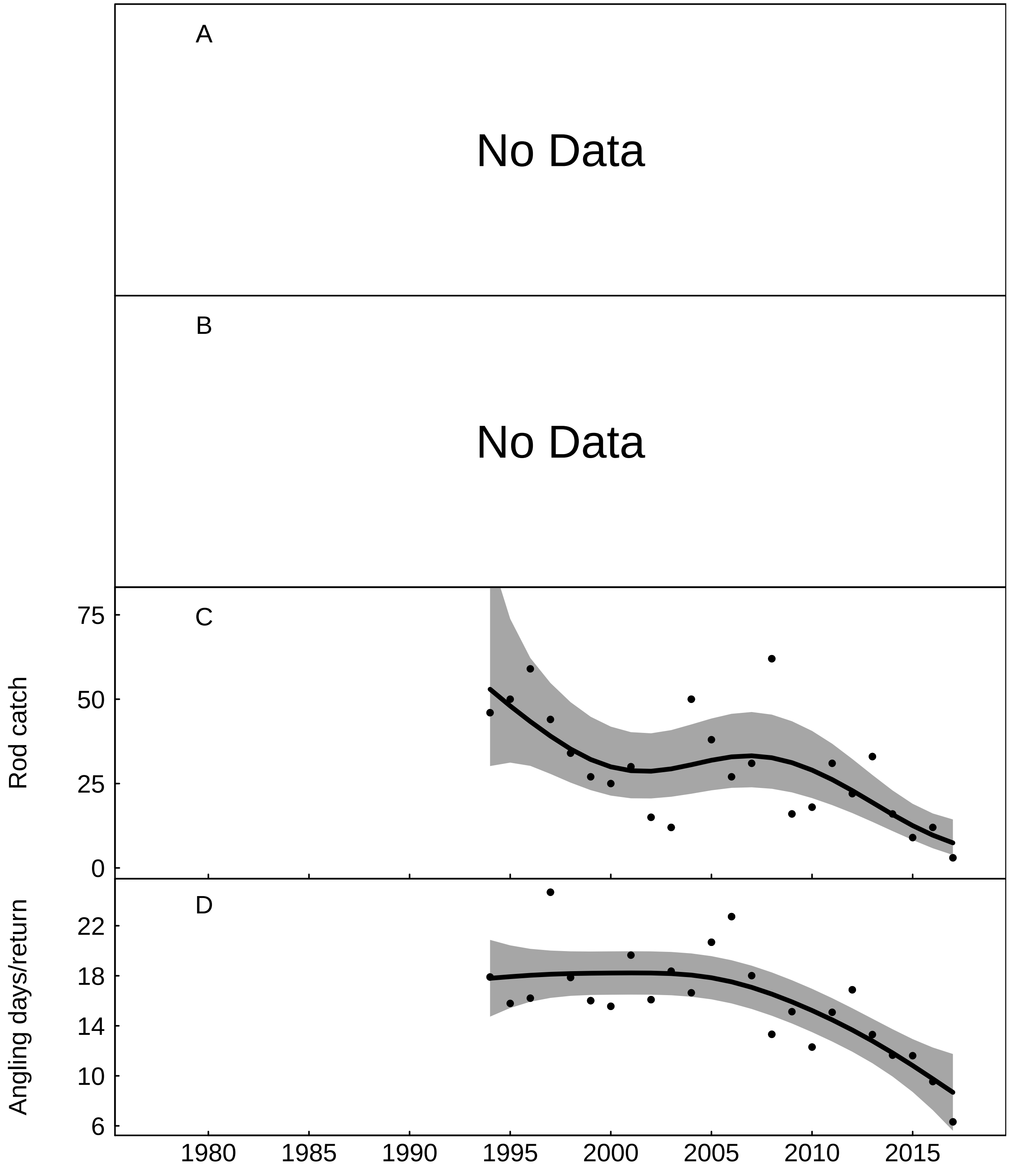


Fig. S53: Trends in the Rheidol for A) No Data, B) No Data, C) rod catch, fitted values from negative binomial generalized additive model, grey areas represent 95% point-wise confidence bands for the smoother, and D) angling effort measured as number of days fished per licence return, fitted values from Gaussian additive model, grey areas represent 95% point-wise confidence bands for the smoother.

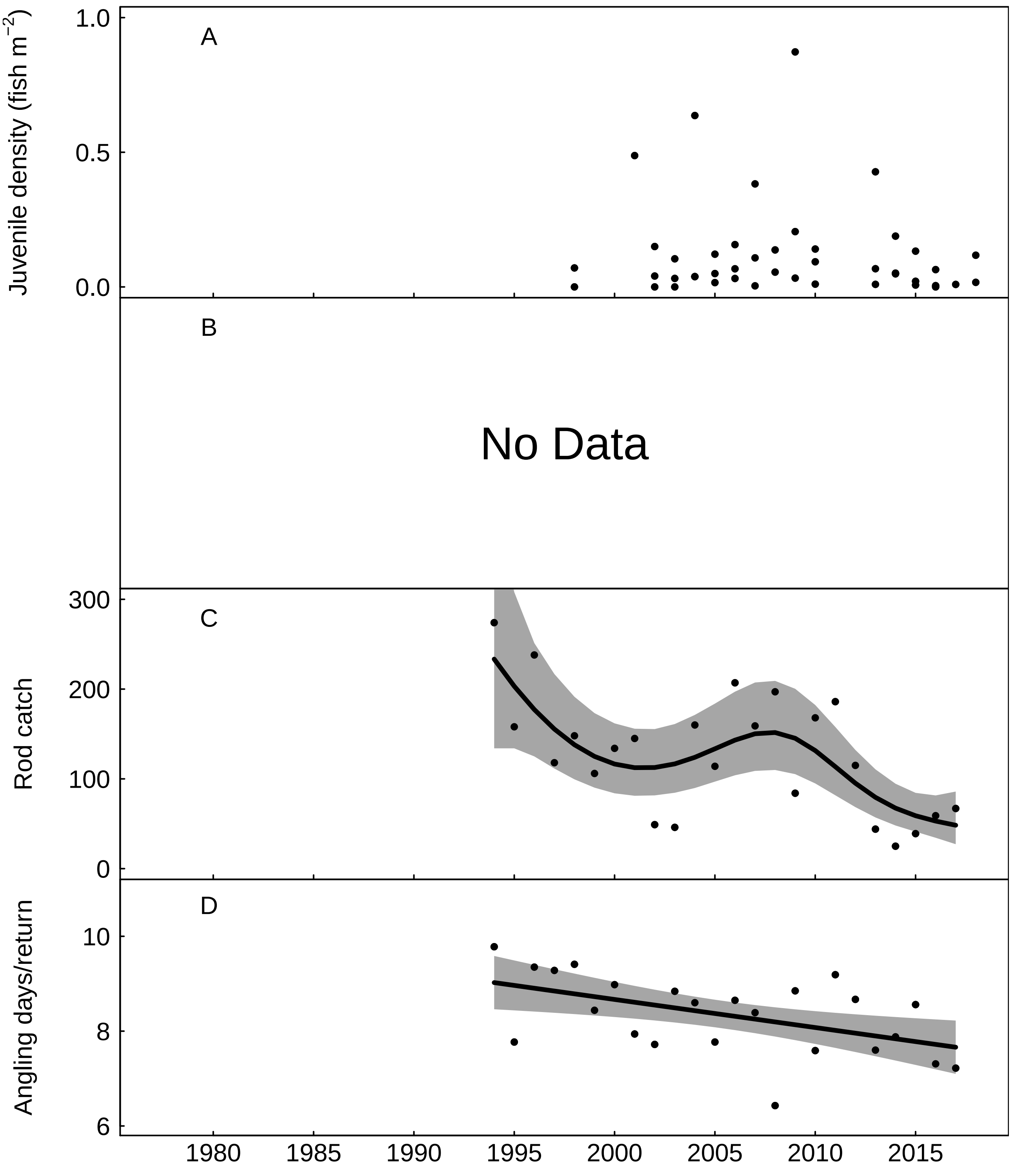


Fig. S54: Trends in the Dyfi for A) juvenile salmon density, B) No Data, C) rod catch, fitted values from negative binomial generalized additive model, grey areas represent 95% point-wise confidence bands for the smoother, and D) angling effort measured as number of days fished per licence return, fitted values from Gaussian additive model, grey areas represent 95% point-wise confidence bands for the smoother.

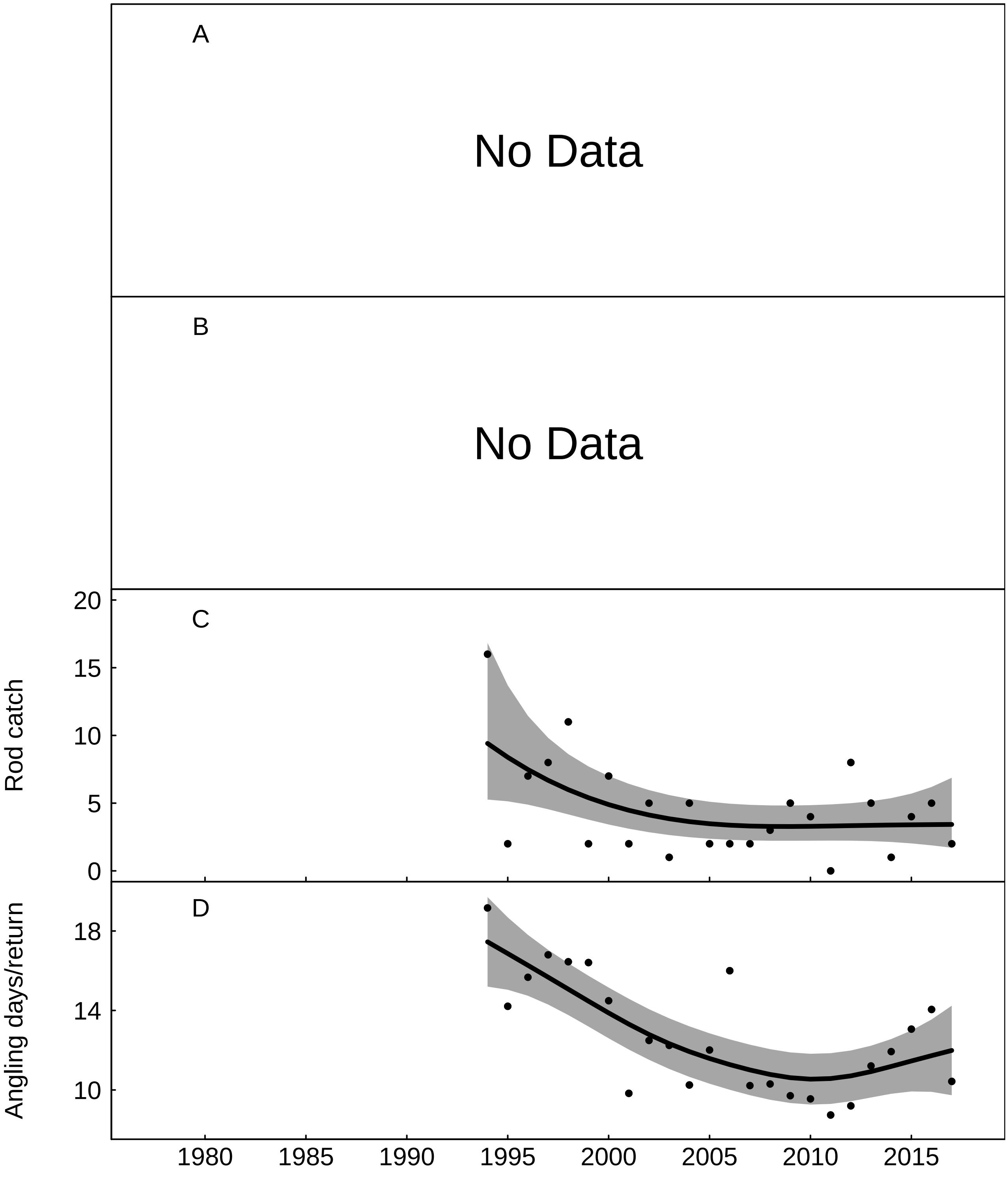


Fig. S55: Trends in the Dysynni for A) No Data, B) No Data, C) rod catch, fitted values from negative binomial generalized additive model, grey areas represent 95% point-wise confidence bands for the smoother, and D) angling effort measured as number of days fished per licence return, fitted values from Gaussian additive model, grey areas represent 95% point-wise confidence bands for the smoother.

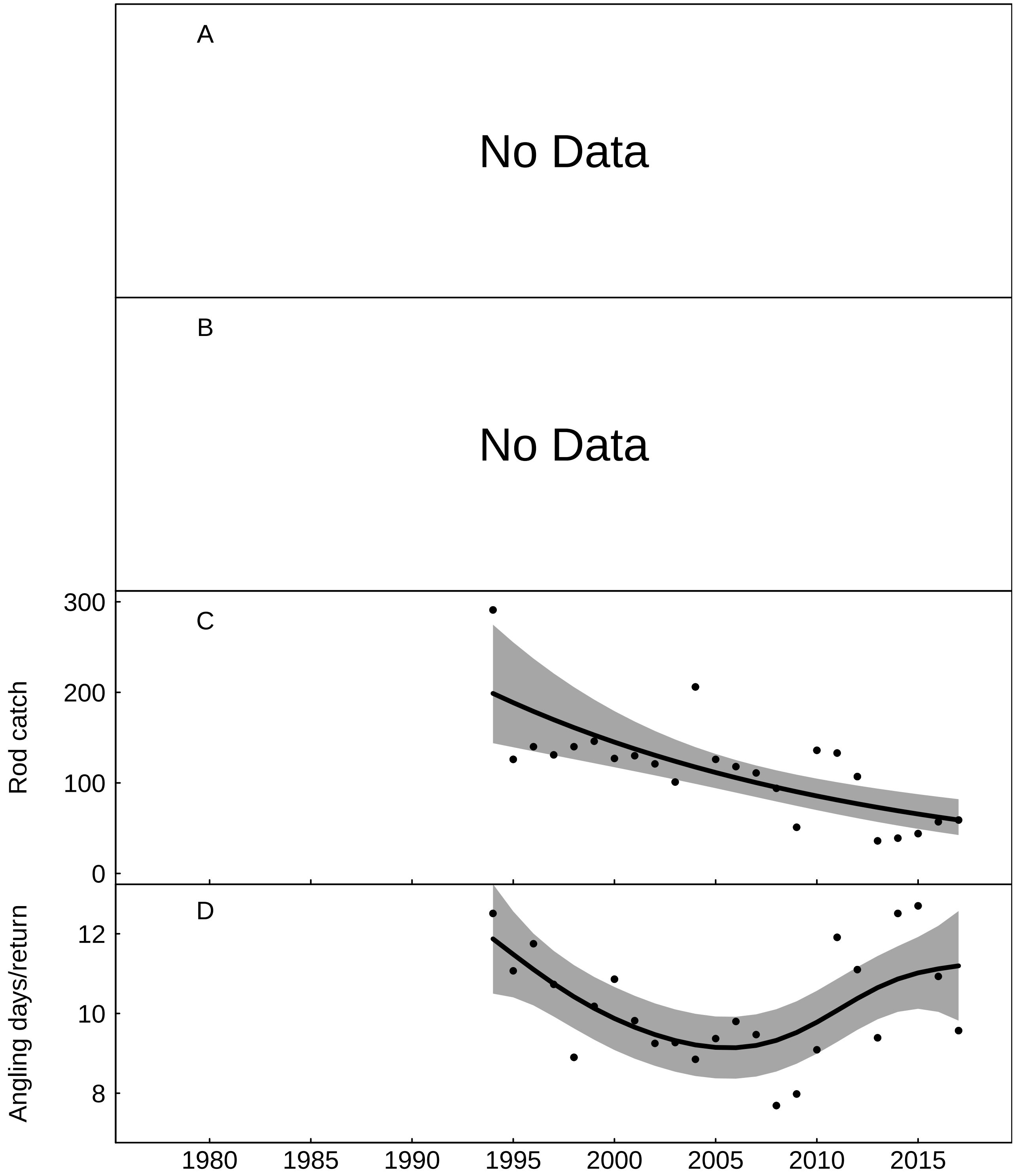


Fig. S56: Trends in the Mawddach for A) No Data, B) No Data, C) rod catch, fitted values from negative binomial generalized additive model, grey areas represent 95% point-wise confidence bands for the smoother, and D) angling effort measured as number of days fished per licence return, fitted values from Gaussian additive model, grey areas represent 95% point-wise confidence bands for the smoother.

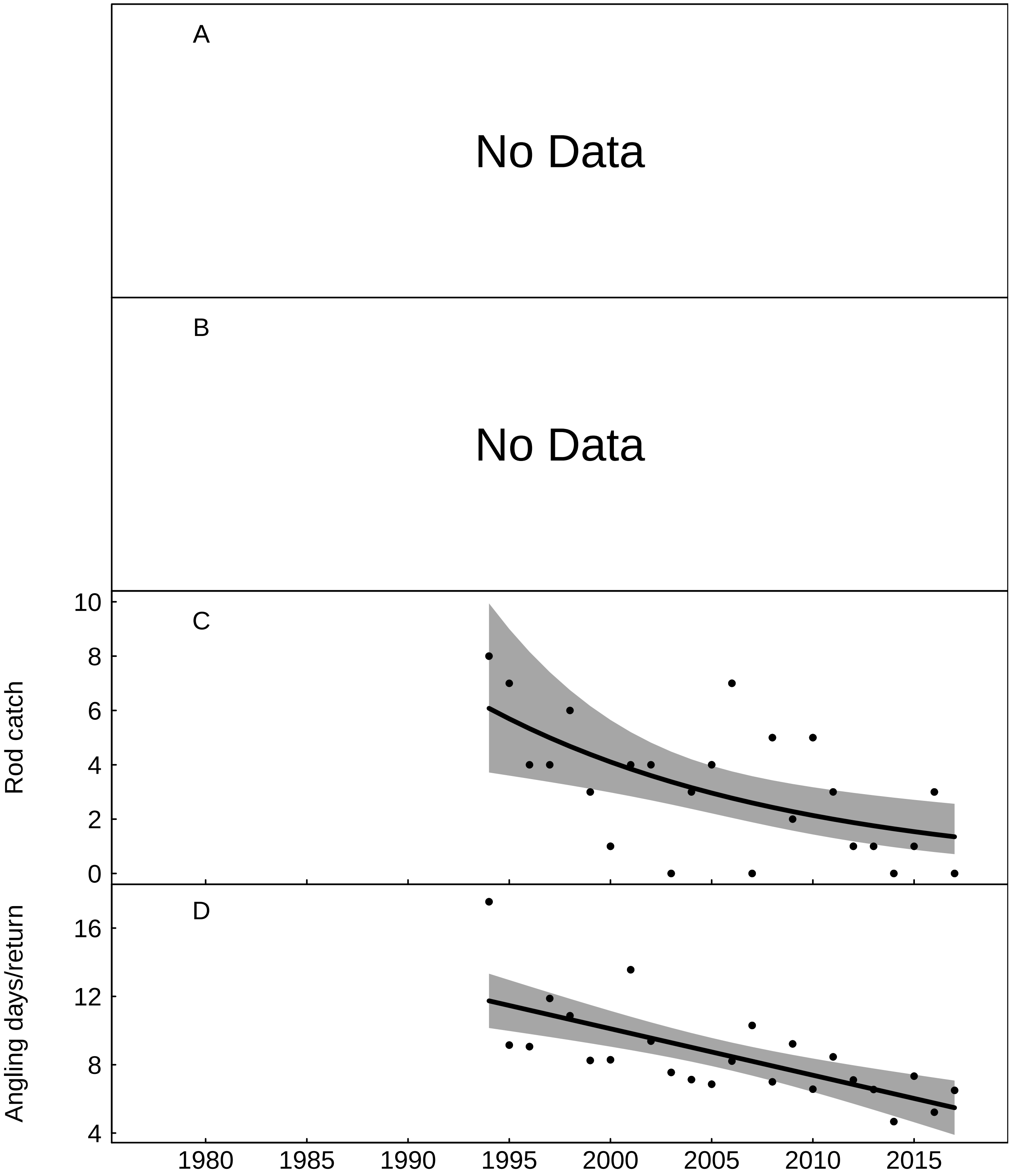


Fig. S57: Trends in the Artro for A) No Data, B) No Data, C) rod catch, fitted values from negative binomial generalized additive model, grey areas represent 95% point-wise confidence bands for the smoother, and D) angling effort measured as number of days fished per licence return, fitted values from Gaussian additive model, grey areas represent 95% point-wise confidence bands for the smoother.

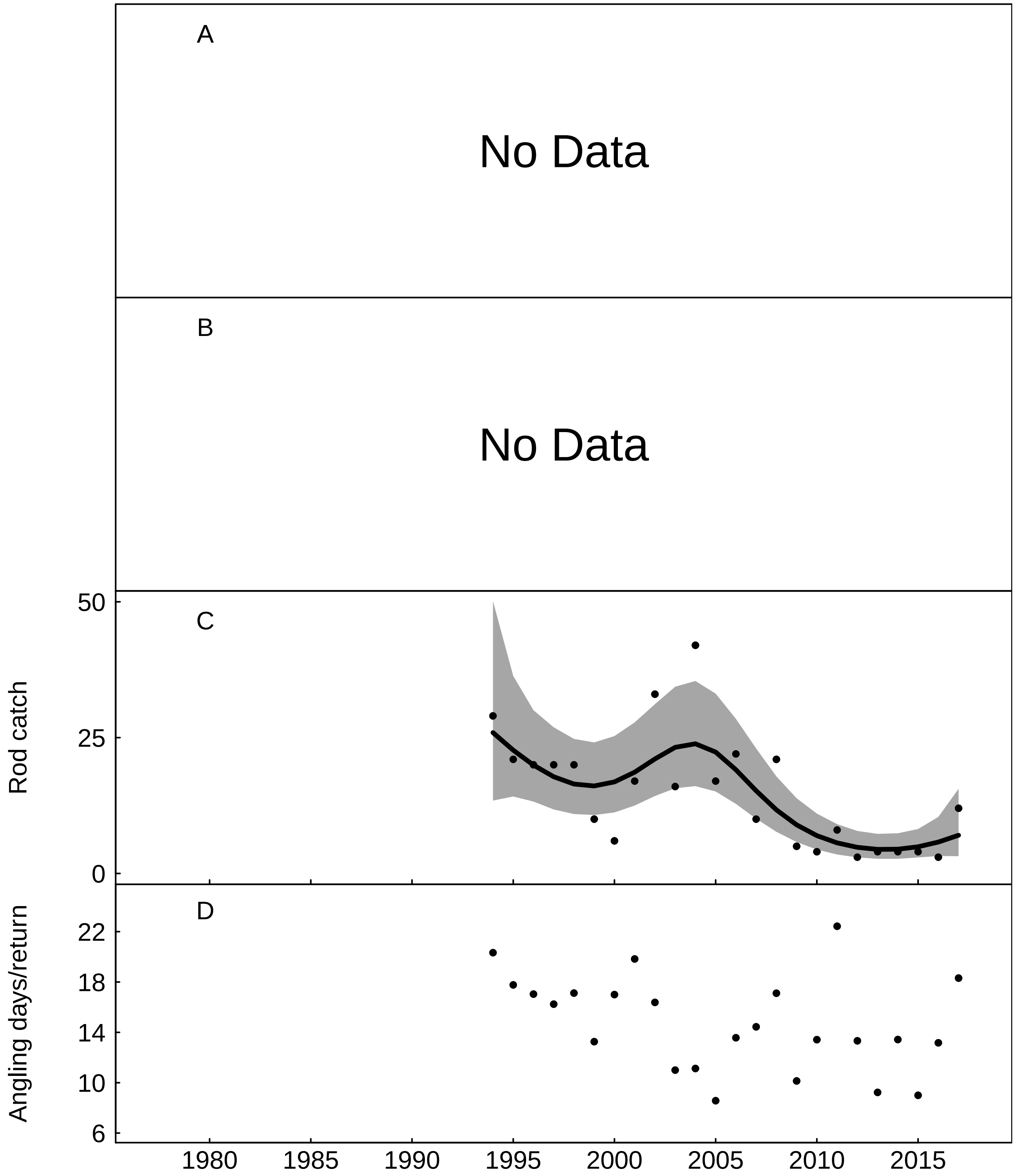


Fig. S58: Trends in the Dwyryd for A) No Data, B) No Data, C) rod catch, fitted values from negative binomial generalized additive model, grey areas represent 95% point-wise confidence bands for the smoother, and D) angling effort measured as number of days fished per licence return.

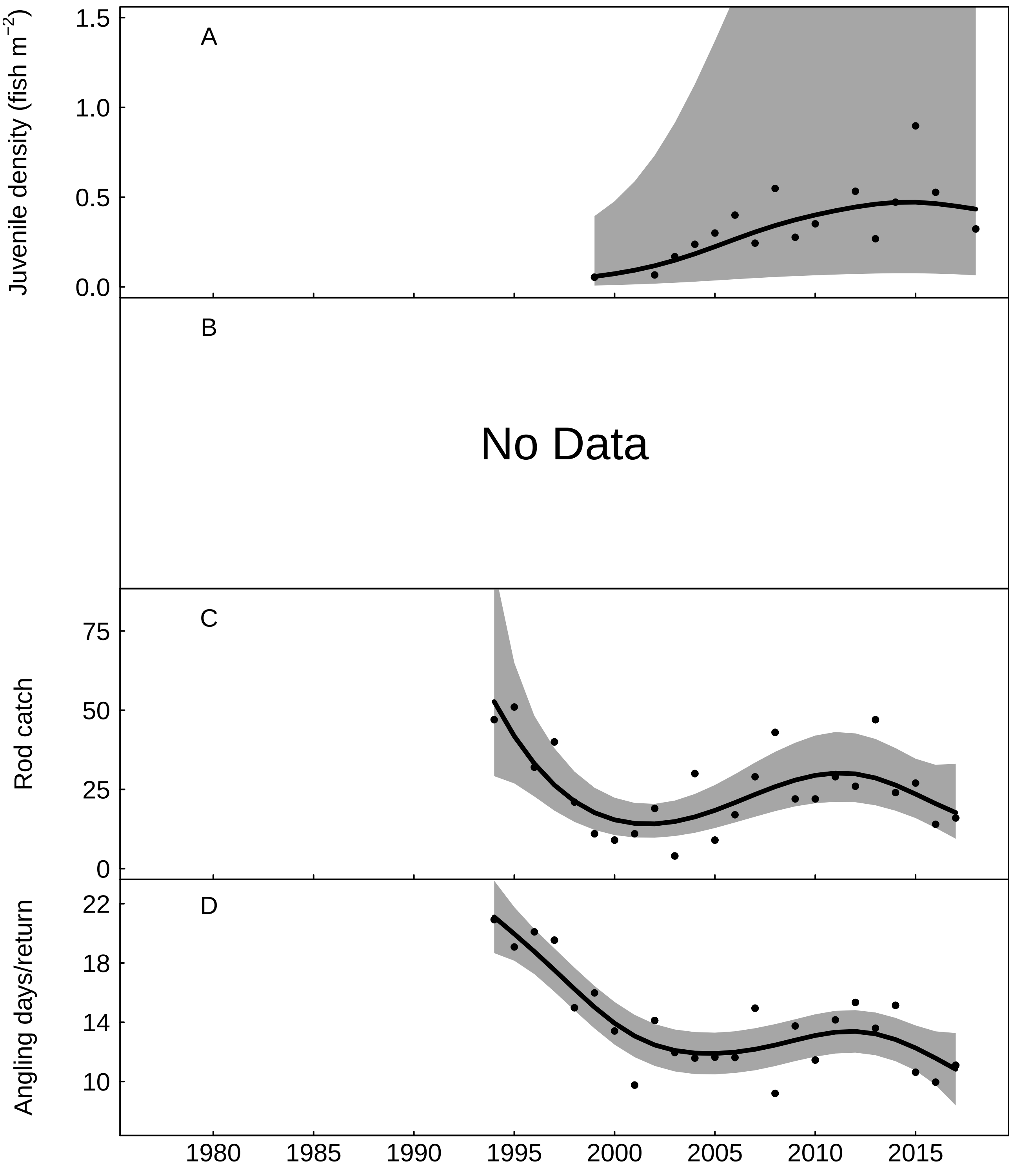


Fig. S59: Trends in the Glaslyn for A) juvenile salmon density, fitted values from Gaussian additive mixed model, grey areas represent 95% point-wise confidence bands for the smoother, B) No Data, C) rod catch, fitted values from negative binomial generalized additive model, grey areas represent 95% point-wise confidence bands for the smoother, and D) angling effort measured as number of days fished per licence return, fitted values from Gaussian additive model, grey areas represent 95% point-wise confidence bands for the smoother.

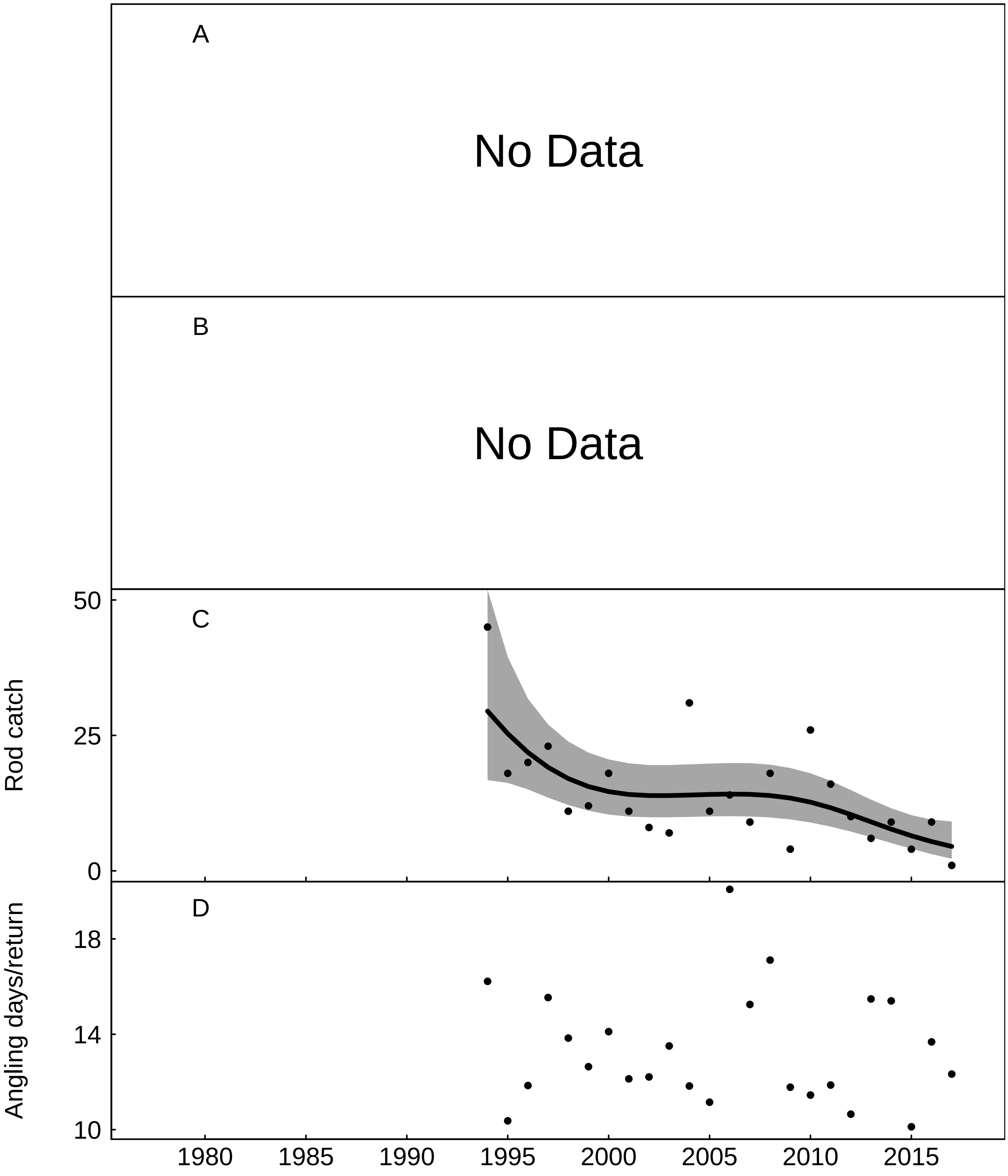


Fig. S60: Trends in the Dwyfawr for A) No Data, B) No Data, C) rod catch, fitted values from negative binomial generalized additive model, grey areas represent 95% point-wise confidence bands for the smoother, and D) angling effort measured as number of days fished per licence return.

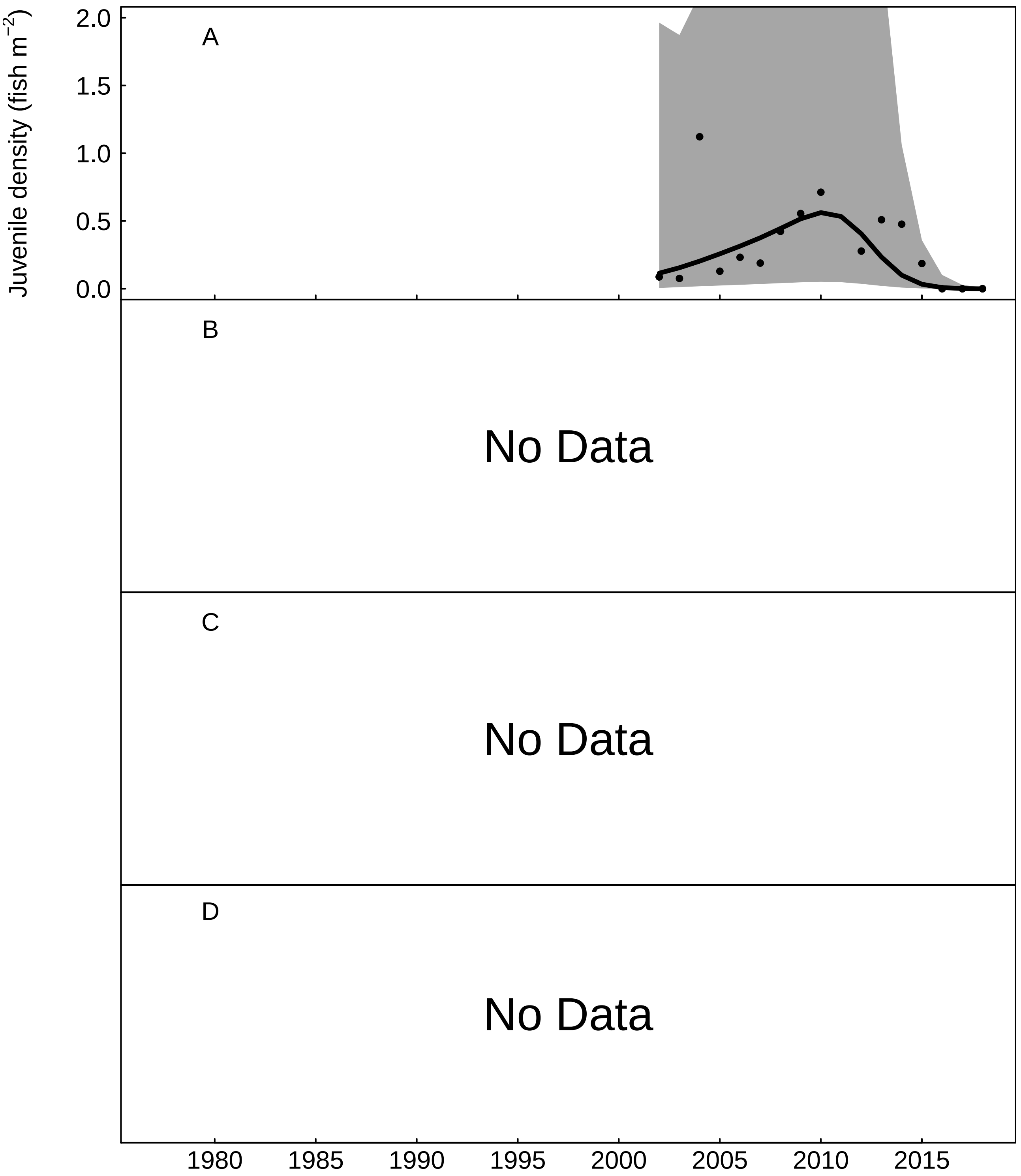


Fig. S61: Trends in the Erch for A) juvenile salmon density, fitted values from Gaussian additive mixed model, grey areas represent 95% point-wise confidence bands for the smoother, B) No Data, C) No Data, and D) No Data.

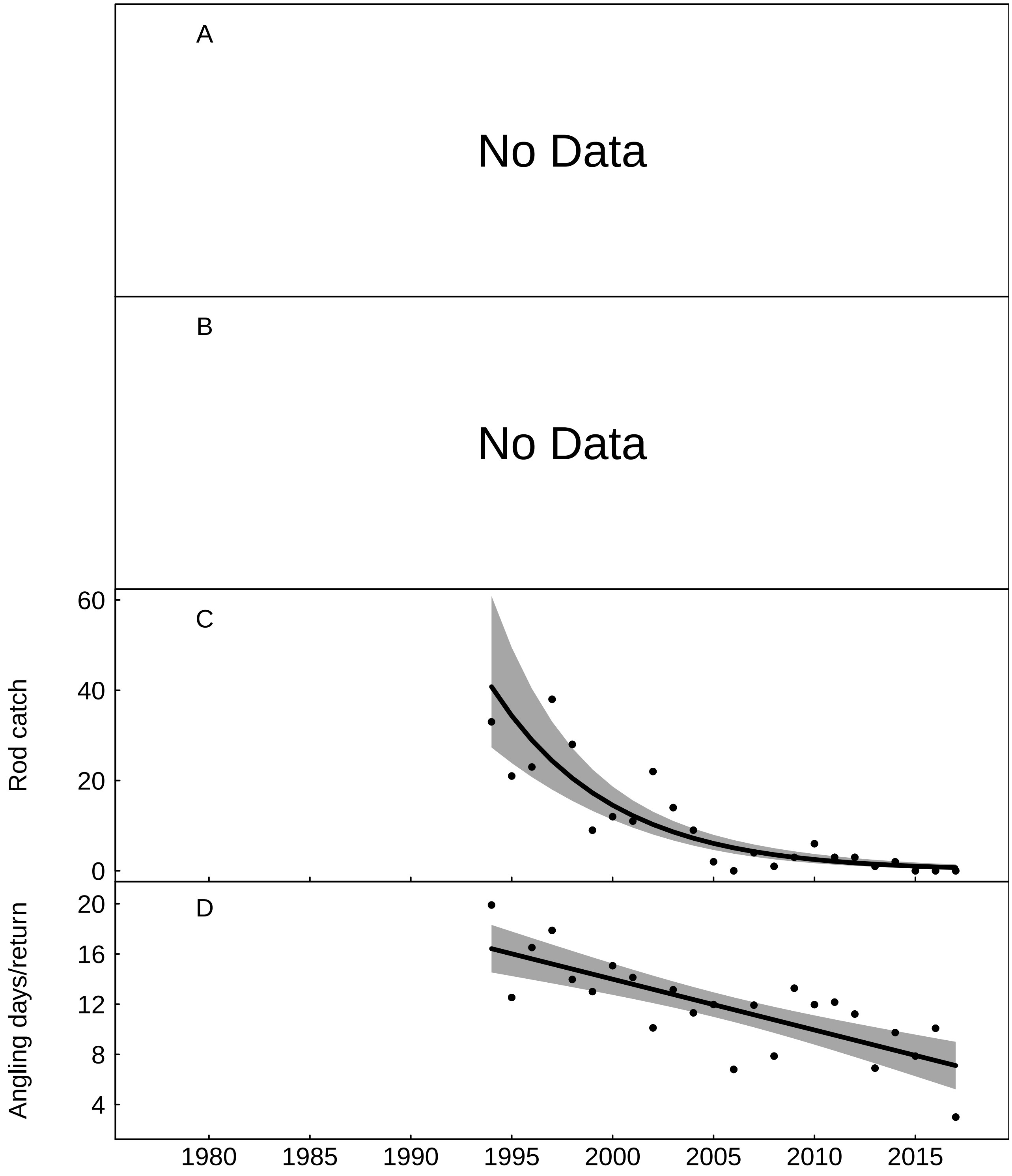


Fig. S62: Trends in the Llyfni for A) No Data, B) No Data, C) rod catch, fitted values from negative binomial generalized additive model, grey areas represent 95% point-wise confidence bands for the smoother, and D) angling effort measured as number of days fished per licence return, fitted values from Gaussian additive model, grey areas represent 95% point-wise confidence bands for the smoother.

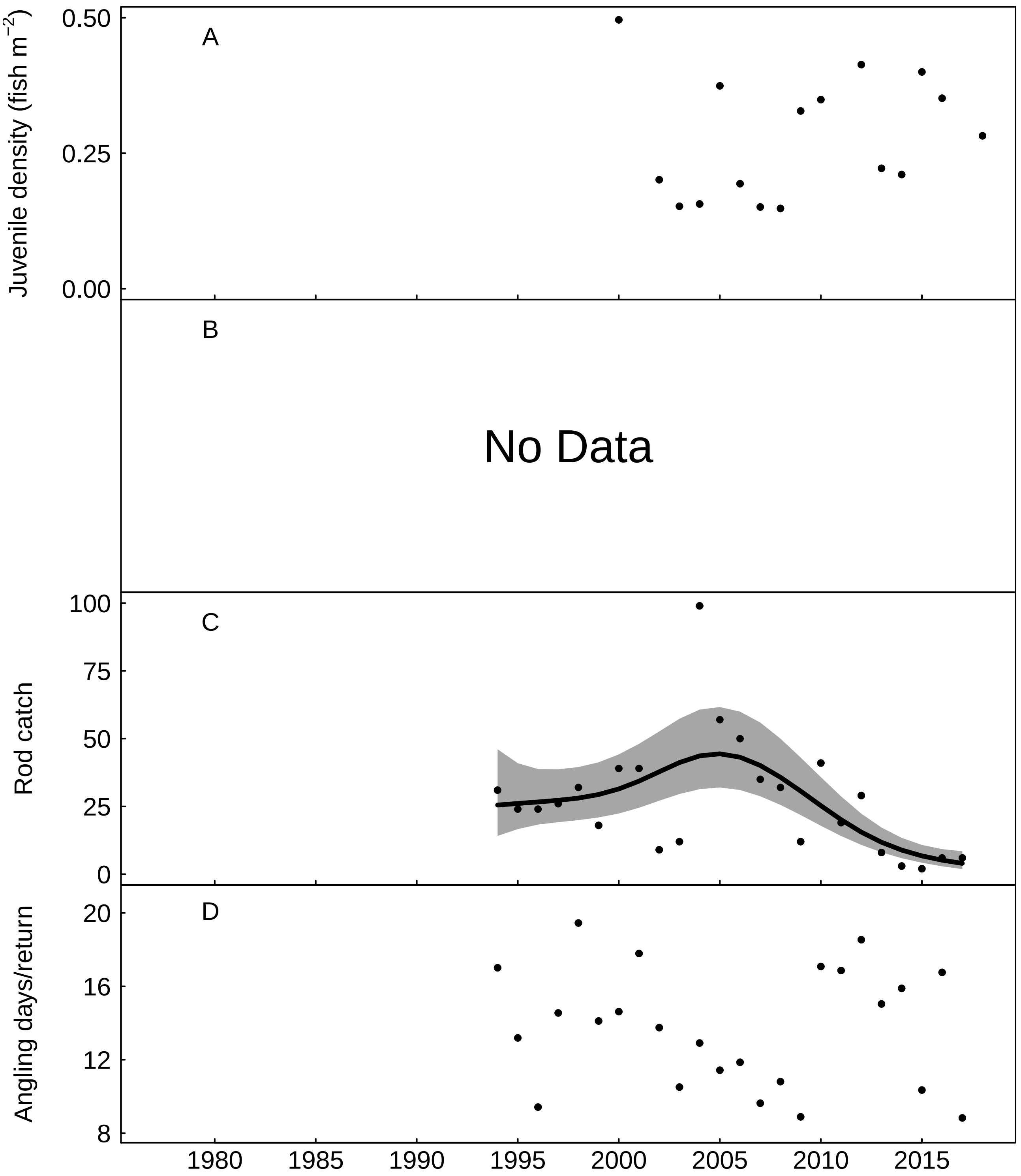


Fig. S63: Trends in the Seiont for A) juvenile salmon density, B) No Data, C) rod catch, fitted values from negative binomial generalized additive model, grey areas represent 95% point-wise confidence bands for the smoother, and D) angling effort measured as number of days fished per licence return.

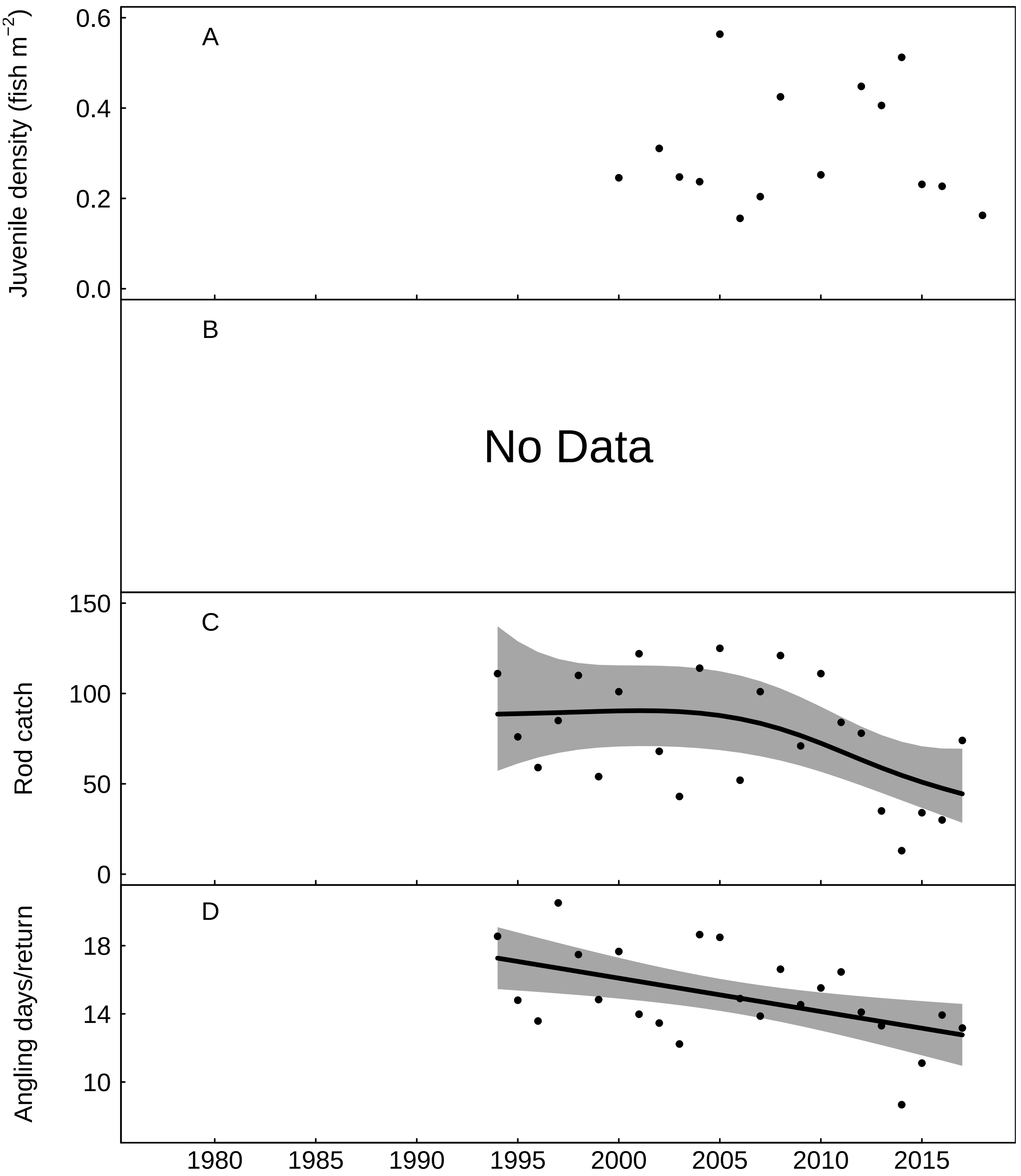


Fig. S64: Trends in the Ogwen for A) juvenile salmon density, B) No Data, C) rod catch, fitted values from negative binomial generalized additive model, grey areas represent 95% point-wise confidence bands for the smoother, and D) angling effort measured as number of days fished per licence return, fitted values from Gaussian additive model, grey areas represent 95% point-wise confidence bands for the smoother.

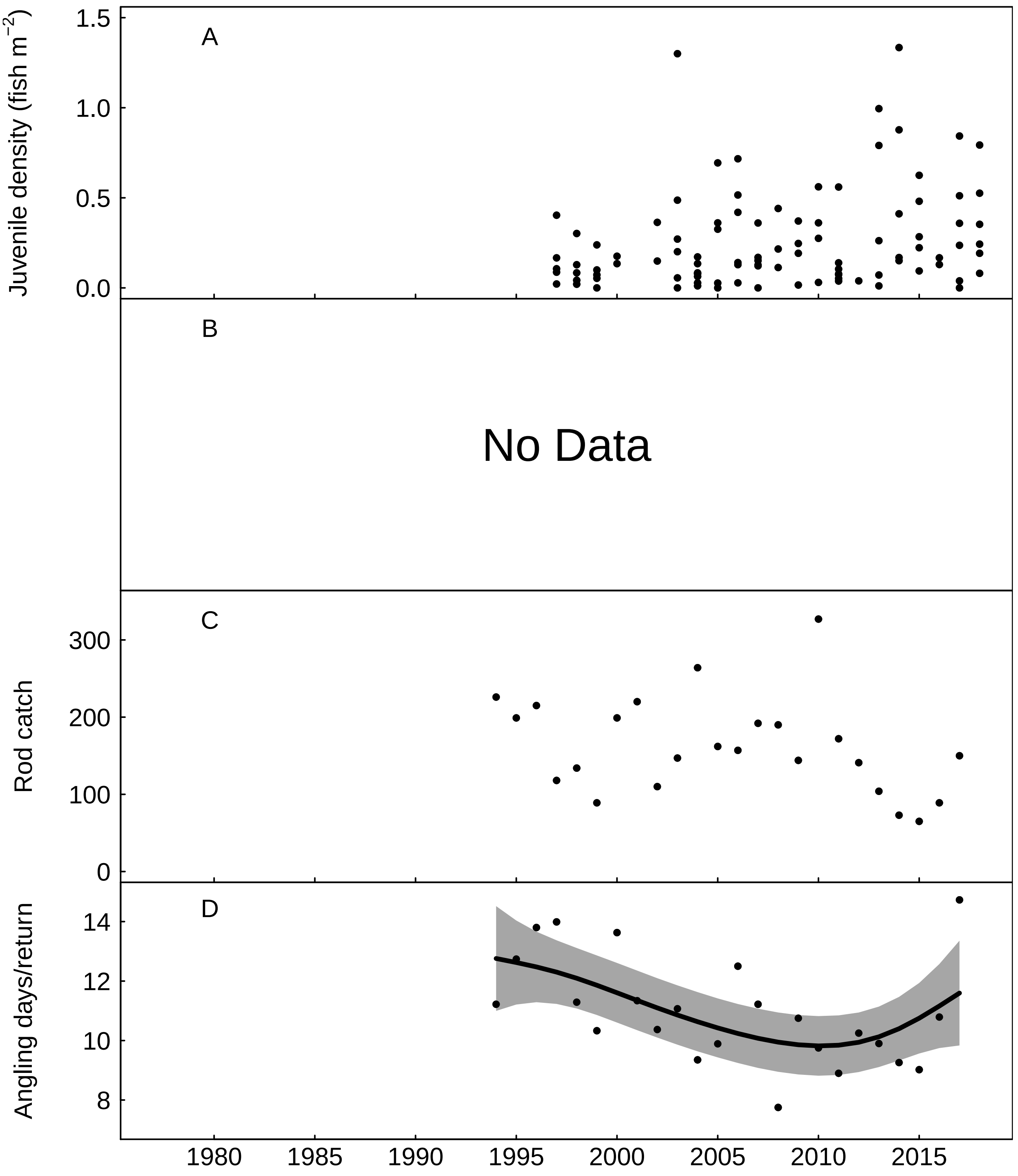


Fig. S65: Trends in the Conway for A) juvenile salmon density, B) No Data, C) rod catch, and D) angling effort measured as number of days fished per licence return, fitted values from Gaussian additive model, grey areas represent 95% point-wise confidence bands for the smoother.

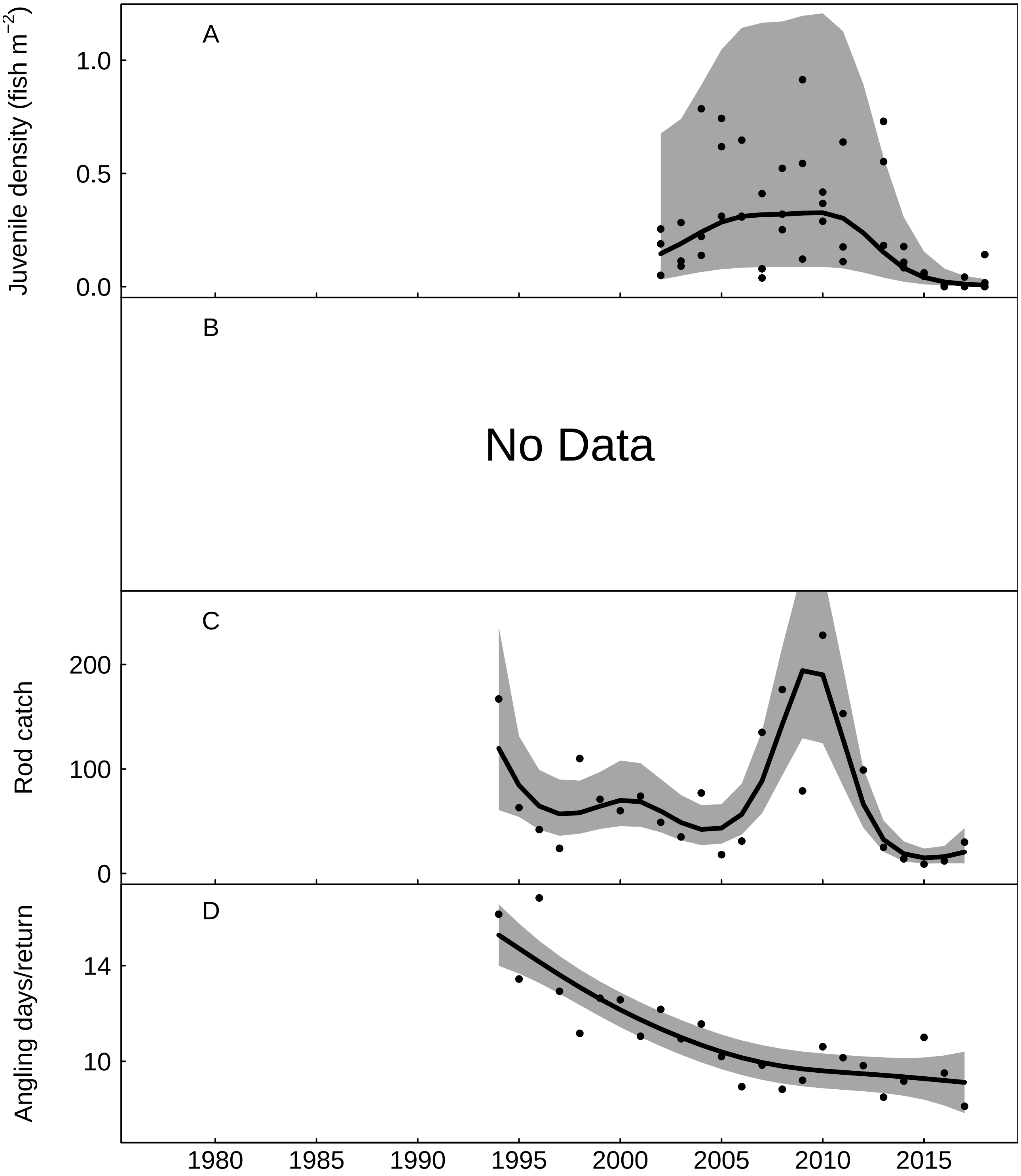


Fig. S66: Trends in the Clwyd for A) juvenile salmon density, fitted values from Gaussian additive mixed model, grey areas represent 95% point-wise confidence bands for the smoother, B) No Data, C) rod catch, fitted values from negative binomial generalized additive model, grey areas represent 95% point-wise confidence bands for the smoother, and D) angling effort measured as number of days fished per licence return, fitted values from Gaussian additive model, grey areas represent 95% point-wise confidence bands for the smoother.

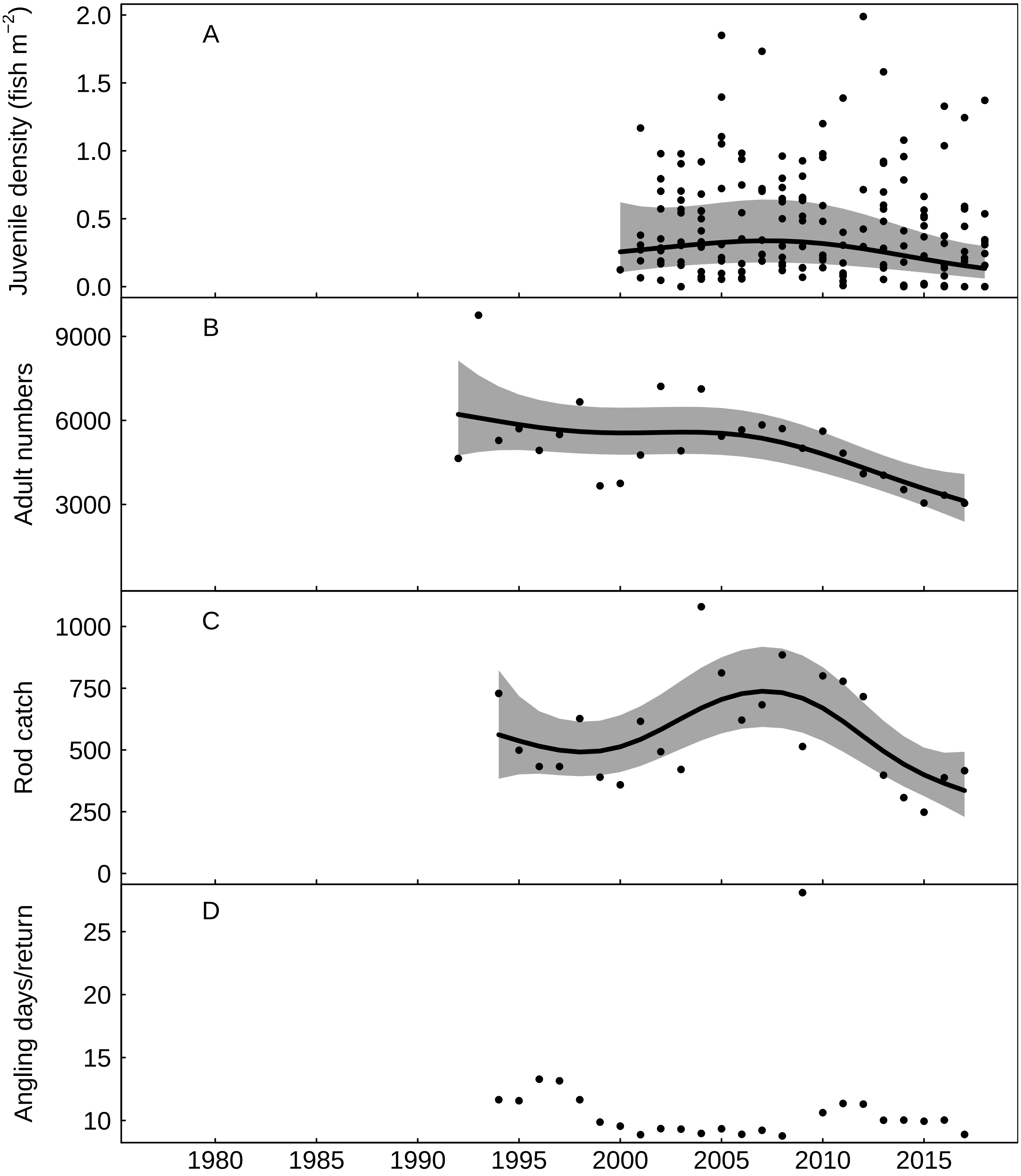


Fig. S67: Trends in the Dee for A) juvenile salmon density, fitted values from Gaussian additive mixed model, grey areas represent 95% point-wise confidence bands for the smoother, B) returning adult numbers, fitted values from negative binomial generalized additive model, grey areas represent 95% point-wise confidence bands for the smoother, C) rod catch, fitted values from negative binomial generalized additive model, grey areas represent 95% point-wise confidence bands for the smoother, and D) angling effort measured as number of days fished per licence return.

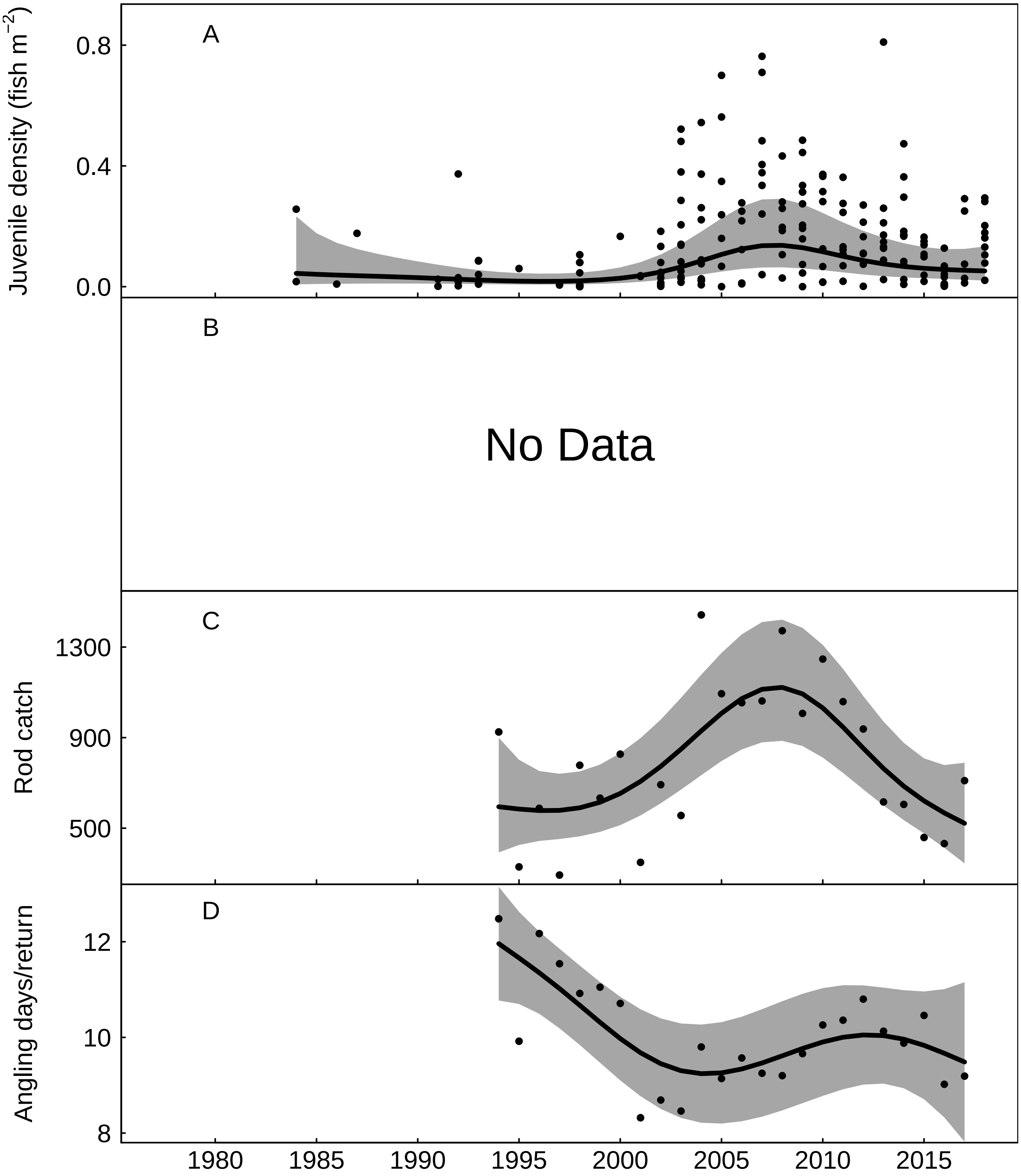


Fig. S68: Trends in the River Ribble for A) juvenile salmon density, fitted values from Gaussian additive mixed model, grey areas represent 95% point-wise confidence bands for the smoother, B) No Data, C) rod catch, fitted values from negative binomial generalized additive model, grey areas represent 95% point-wise confidence bands for the smoother, and D) angling effort measured as number of days fished per licence return, fitted values from Gaussian additive model, grey areas represent 95% point-wise confidence bands for the smoother.

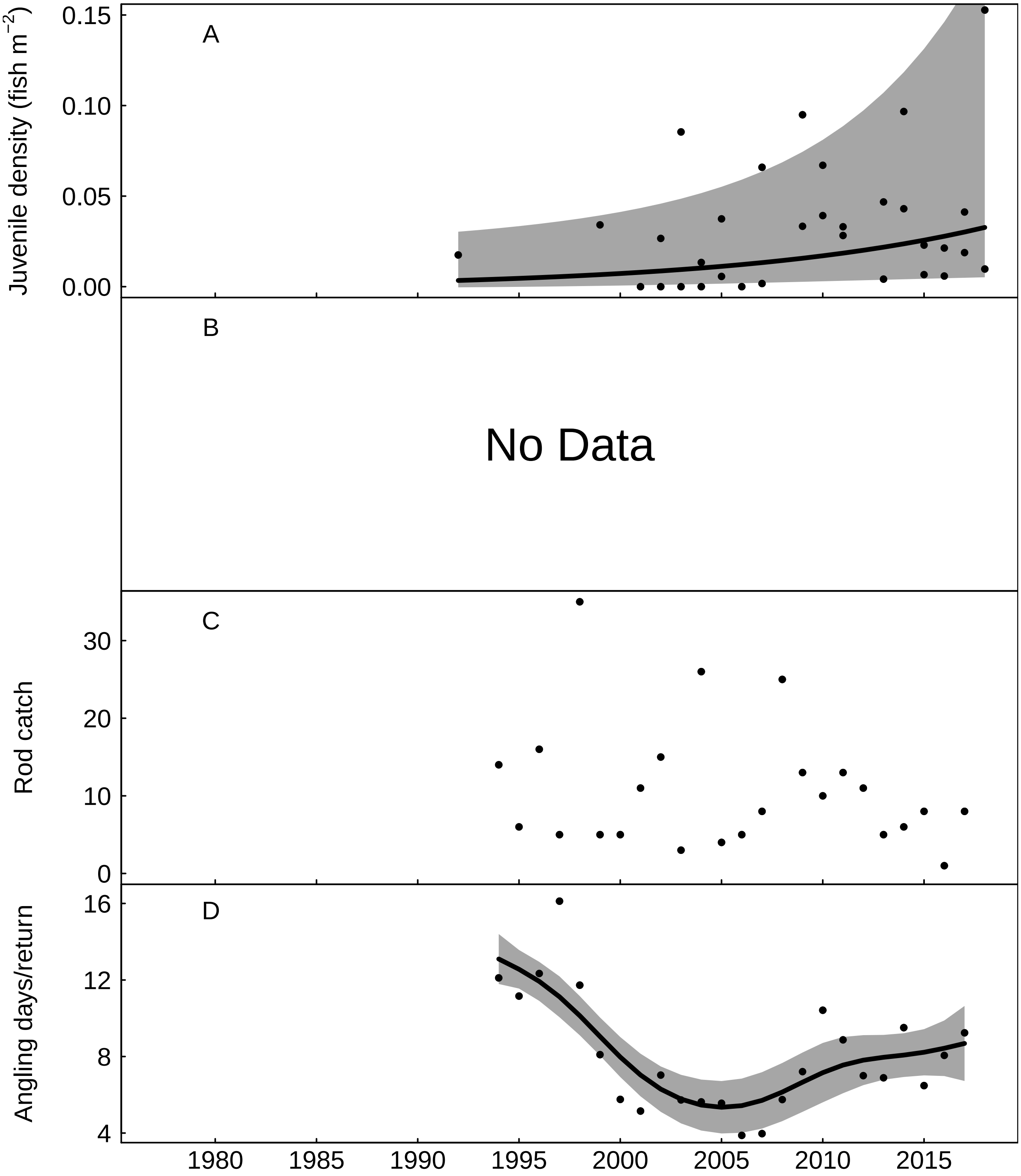


Fig. S69: Trends in the River Wyre for A) juvenile salmon density, fitted values from Gaussian additive mixed model, grey areas represent 95% point-wise confidence bands for the smoother, B) No Data, C) rod catch, and D) angling effort measured as number of days fished per licence return, fitted values from Gaussian additive model, grey areas represent 95% point-wise confidence bands for the smoother.

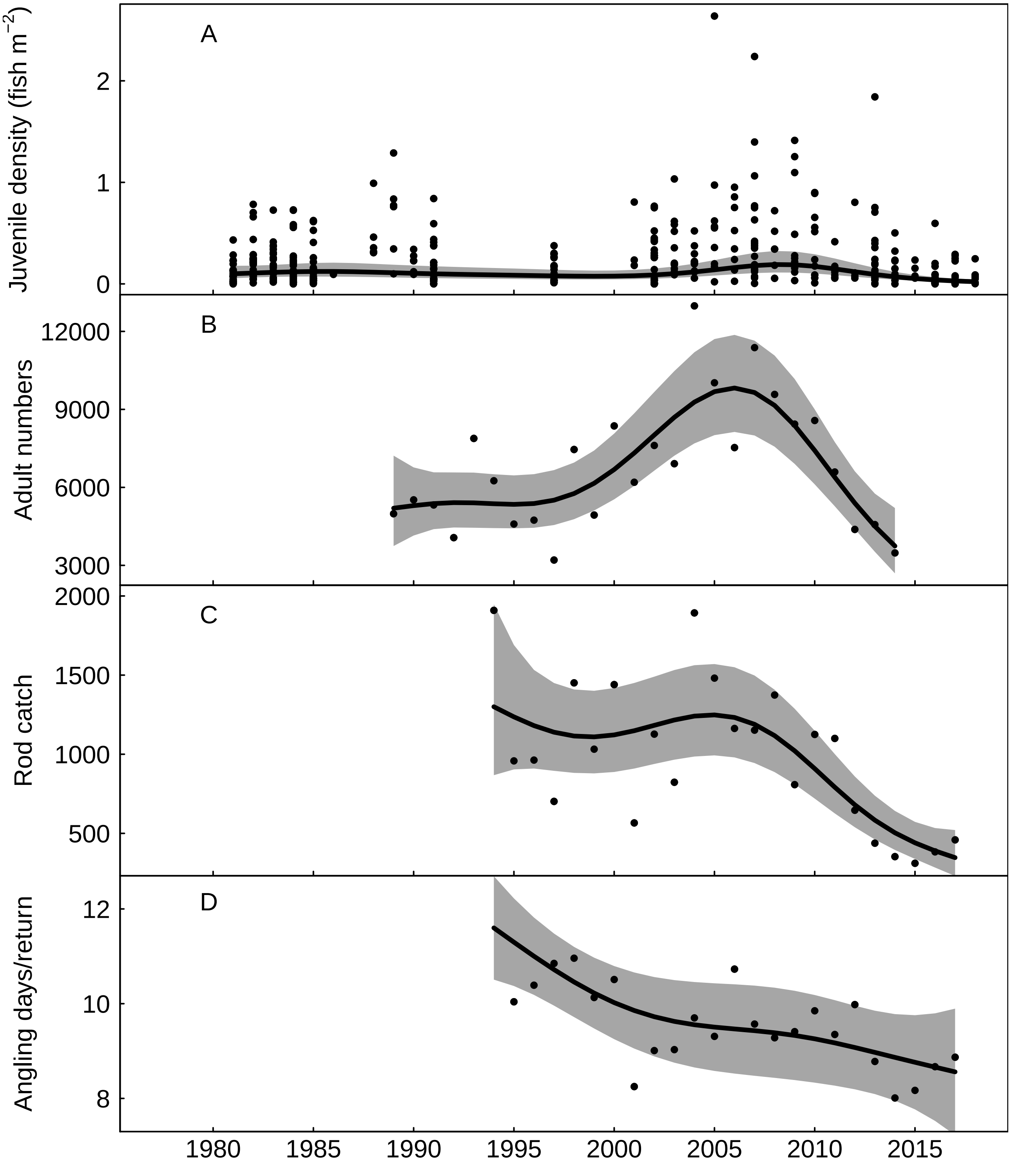


Fig. S70: Trends in the River Lune for A) juvenile salmon density, fitted values from Gaussian additive mixed model, grey areas represent 95% point-wise confidence bands for the smoother, B) returning adult numbers, fitted values from negative binomial generalized additive model, grey areas represent 95% point-wise confidence bands for the smoother, C) rod catch, fitted values from negative binomial generalized additive model, grey areas represent 95% point-wise confidence bands for the smoother, and D) angling effort measured as number of days fished per licence return, fitted values from Gaussian additive model, grey areas represent 95% point-wise confidence bands for the smoother.

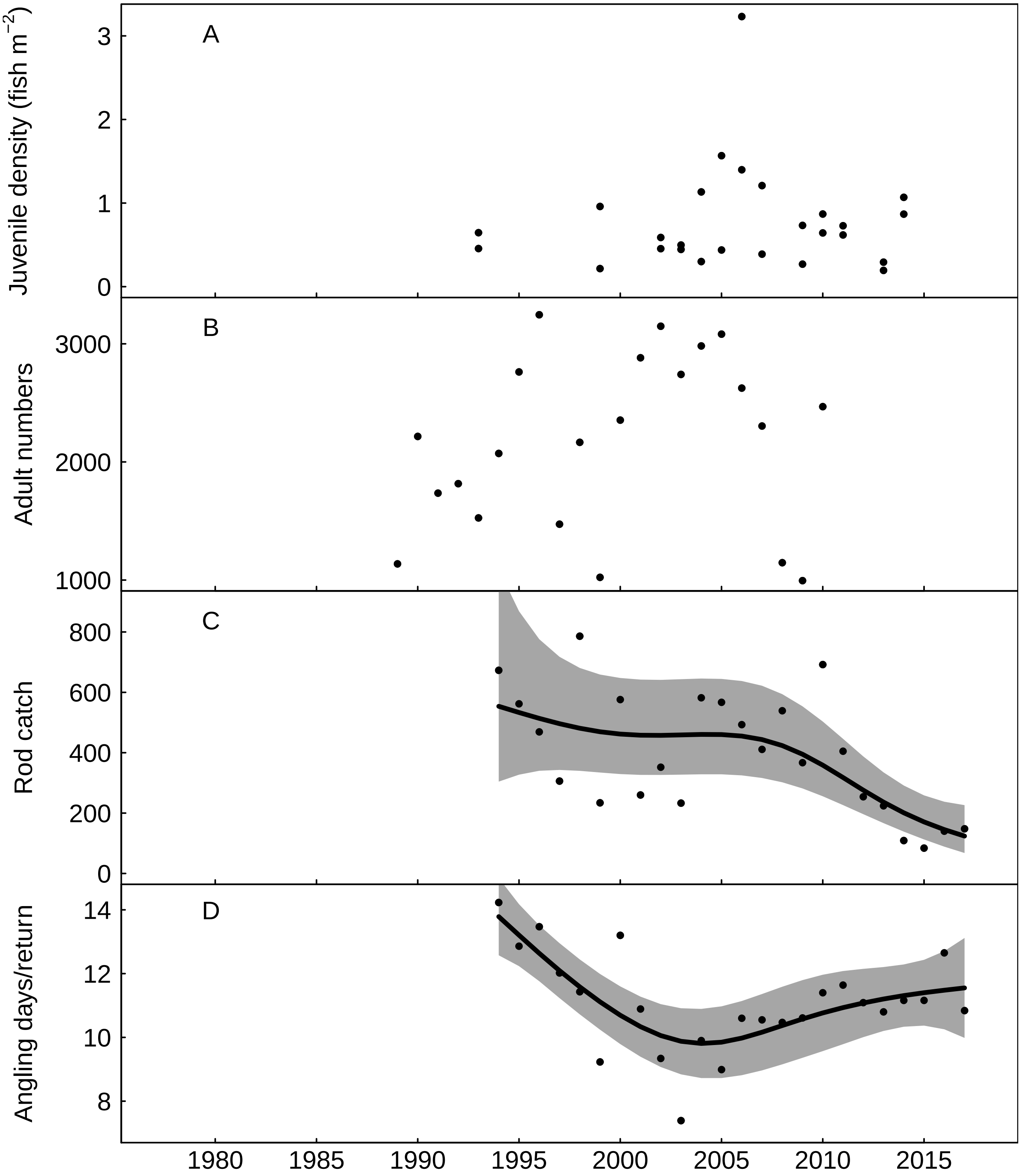


Fig. S71: Trends in the River Kent for A) juvenile salmon density, B) returning adult numbers, C) rod catch, fitted values from negative binomial generalized additive model, grey areas represent 95% point-wise confidence bands for the smoother, and D) angling effort measured as number of days fished per licence return, fitted values from Gaussian additive model, grey areas represent 95% point-wise confidence bands for the smoother.

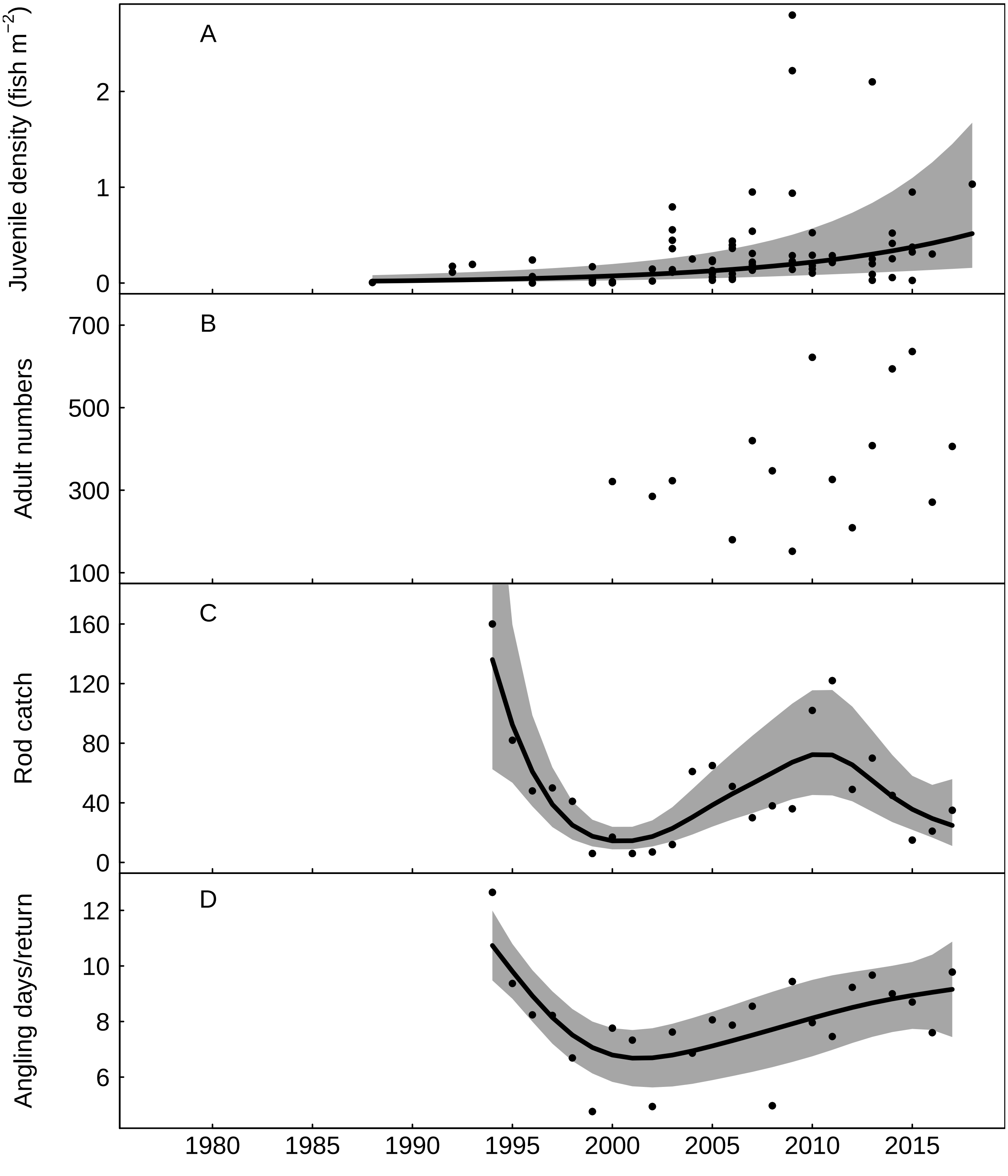


Fig. S72: Trends in the River Leven for A) juvenile salmon density, fitted values from Gaussian additive mixed model, grey areas represent 95% point-wise confidence bands for the smoother, B) returning adult numbers, C) rod catch, fitted values from negative binomial generalized additive model, grey areas represent 95% point-wise confidence bands for the smoother, and D) angling effort measured as number of days fished per licence return, fitted values from Gaussian additive model, grey areas represent 95% point-wise confidence bands for the smoother.

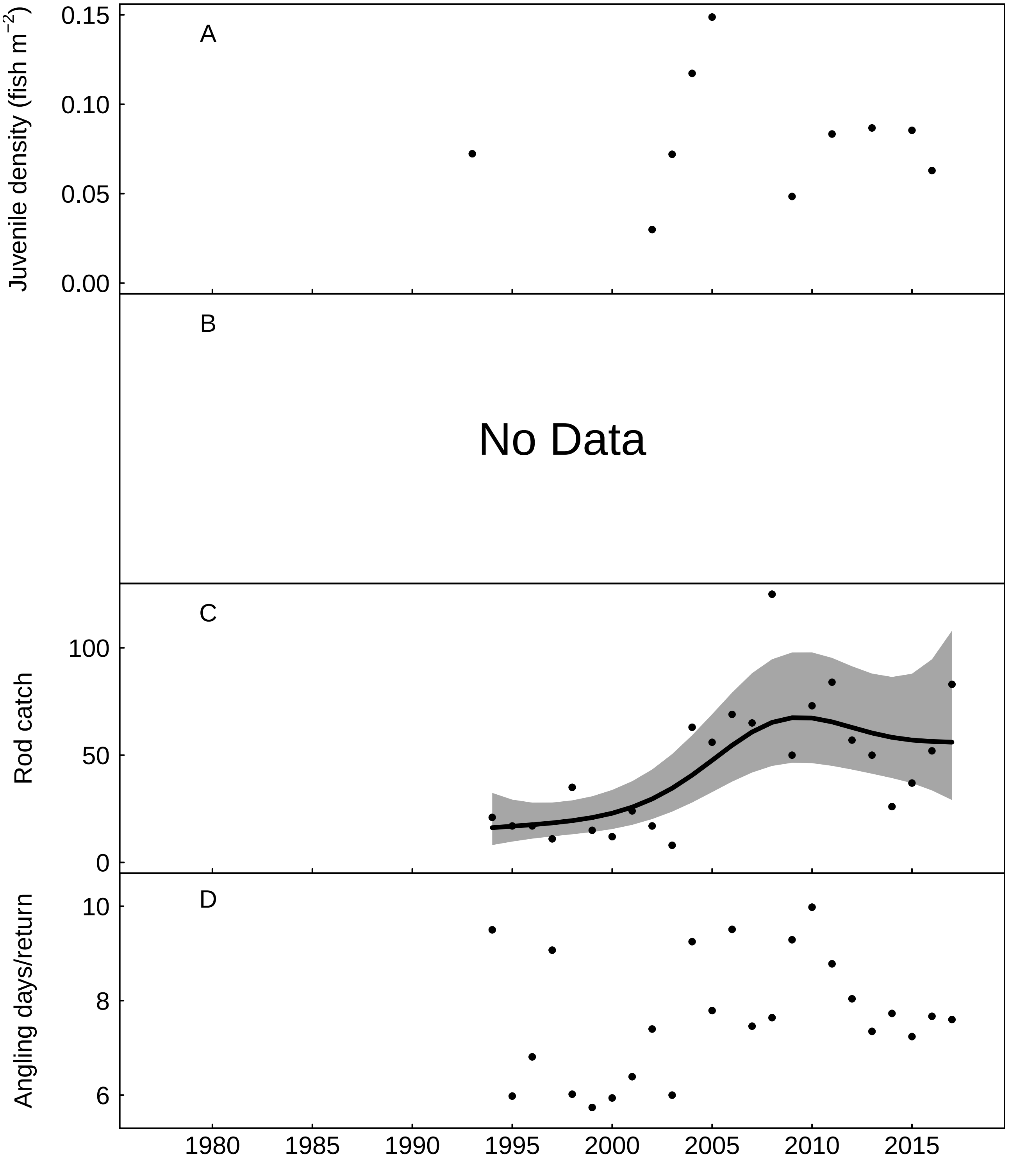


Fig. S73: Trends in the River Duddon for A) juvenile salmon density, B) No Data, C) rod catch, fitted values from negative binomial generalized additive model, grey areas represent 95% point-wise confidence bands for the smoother, and D) angling effort measured as number of days fished per licence return.

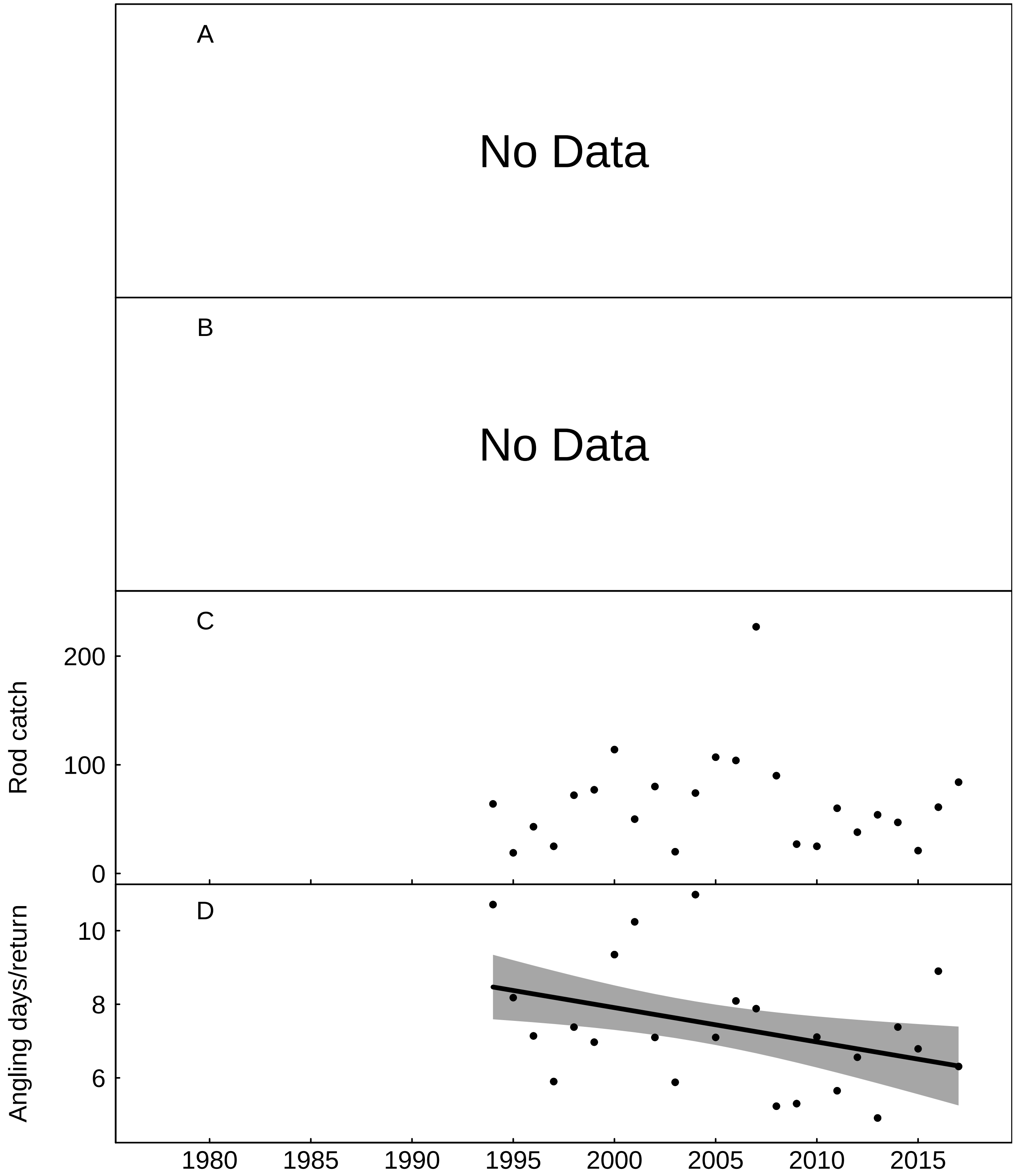

Fig. S74: Trends in the River Esk (Cumbria) for A) No Data, B) No Data, C) rod catch, and D) angling effort measured as number of days fished per licence return, fitted values from Gaussian additive model, grey areas represent 95% point-wise confidence bands for the smoother.

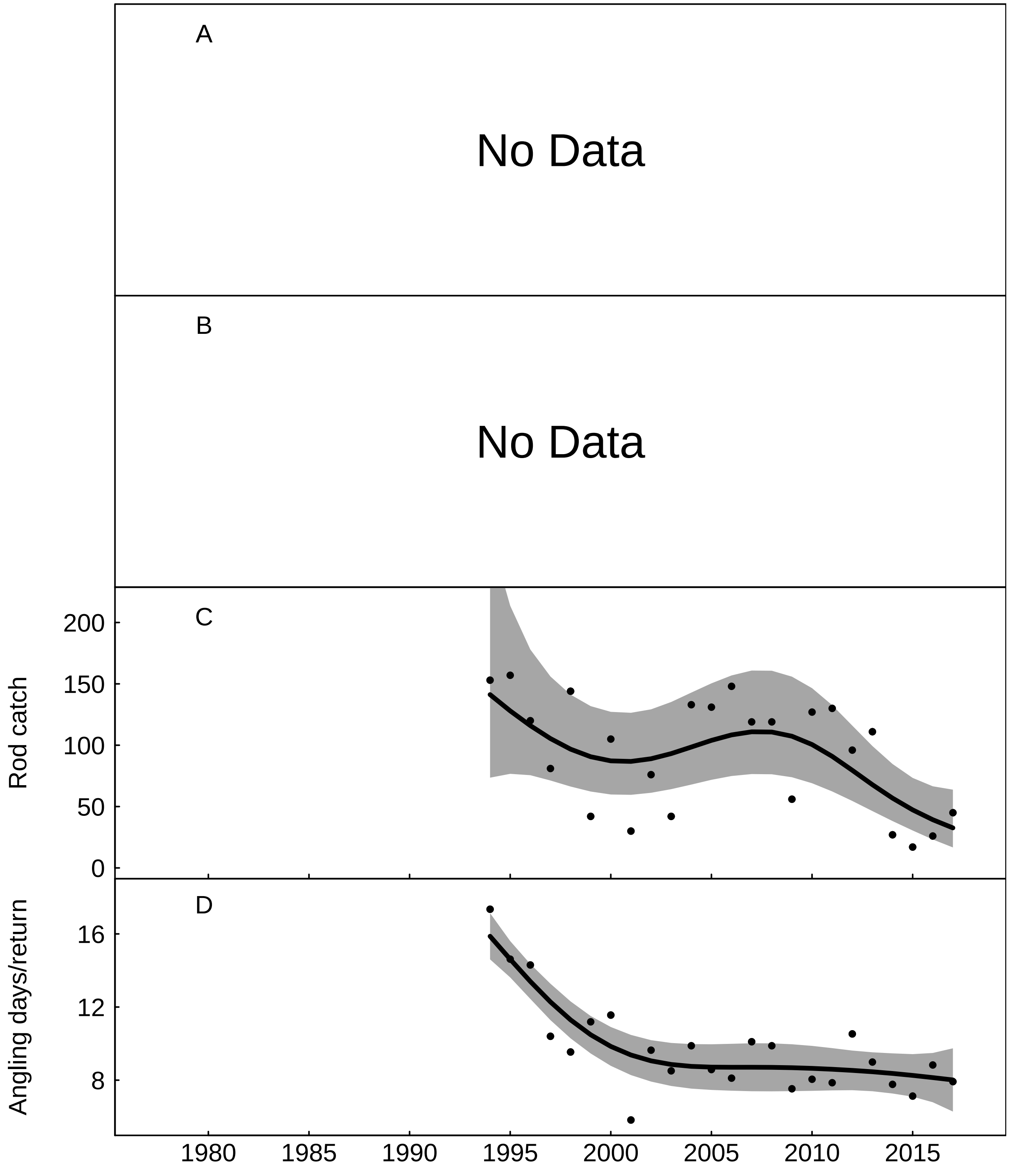


Fig. S75: Trends in the River Irt for A) No Data, B) No Data, C) rod catch, fitted values from negative binomial generalized additive model, grey areas represent 95% point-wise confidence bands for the smoother, and D) angling effort measured as number of days fished per licence return, fitted values from Gaussian additive model, grey areas represent 95% point-wise confidence bands for the smoother.

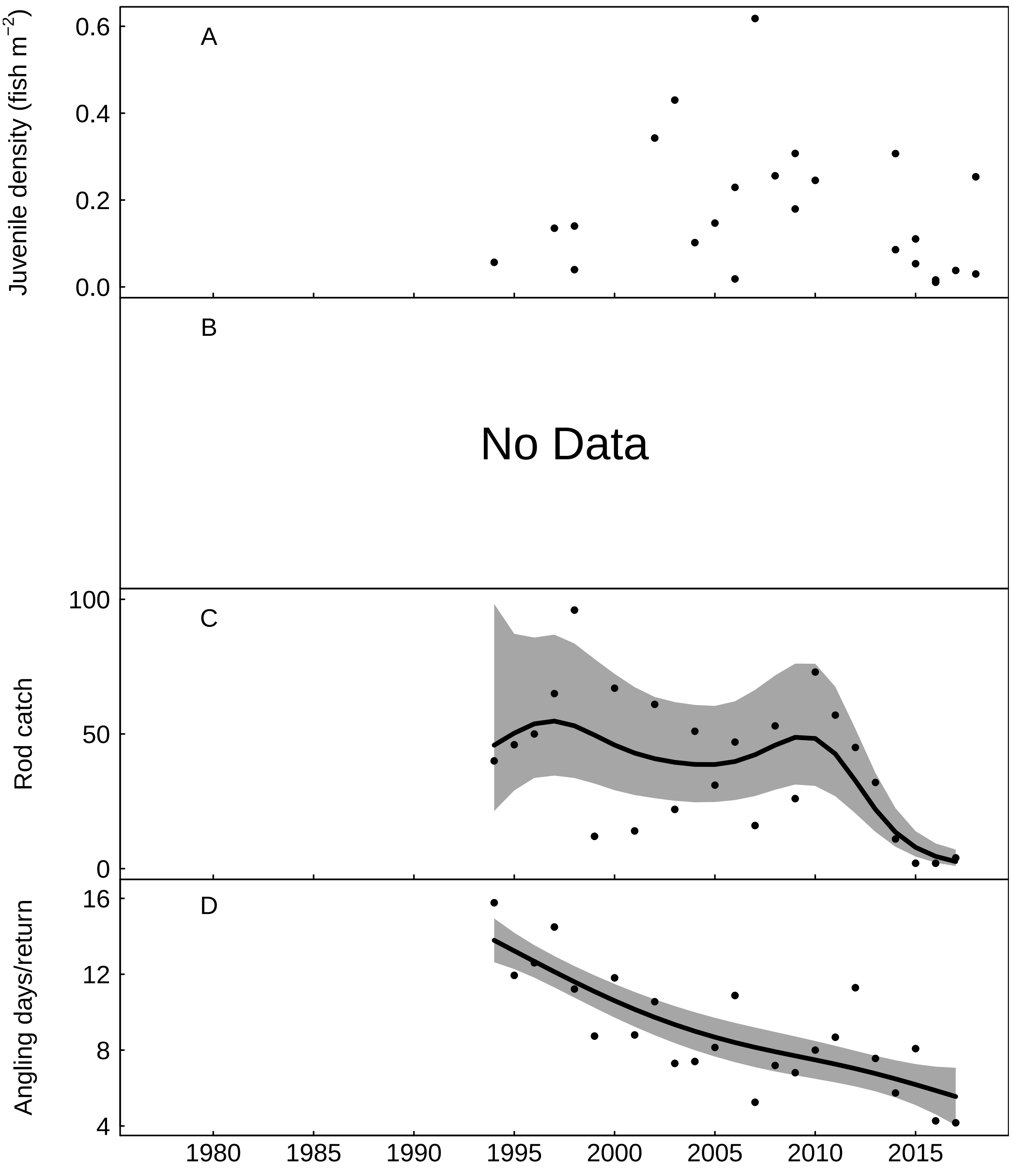


Fig. S76: Trends in the River Calder for A) juvenile salmon density, B) No Data, C) rod catch, fitted values from negative binomial generalized additive model, grey areas represent 95% point-wise confidence bands for the smoother, and D) angling effort measured as number of days fished per licence return, fitted values from Gaussian additive model, grey areas represent 95% point-wise confidence bands for the smoother.

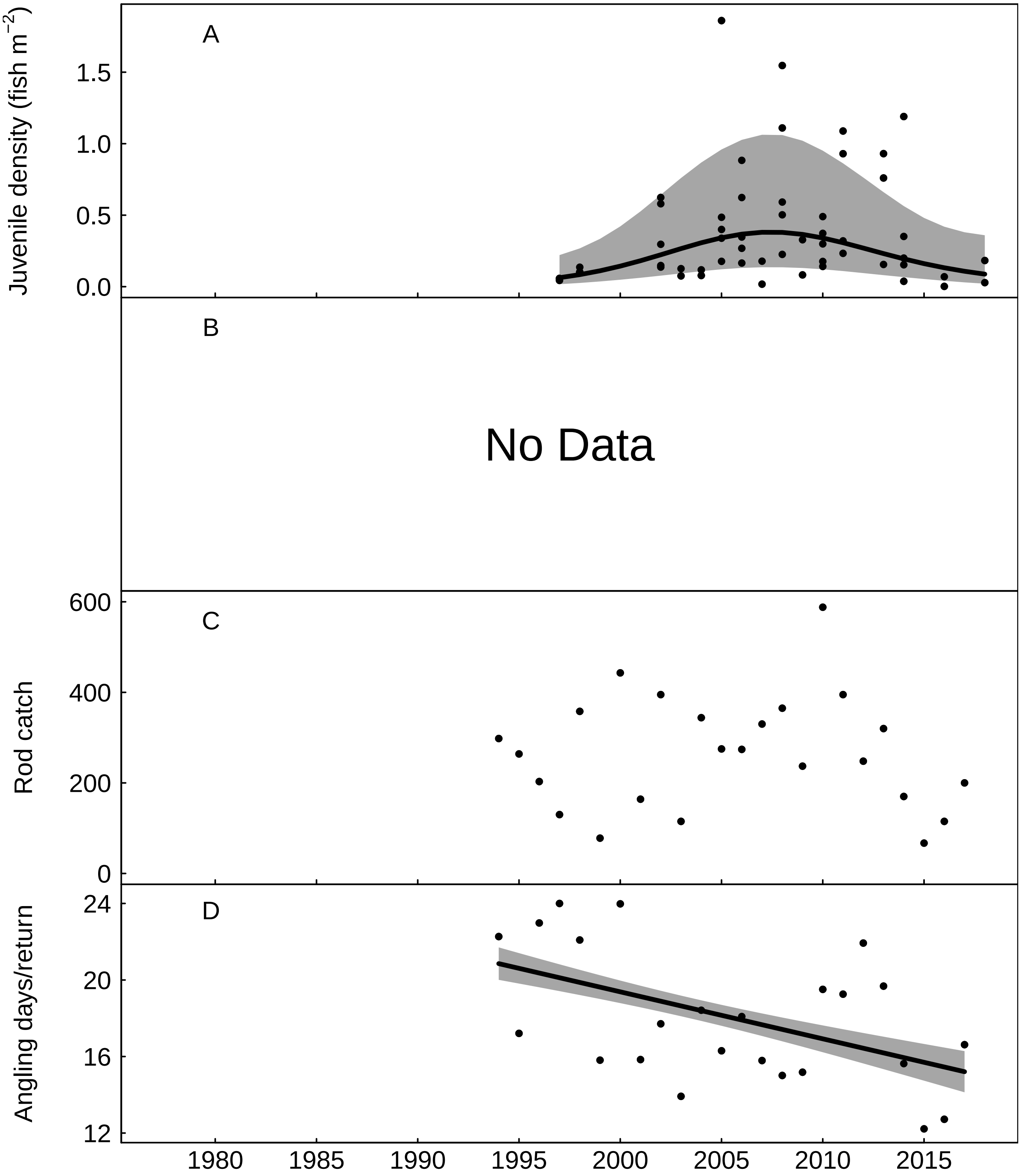


Fig. S77: Trends in the River Ehen for A) juvenile salmon density, fitted values from Gaussian additive mixed model, grey areas represent 95% point-wise confidence bands for the smoother, B) No Data, C) rod catch, and D) angling effort measured as number of days fished per licence return, fitted values from Gaussian additive model, grey areas represent 95% point-wise confidence bands for the smoother.

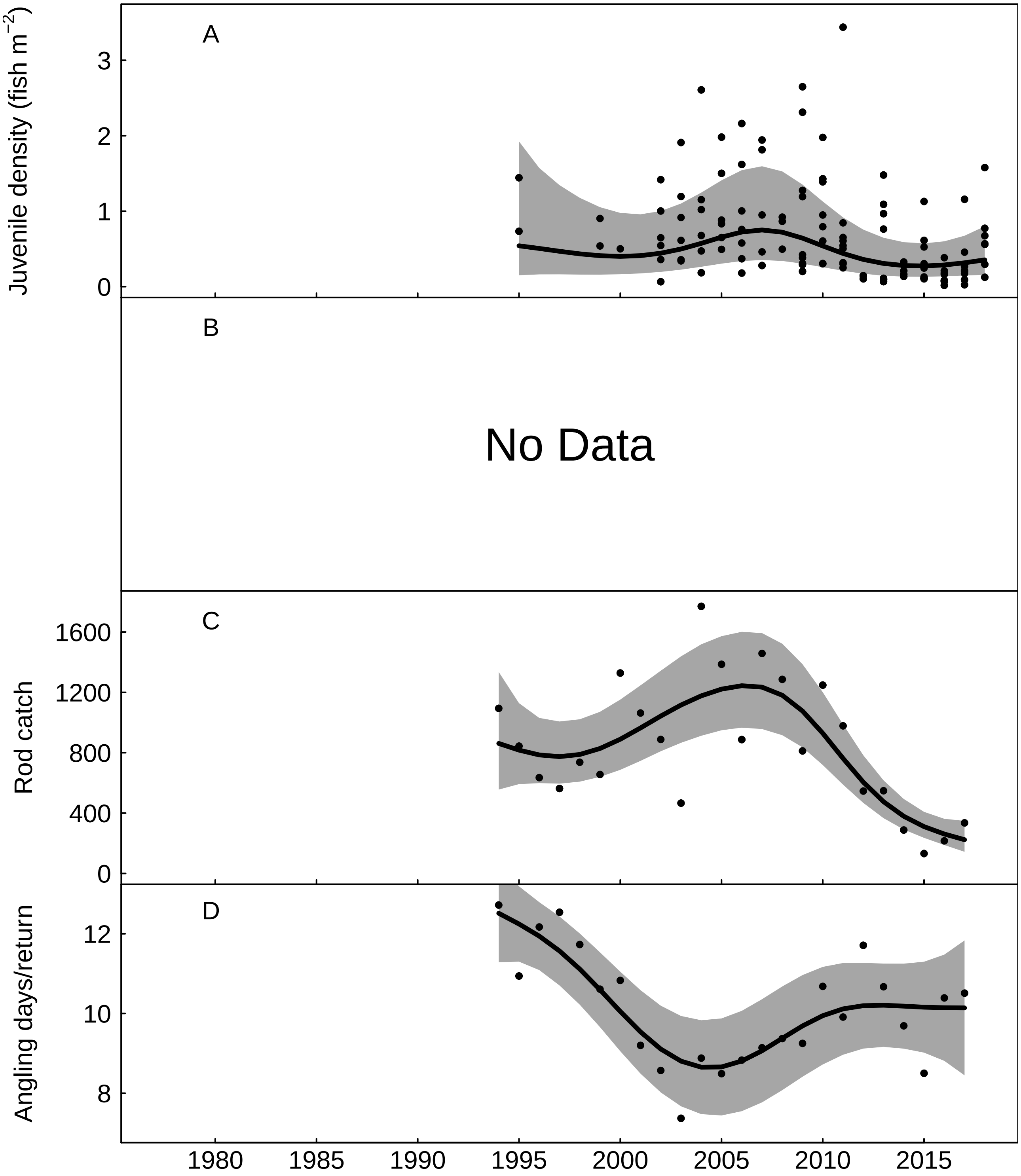


Fig. S78: Trends in the River Derwent (Cumbria) for A) juvenile salmon density, fitted values from Gaussian additive mixed model, grey areas represent 95% point-wise confidence bands for the smoother, B) No Data, C) rod catch, fitted values from negative binomial generalized additive model, grey areas represent 95% point-wise confidence bands for the smoother, and D) angling effort measured as number of days fished per licence return, fitted values from Gaussian additive model, grey areas represent 95% point-wise confidence bands for the smoother.

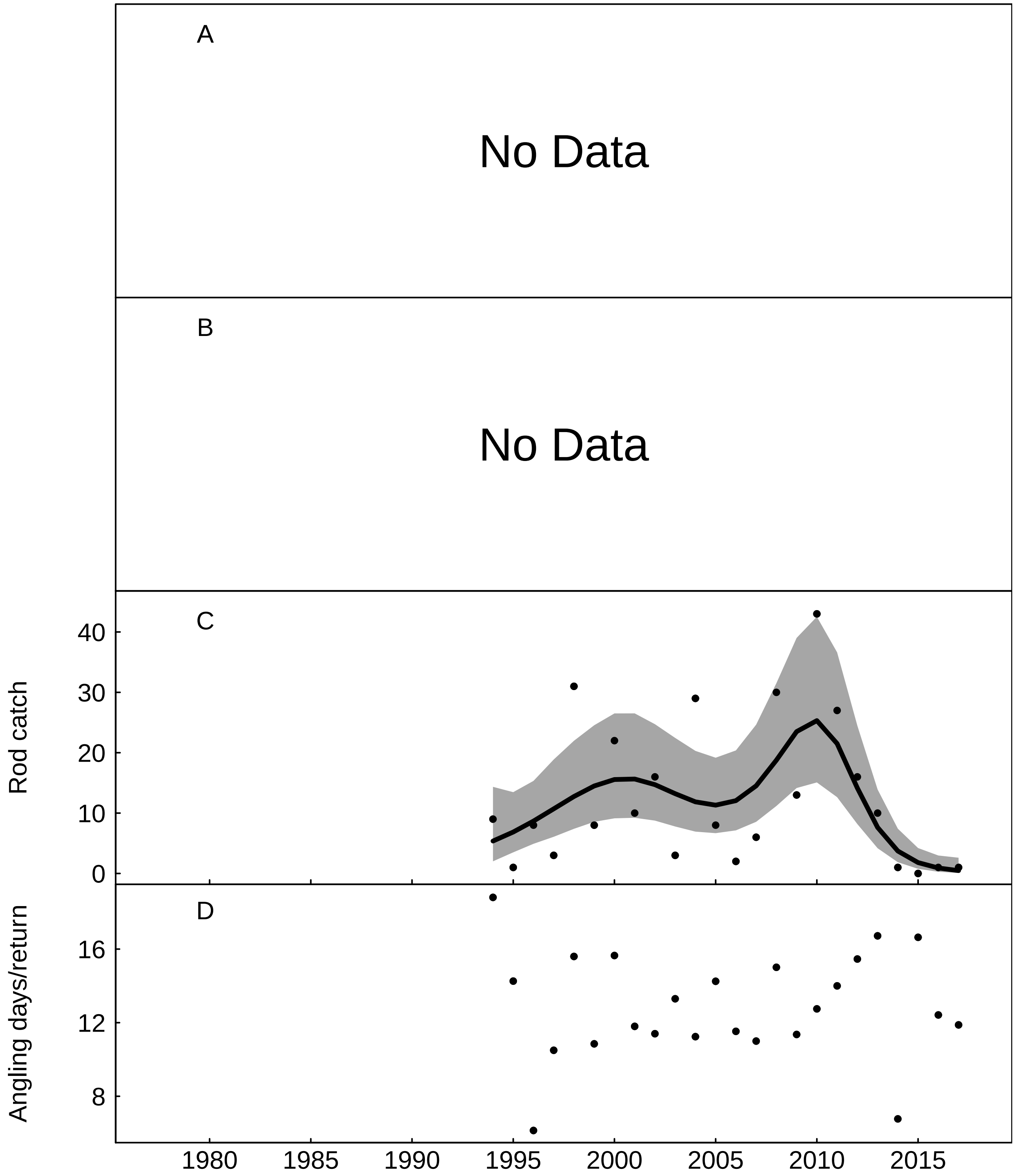


Fig. S79: Trends in the River Ellen for A) No Data, B) No Data, C) rod catch, fitted values from negative binomial generalized additive model, grey areas represent 95% point-wise confidence bands for the smoother, and D) angling effort measured as number of days fished per licence return.

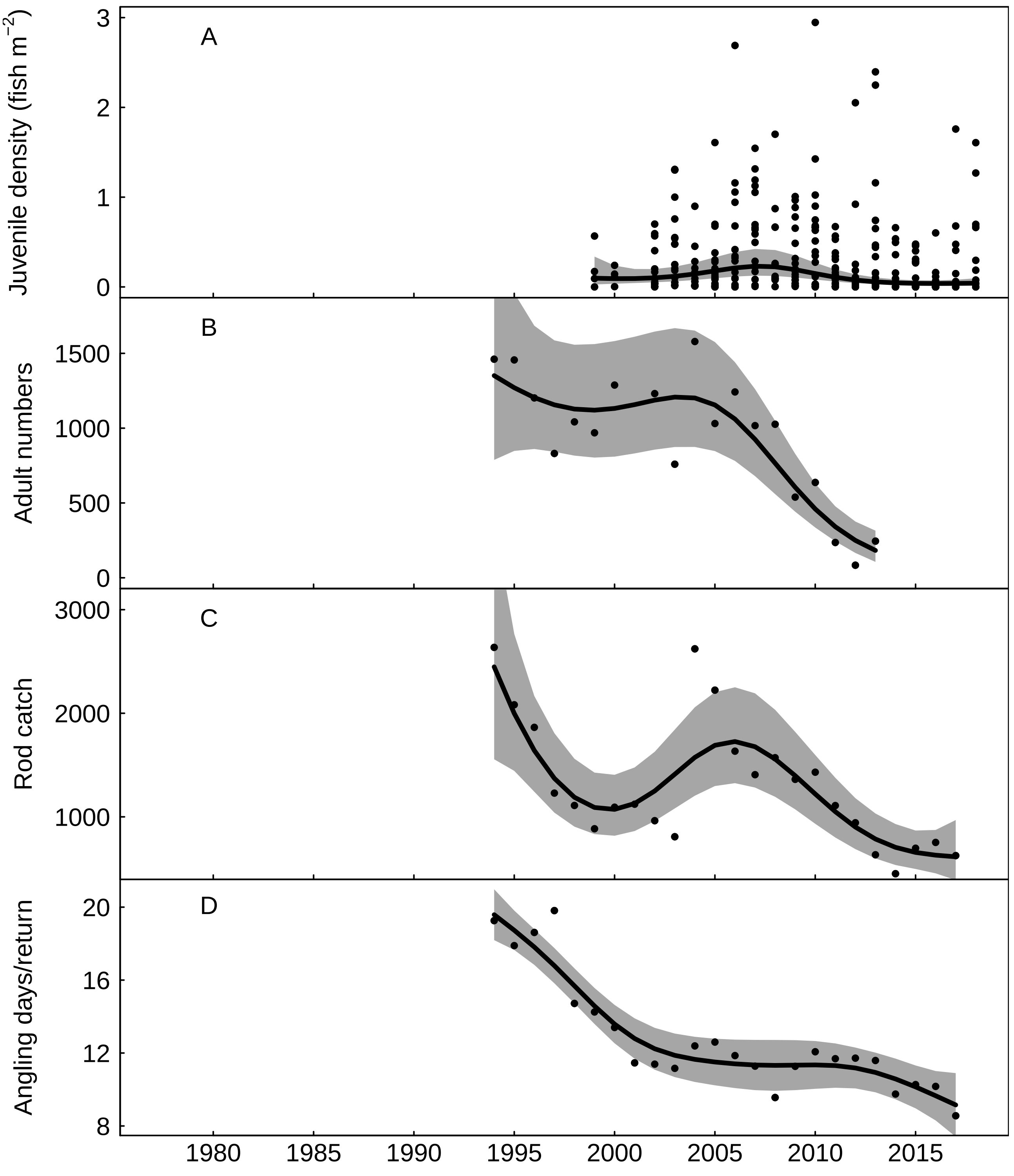


Fig. S80: Trends in the River Eden for A) juvenile salmon density, fitted values from Gaussian additive mixed model, grey areas represent 95% point-wise confidence bands for the smoother, B) returning adult numbers, fitted values from negative binomial generalized additive model, grey areas represent 95% point-wise confidence bands for the smoother, C) rod catch, fitted values from negative binomial generalized additive model, grey areas represent 95% point-wise confidence bands for the smoother, and D) angling effort measured as number of days fished per licence return, fitted values from Gaussian additive model, grey areas represent 95% point-wise confidence bands for the smoother.
